# Genome-wide profiling of druggable active tumor defense mechanisms to enhance cancer immunotherapy

**DOI:** 10.1101/843185

**Authors:** Rigel J. Kishton, Shashank J. Patel, Suman K. Vodnala, Amy E. Decker, Yogin Patel, Madhusudhanan Sukumar, Tori N. Yamamoto, Zhiya Yu, Michelle Ji, Amanda N. Henning, Devikala Gurusamy, Douglas C. Palmer, Winifred Lo, Anna Pasetto, Parisa Malekzadeh, Drew C. Deniger, Kris C. Wood, Neville E. Sanjana, Nicholas P. Restifo

## Abstract

All current highly effective anti-tumor immunotherapeutics depend on the activity of T cells, but tumor cells can escape immune recognition by several mechanisms including loss of function in antigen presentation and inflammatory response genes, expression of immunomodulatory proteins and an immunosuppressive tumor microenvironment. In contrast, the comprehensive identification of strategies that sensitize tumor cells to immunotherapy *in vivo* has remained challenging. Here, we combine a two-cell type (2CT) whole-genome CRISPR-Cas9 screen with dynamic transcriptional analysis (DTA) of tumor upon T cell encounter to identify a set of genes that tumor cells express as an active defense against T cell-mediated killing. We then employed small molecule and biologic screens designed to antagonize gene products employed by tumor cells to actively defend against T cell-mediated tumor destruction and found that the inhibition of BIRC2, ITGAV or DNPEP enhanced tumor cell destruction by T cells. Mechanistically, we found that BIRC2 promoted immunotherapy resistance through inhibiting non-canonical NF-κB signaling and limiting inflammatory chemokine production. These findings show the path forward to improving T cell-mediated tumor destruction in the clinic.

## Introduction

Cancer immunotherapies, which harness the body’s immune system to attack tumor cells, have driven dramatic improvements in clinical outcomes in patients with certain advanced cancers (Maude et al., 2014; Rosenberg et al., 2011; Topalian et al., 2015). Patients with such cancers can experience significant benefit after being treated with immune checkpoint blockade (Topalian et al., 2012) or with adoptive cell therapy (ACT) of tumor infiltrating lymphocytes (TIL), and chimeric antigen receptor (CAR) or T cell receptor (TCR)-engineered T cells (Rosenberg and Restifo, 2015). However, while much progress has been made, many patients with very common cancers such as most cancers of the breast, colon, pancreas, prostate and brain receive limited or no benefit from existing immunotherapy approaches (Park et al., 2018; Yamamoto et al., 2019). Consequently, strategies to improve the efficacy of cancer immunotherapy are needed to improve patient outcomes.

Multiple factors likely play a role in regulating the efficacy of immunotherapy, including the characteristics of anti-tumor T cells (Kishton et al., 2017; Vodnala et al., 2019), tumor microenvironmental factors including the presence of inhibitory factors such as immunosuppressive cell subsets (Facciabene et al., 2012) or excess potassium (Eil et al., 2016), and a number of tumor intrinsic factors. The loss or mutation of tumor cell genes required for antigen presentation and the response to inflammatory cytokines, such as B2M (Restifo et al., 1996) and JAK1/2, can profoundly limit the ability of T cells to eliminate tumors (Patel et al., 2017; Zaretsky et al., 2016). Additionally, the activity of various cellular signaling pathways in tumors, including beta-catenin (Wang et al., 2018) and the PI3K signaling pathway (Sharma et al., 2017), have been demonstrated to provide a survival advantage for tumor cells in the context of immunotherapy. The identification of tumor escape mechanisms may be useful for selecting patients likely to respond to immunotherapy, but have limited utility in the creation of novel strategies for improving therapeutic effects.

A number of approaches have been pursued to optimize the efficacy of cancer immunotherapy, including improving the quality of cell products used in ACT (Gattinoni et al., 2012), altering the tumor microenvironment, and pursuing strategies to combine various pharmacological inhibitors and biologics with immunotherapy treatments as a combination therapy (Robert et al., 2016). Recent efforts to investigate the potential of combining targeted pharmacological inhibitors with immunotherapy have demonstrated potential to improve clinical efficacy (Ribas et al., 2019), indicating such approaches may play an important role in improving the responses of patients to immunotherapy. We sought to systematically sift through strategies and solutions to identify methods for enhancing cell-based immunotherapy on a whole genome scale.

We hypothesized that, as often observed in the context of cancer cells treated with targeted chemotherapies (Singleton et al., 2017), the engagement of anti-tumor T cells with target tumor cells might drive tumor transcription of genes that promote cell survival, mediating resistance to immunotherapy. These genes, comprising a tumor active defense mechanism against immunotherapy, would represent important potential targets for combination therapy. We sought to identify the genetic components of active tumor defense against immunotherapy by combining a genome-scale CRISPR/Cas9 loss of function screen in tumor cells with dynamic transcriptional analysis (DTA) of tumor cells as they interact with antigen-specific T cells. The integration of these cross-platform, multi-omics approaches allowed us to identify a set of tumor genes, including *BCL2* and *BIRC2,* whose expression was both increased during T cell engagement and required for tumor cell survival in this context. We evaluated the potential for targeting these tumor genes, along with other top hits from the CRISPR screen to improve the efficacy of immunotherapy and uncovered several small molecules and biologics that augmented T cell elimination of tumor cells. Finally, we explored the genetic mechanism through which *BIRC2* acts to limit tumor cell susceptibility to T cell-mediated killing, finding a novel role for *BIRC2* in suppressing tumor cell inflammatory signaling and chemokine production by inhibiting the non-canonical NF-κB signaling pathway. Taken together, these results demonstrate a cross-platform, multi-omics strategy to identify and evaluate clinically relevant tumor targets for use in combinatorial immunotherapy treatment regimens.

## Results

### A two-cell type (2CT) genome-scale CRIPSR/Cas9 screen optimized to identify tumor genes that promote survival in the context of T cell engagement

To identify active tumor defense genes, we first sought to define tumor genes that are essential for tumor cell survival in the context of tumor antigen-specific T cell engagement. To achieve this, we utilized a two-cell type (2CT) co-culture system in which primary human T cells are transduced with an HLA-A*0201-restricted T cell receptor (TCR) specific for the NY-ESO-1 antigen (NY-ESO-1:157-165 epitope), then co-cultured with NY-ESO-1^+^ Mel624 human melanoma tumor cells (Patel et al., 2017). Our previous study (Patel et al., 2017), in which we sought to identify tumor genes that are essential for effective immunotherapy, utilized an experimental design in which transduced T cells were cultured with Mel624 melanoma cells at a 0.3 ratio (1:3 E:T) for 12 hours, resulting in a strong T cell selective pressure that eliminated approximately 75% of tumor cells. This experimental design allowed for the identification of tumor genes whose loss promoted cell survival. The goal of our current study was diametrically opposite: Rather than seeking genes involved in tumor escape, we sought to identify genes whose loss *promoted* tumor cell death in the context of T cell engagement. To optimize the 2CT assay for this setting, we reduced the T cell selective pressure by shortening co-culture duration to 6 hours, resulting in the elimination of approximately 25% of tumor cells at a 1:3 E:T ratio (**Figure 1A**). With these reduced T cell selective pressure conditions established, we transduced Mel624 cells with the GeCKOv2 library (**Figure 1B**) and exposed the transduced tumor cells to ESO T cells at a 1:3 E:T ratio for 6 h. We utilized deep sequencing to profile sgRNA library representation in tumor cells before and after T cell co-culture and, reasoning that sgRNAs targeting genes that promote tumor cell survival would have reduced abundance in surviving tumor cells, ranked all genes by two different scoring metrics: second-most depleted sgRNA (**Figure 1C**) and the RNAi Gene Enrichment Ranking (RIGER) metric (**Figure 1D**). Top hits identified from the screen included genes such as *TRAF2, BIRC2, ALG11* and *mir1299.* Gene ontology analysis of significantly depleted genes indicated pathway enrichments of death receptor signaling, EIF2 signaling, endoplasmic reticulum stress pathways, and the unfolded protein response (**Figure 1E**). Demonstrating the reproducibility of this screening method, we found that top genes identified in the 2CT screen including *mir1299, FAM32A, EIF3I, BIRC2* and *BCL2* were also depleted in a replicate screen (**Figure 1F, Table 1)**.

**Figure 1.**
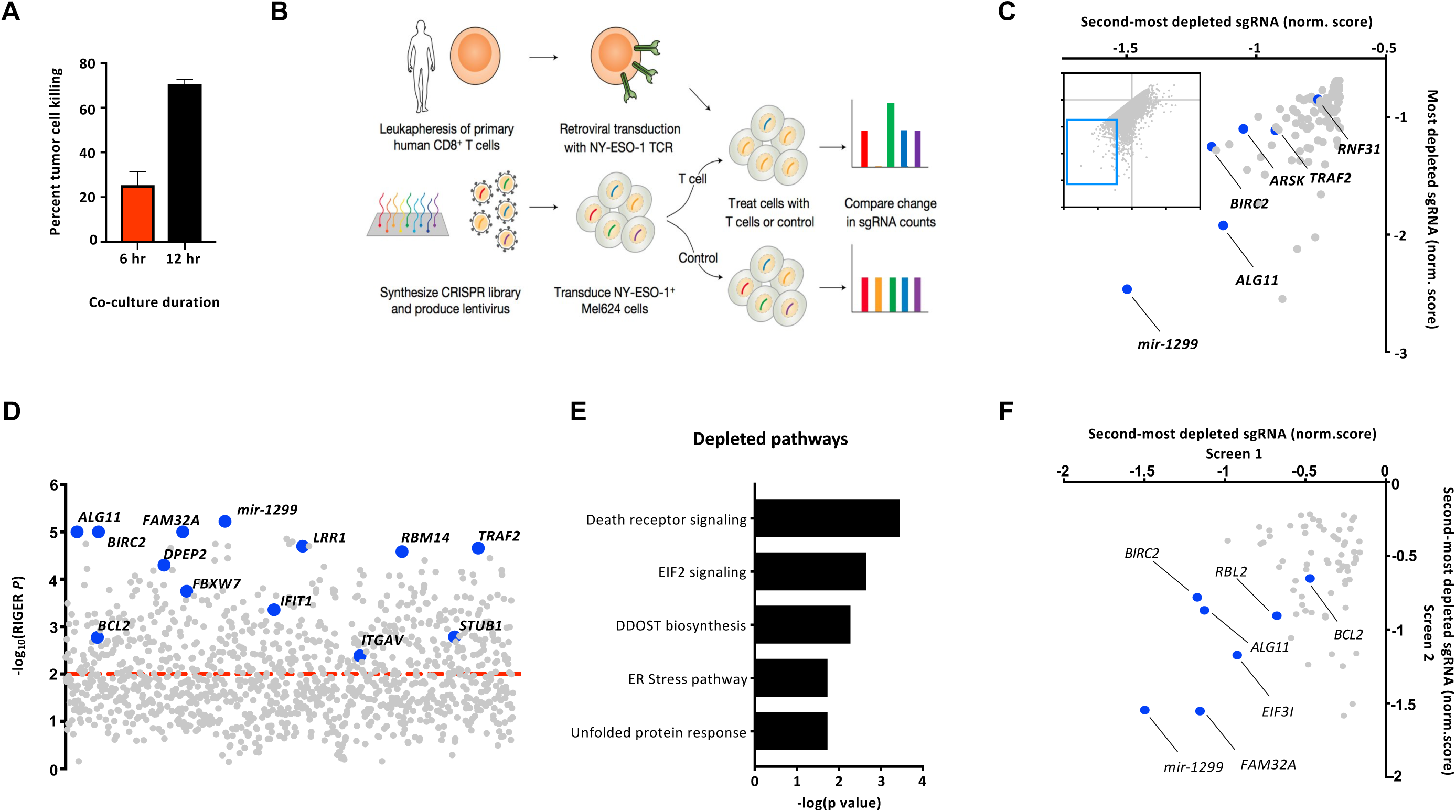
A genome-scale 2CT assay optimized for identifying tumor genes promoting resistance to T cell-mediated death. **(A)** Mel624 melanoma cells were co-cultured with ESO T cells at a 1:3 E:T ratio for 6 or 12 hr and tumor cell survival was measured by flow cytometry. **(B)** Experimental design of genome-scale CRISPR/Cas9 2CT screen in which Mel624 cells were transduced with the GECKOv2 library and cultured with ESO T cells for 6 hr. sgRNA representation was measured in control and ESO T cell treated tumor cells. Depletion of sgRNA representation in ESO T cell treated tumor cells was compared with control treated cells and depicted as **(C)** a scatterplot of the normalized enrichment of the most-depleted sgRNA vs. the second-most depleted sgRNA for all genes after ESO T cell treatment (top 100 most depleted genes depicted in enlarged region) or **(D)** RIGER analysis was performed to identify top depleted genes. **(E)** Top 250 genes for which sgRNAs were significantly (*P* < 0.05) depleted by RIGER ranking were analyzed by Ingenuity Pathway Analysis (IPA) and enriched pathways were determined. **(F)** Second-most depleted sgRNAs for genes treated with ESO T cells were compared in biological replicate screens. Data is representative of two biological replicate experiments with four technical replicates in each condition **(A)** or depicts data from one **(C-E)** or two **(F)** genome-scale CRISPR screens.

### T cell engagement drives tumor cell transcriptional activation of pro-survival pathways

We next utilized dynamic transcriptional analysis (DTA) of the 2CT system to identify tumor genes that are differentially expressed upon T cell engagement with tumor cells. We co-cultured ESO T cells with unmodified Mel624 cells for 0 or 6 h (**Figure 2A**). ESO T cells and tumor cells were each purified by FACS sorting **(Figure S1A)** and mRNA was extracted and sequenced. As expected, upon encounter with NY-ESO-1 expressing tumor cells, ESO T cells increased the expression of inflammatory cytokines such as *IL2* and *IFNG*, cytolytic molecules including *GZMB*, markers of T cell activation *CD69* and *TNFRSF9* (encoding 4-1BB), transcription factors promoting T cell effector function including *ZEB2,* and pathways supporting anabolic growth. Conversely, tumor-experienced T cells acutely repressed the expression of genes associated with memory and stemness such as *KLF2, TCF7, CD27,* and *LEF1* (**Figure 2B, Figure S1B)**. After co-culture with ESO T cells, Mel624 tumor cells upregulated the expression of genes associated with cellular mediators of inflammation including TNFR and NF-κB pathway signaling. Tumor cells also upregulated the expression of genes that have been associated with resistance to cell death such as *BIRC2, BIRC3, BCL2A1* (**Figure 2C-D**). We also observed that tumor cells downregulated the expression of genes associated with Notch pathway signaling after T cell engagement (**Figure 2D**). Previous reports have linked the NF-kB signaling pathway with resistance to immunotherapy (Pan et al., 2018). Consistent with this, we found that T cell-experienced tumor cells upregulated a set of NF-κB regulated genes associated with resistance to apoptosis (Dutta et al., 2006), while repressing the expression of the apoptotic effector molecule *BAX* (**Figure 2E**).

**Figure 2.**
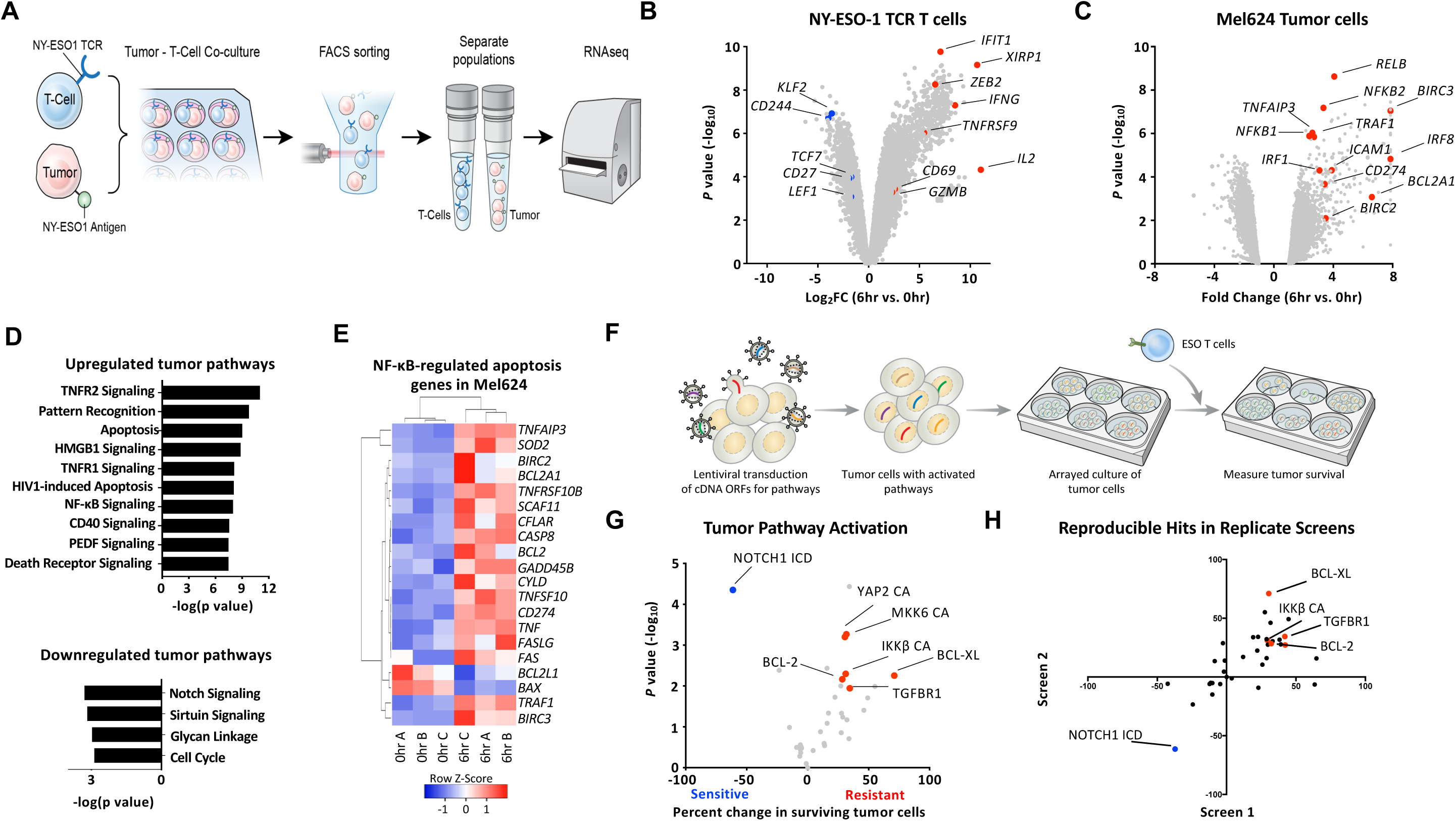
Dynamic transcriptional profiling of 2CT interaction reveals tumor transcriptional response to T cell engagement. **(A)** Experimental design of transcriptional profiling of dynamics of ESO T cell interaction with Mel624 tumor cells. Cells were co-cultured at a 1:3 E:T ratio for 0 or 6 hr and each cell population was sorted by FACS. mRNA expression levels were quantified by RNA sequencing. **(B)** Volcano plot representing differentially expressed genes analyzed by RNA-seq of CD3+mTCRb+ T cells that were co-cultured with Mel624 cells for 0 or 6 hr. Abundance is represented as relative fold change (*x* axis) versus significance (*y* axis). **(C)** Volcano plot representing differentially expressed genes analyzed by RNA-seq of Mel624 cells that were co-cultured with ESO T cells for 0 or 6 hr. Abundance is represented as relative fold change (*x* axis) versus significance (*y* axis). **(D)** Ingenuity Pathway Analysis (IPA) of Mel624 genes that were significantly upregulated or downregulated upon ESO T cell co-culture. **(E)** Heatmap of NF-κB regulated apoptosis gene expression in Mel624 cells following a 0 or 6 hr co-culture with ESO T cells. **(F)** Experimental design of tumor pathway activation assay. A375 melanoma cells were transduced via lentivirus with cDNA ORFs encoding cellular signaling pathway activating constructs. Transduced cells were cultured with ESO T cells in an arrayed fashion, and the impact of pathway activation on susceptibility to T cell-mediated killing was assessed. **(G)** Percent change in tumor cell death (compared with control construct) for each pathway-activated cell line. Relative percent change in tumor cell death is plotted (*x* axis) versus significance (*y* axis). **(H)** Percent change in tumor cell death in replicate pathway activation screens. Data is pooled from three independent experiments **(B-E)**, is representative of 2 independent experiments with four technical replicates per group **(G)** or depicts the results of 2 independent experiments **(H)**.

To determine whether the transcriptional and signaling pathway alterations we observed in tumor cells following T cell interaction had functional consequences in regulating tumor cell susceptibility to immunotherapy, we systematically engaged 17 tumor cellular signaling pathways through lentiviral transduction of pathway activating constructs (Martz et al., 2014). Pathway-activated tumor cells were co-cultured with ESO T cells in an arrayed fashion, and the effects of activating each cell signaling pathway on tumor cell survival were measured (**Figure 2F**). Consistent with previous reports showing NF-kB signaling can promote tumor evasion of immune-mediated destruction (Pan et al., 2018), and with our data indicating upregulation of NF-κB regulated pro-survival genes in tumor cells after ESO T cell engagement, activation of NF-κB signaling or constitutive expression of *BCL2* or *BCL-XL* resulted in tumor cell resistance to T cell-mediated destruction (**Figure 2G, 2H**). Conversely, constitutive activation of the NOTCH1 signaling cascade sensitized tumor cells to T cell-mediated death (**Figure 2G, 2H**).

### Tumor cells actively express defense mechanisms to avert T cell-mediated destruction

Tumor cell genes that are induced by T cell-encounter and functionally promote tumor cell survival may represent important tumor-intrinsic mechanisms that actively promote immunotherapy resistance. To determine whether such genes could be identified, we focused on genes whose targeting sgRNAs were depleted in both biological replicate CRISPR screens and assessed mRNA expression changes in these tumor genes following T cell encounter. We found 10 CRISPR-depleted genes that had statistically significant increases in mRNA expression after T cell engagement **(Table 2)**, including *SCAF11*, *BIRC2*, *RPAP2*, and *BCL2* (**Figure 3A**). To validate the functional role of these genes in mediating tumor cell resistance to T cell-induced apoptosis, we transduced A375 melanoma cells, an additional human melanoma that expresses both HLA-A*0201 and NY-ESO-1 antigen, with Cas9 and three newly-designed sgRNA constructs targeting each candidate gene. We co-cultured these cells with ESO T cells and assessed the impact of gene knockout on tumor cell survival. We considered a candidate gene to be validated if at least two independent gene-targeted sgRNAs resulted in increased tumor cell death upon T cell co-culture. Five of the candidate genes met these criteria, including *BCL2*, *BIRC2*, *LPGAT1*, *SCAF11* and *TOR1AIP1* (**Figure 3B**). These genes represent strong candidates of actively-induced tumor cell-intrinsic mechanisms of resistance to T cell-mediated killing.

**Figure 3.**
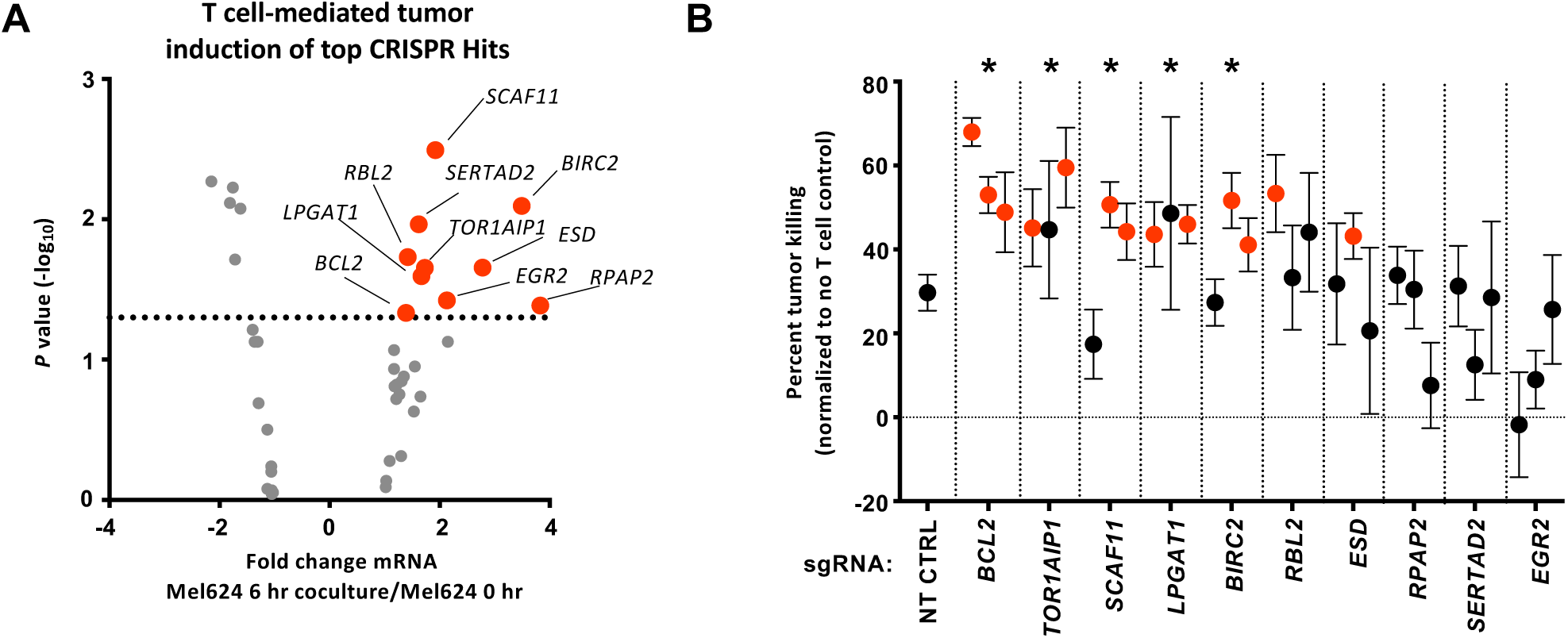
Identification of T cell-induced tumor-intrinsic genes which act to promote survival. **(A)** Volcano plot representing differentially expressed tumor genes that were identified as being significantly depleted from duplicate CRISPR 2CT screens by RIGER analysis (n=42 genes). Expression of genes was analyzed by RNA-seq as in Figure 2 with abundance represented as relative fold change (*x* axis) versus significance (*y* axis). Ten genes which met CRISPR screen depletion criteria were significantly induced by T cell engagement (red dots; *P* value < 0.05, fold change > 1.3. **(B)** The effect of gene knockout on tumor susceptibility to killing by T cells was measured by flow cytometry. Statistically significant sgRNAs for individual genes are indicated by red dots, as they resulted in a significant increase in tumor killing by T cells relative to non-targeting sgRNA (*p* < 0.05). A gene was considered validated (* at top) if two or more sgRNAs were statistically significant. Data is pooled from three independent experiments **(A)** or is representative of three independent experiments **(B)**. *P* values calculated for negatively enriched gene-targeting sgRNAs compared to control sgRNA by two tailed Student’s *t* test.

### CRISPR-informed drug screen identifies combinatorial immunotherapy targets

To assess the potential utility of targeting active tumor defense mechanisms for combination therapy, we developed a workflow for identifying potential targets for small molecule or biologic inhibition based on these datasets. As an initial step, we considered the five identified and validated tumor active defense genes (**Figure 4A**) along with the top 250 second-most depleted genes from the duplicate CRISPR screens ordered by rank-sum analysis. Genes were first mapped to coded proteins **(Table 3)**, and these proteins were queried for availability of targeted inhibitors using the Drug Gene Interaction Database (DGIDB) (Griffith et al., 2013) **(Table 4)** followed by manual curation. We identified 15 targets for which inhibitors were available, including BIRC2, BCL2, ITGAV and DNPEP **(Table 5)**. When possible, we obtained two independent inhibitors per target to limit drug-specific effects. To assess the impact of inhibitors on T cell elimination of tumor cells, we first performed a high throughput assay in which ESO T cells were co-cultured with A375 melanoma cells in the presence of vehicle or each inhibitor for 16h (**Figure 4A**). Assays were performed at two concentrations of each inhibitor and tumor cell viability was measured by colorimetric viability dye. We found that a number of inhibitors were capable of increasing A375 killing compared with ESO T cells alone (**Figure 4B, 4D**). We reasoned that inhibitors of interest should both significantly increase tumor cell destruction in combination with ESO T cells when compared with ESO T cells alone and should also cause higher tumor cell killing as a combination with ESO T cells compared with inhibitor treatment alone. Our screen revealed several inhibitors that met these criteria, including inhibitors of BIRC2 (birinapant and LCL161), ITGAV (Cilengitide and a monoclonal antibody against ITGAV), DNPEP (CHR2797), BCL2 (Abt199), and ERRα (C29) (**Figure 4C, 4E**).

**Figure 4.**
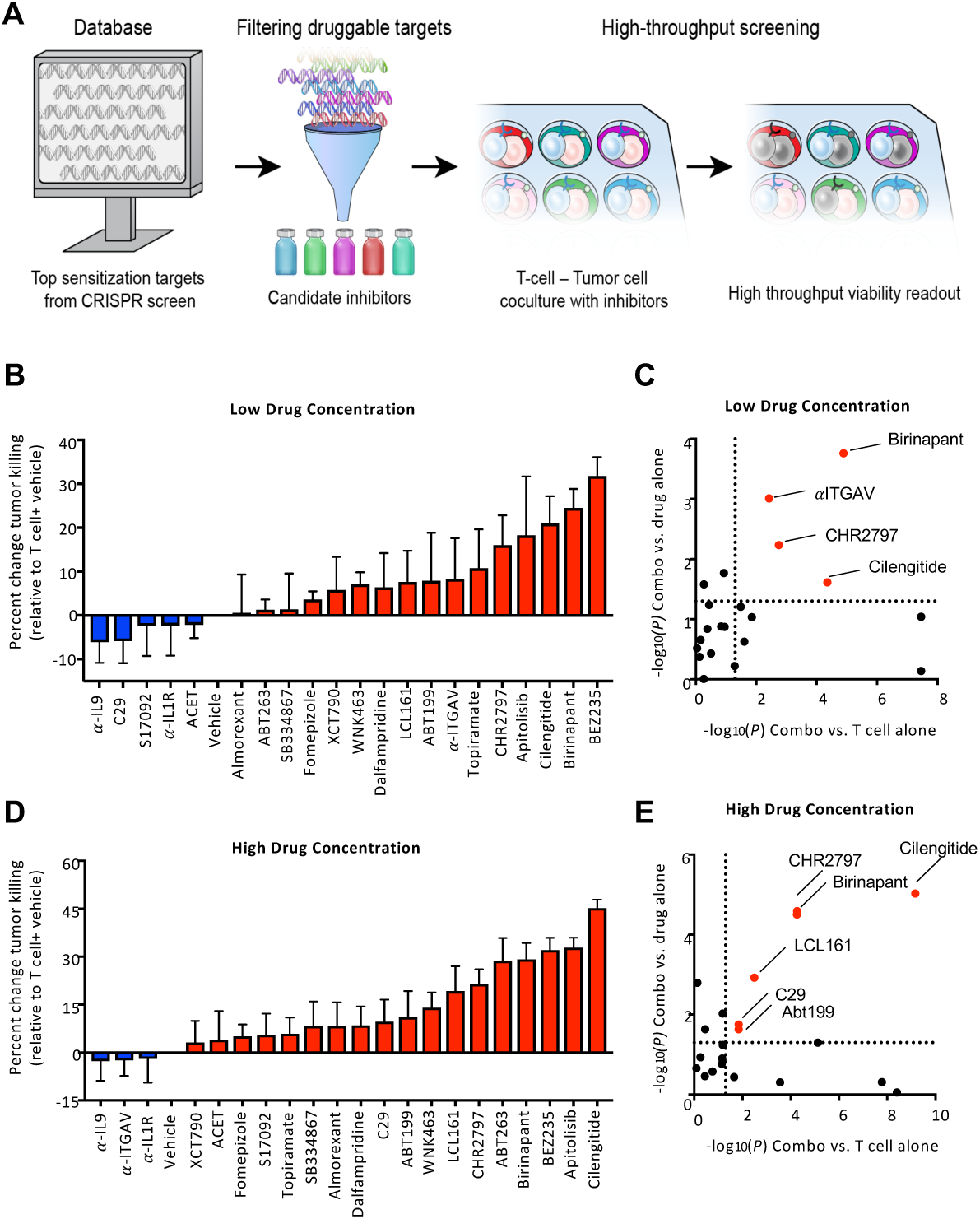
Inhibitors of CRISPR screen-identified genes sensitizes tumor cells to T cell-mediated killing. **(A)** Workflow utilized to identify and screen inhibitors for capacity to increase T cell-mediated destruction of tumor cells. Top 250 hits from CRISPR screen were filtered to identify targets with available inhibitors. A375 melanoma cells were co-cultured with ESO T cells at a 1:2 E:T ratio for 16hr in the presence or absence of inhibitors. Co-cultures washed and tumor cell viability was measured with WST1 viability reagent. The impact of inhibitors on ESO T cell killing of A375 cells was measured at low (500 nM except for α-ITGAV at 250 ng/mL and Fomepizole and Dalfampridine at 500 μM) and high (5 μM except for α-ITGAV at 2.5 μg/mL and Fomepizole and Dalfampridine at 5 mM) concentrations **(B, D)**. **(C,E)** Plot of *p* values (significance threshold *P* < 0.05, dotted line) between tumor elimination by the combination inhibitor and ESO T cell treatment versus ESO T cells alone (*x* axis) and tumor elimination by the combination treatment versus inhibitor alone (*y* axis). Data is representative of three independent experiments with four technical replicates per sample **(B-E)**. *P* values calculated by two tailed Student’s *t* test.

### Combination treatment increases T cell elimination of melanoma and GI tumor cells

We next tested the effects of these inhibitors in additional 2CT assays in which we assessed the impact of combination therapy on the ability of T cells to eliminate tumors across several tumor antigens and cancer types. To increase the precision of our readouts, we co-cultured TCR transduced T cells with tumor lines expressing target antigens for 16h in the presence or absence of drugs and tumor cell viability was measured at a single cell level using FACS (**Figure 5A**). Consistent with our initial findings, we found that inhibitors of BIRC2 (birinapant and LCL161), ITGAV (cilengitide and anti-ITGAV) and DNPEP (CHR2797) increased the killing of A375 melanoma cells after co-culturing ESO T cells (**Figure 5B, S2A)**. However, we did not observe increased tumor cell elimination by ESO T cells when BCL2 (Abt199) or ERRα (C29) were inhibited in this assay, suggesting that the observed impacts on cell viability in the high throughput assay could reflect toxicity against anti-tumor T cells. Increases in tumor cell death in the presence of inhibitors was dependent on the presence of T cells for all validated hits (**Figure S2B)**. Importantly, we also found that, except for α-ITGAV, these drugs increased the killing of a second melanoma line (Mel624 expressing both NY-ESO-1 and MART-1 antigens) after co-culture with both ESO T cells (**Figure 5C**) and T cells transduced with a TCR that confers recognition of MART-1 (**Figure 5D**).

**Figure 5.**
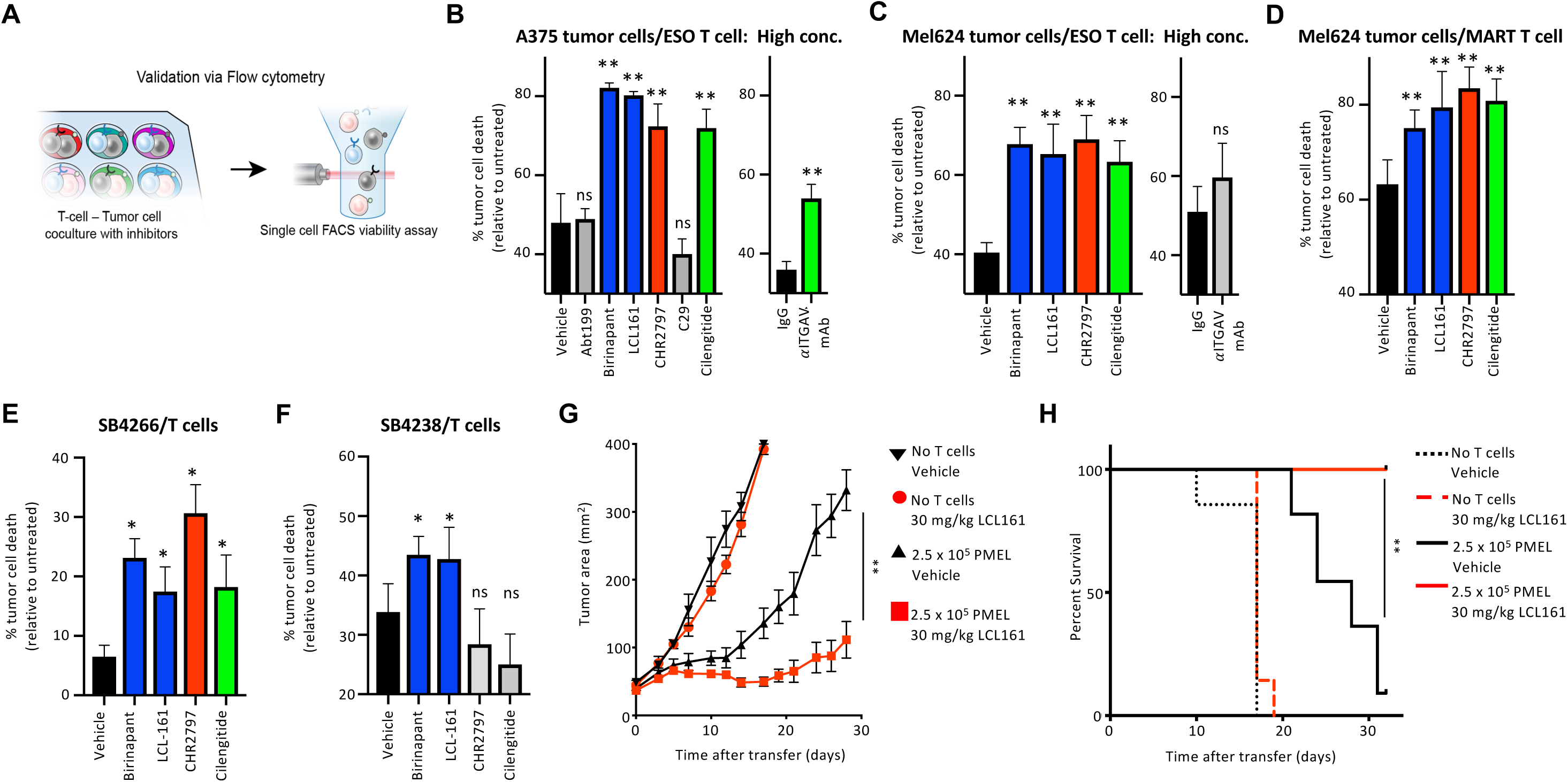
Inhibitors sensitize tumor cells to T cell-mediated killing irrespective of antigen target. **(A)** Depiction of experimental design. Tumor cells were co-cultured with T cells at an E:T ratio of 1:2 for 16 hr in the presence or absence of inhibitors and tumor cell viability was measured by flow cytometry. **(B)** A375 melanoma cells were co-cultured with ESO T cells along with inhibitors at 5μM except for α-ITGAV at 2.5 μg/mL, and tumor cell elimination was measured by flow cytometry. **(C)** Mel624 melanoma cells were co-cultured with ESO T cells along with inhibitors at 5μM except for α-ITGAV at 2.5 μg/mL, and tumor cell elimination was measured by flow cytometry. **(D)** Mel624 melanoma cells were co-cultured with MART-1 T cells along with inhibitors at 5μM except for α-ITGAV at 2.5 μg/mL, and tumor cell elimination was measured by flow cytometry. **(E)** Colon cancer cell line SB4266 was co-cultured with T cells transduced with a TCR reactive against p53 R248W mutation along with inhibitors at 5μM. Tumor cell death was measured by flow cytometry. **(F)** Colon cancer cell line SB4238 was co-cultured with T cells transduced with a TCR reactive against NCKAP1 D438Y along with inhibitors at 5μM. Tumor cell elimination was measured by flow cytometry. **(G, H)** Subcutaneous tumor growth in mice receiving ACT of PMEL1 T cells along with vehicle or LCL161 (30 mg drug/kg body weight). Mice were treated with LCL161 via IP injection every 48hr beginning 24hr after PMEL1 T cell infusion for a total of 5 doses. Tumor area **(G)** and mouse survival **(H)** are shown. Data are representative of four **(B-C)**, three **(D, G-H)** or two **(E-F)** independent experiments with four technical replicates per groups **(B-F)** or 10 mice per group **(G-H)**. * *P* < 0.05, ** *P* < 0.01. *P* values for *in vitro* assays **(B-F)** calculated by one-way ANOVA with multiple comparisons corrected with Dunnett adjustment. *P* values for *in vivo* assays **(G-H)** calculated by Wilcoxon rank sum test **(G)** and or log-rank test **(H)**. Inhibitors of the same targets which were found to significantly increase T cell killing of tumor are indicated in the same color. BIRC2 (Blue bars), DNPEP (Red Bars), ITGAV (Green bars).

To determine whether any of the drugs could increase neo-antigen-reactive T cell elimination of epithelial tumor cells, we utilized two independent patient-derived colon cancer cell lines, SB4238 and SB4266. These lines were co-cultured with T cells engineered to express TCRs reactive against mutated proteins expressed by each tumor line. In the case of SB4238, a non-synonymous mutation in NCKAP1 (G1312T, **Figure S2C**) resulted in an D438Y amino acid substitution that was specifically recognized by a TCR identified in patient tumor infiltrating lymphocytes (TILs) **(Figure S2D, S2E)**. A TCR reactive against SB4266 through specific recognition of p53 R248W was previously described (Malekzadeh et al., 2019). We found that TCR-transduced T cells alone were capable of eliminating a modest percentage of SB4266 cells, and this increased in the presence of inhibitors of BIRC2 (birinapant and LCL161), DNPEP (CHR2797), and ITGAV (cilengitide) (**Figure 5E**). However, only BIRC2 inhibitors birinapant and LCL161 increased neo-antigen reactive T cell elimination of SB4238 tumor cells (**Figure 5F**).

As BIRC2 inhibition consistently increased T cell elimination of tumor cells irrespective of recognized antigens or cancer types, we next assessed whether the *in vivo* combination treatment of either LCL161 or birinapant along with antigen-specific T cells could increase tumor control. Indeed, we found that dosing mice bearing established B16 tumors with either LCL161 or birinapant increased the ability of PMEL-transgenic T cells to control tumor growth (**Figure 5G, S2F)** and increased overall survival time (**Figure 5H**). In summary, the use of a genome-scale CRISPR screen to inform the selection of inhibitors for increasing tumor susceptibility to T cell killing led to the identification of several druggable tumor targets, with BIRC2 displaying the greatest potential across several tumor antigens and histologies.

### BIRC2 negatively regulates tumor cell inflammation

As BIRC2 was found to be a component of the active tumor defense against T cell engagement and also could be targeted pharmacologically across a number of settings to increase the efficacy of T cell elimination of multiple tumor cell lines, we sought to further characterize the role and regulation of BIRC2 in tumor cells. Consistent with our observation that T cell engagement increased mRNA expression of *BIRC2* in tumor cells, treatment of A375 or Mel624 melanoma cells with recombinant IFNγ or TNFα (to mimic T cell inflammatory cytokine release) was sufficient to increase the protein expression of BIRC2 (**Figure 6A).** Inhibitors of BIRC2 have been reported to attenuate the cytotoxic effects of TNFα in tumor cells (Vredevoogd et al., 2019), yet genetic studies of BIRC2 function in the context of tumor cell resistance to immunotherapy have not been extensively performed. We utilized CRISPR/Cas9 with newly designed sgRNAs to specifically knock out BIRC2 (**Figure 6B**) and performed mRNA sequencing. BIRC2 knockout cells had increased mRNA expression of pro-inflammatory chemokines and cytokines including *CCL5*, *CSF2*, *CXCL10,* and *CXCL11*, along with increased expression of genes involved in T cell binding and recognition of tumor cells such as *ICAM1* and *HLA-DQA2* (**Figure 6C**). Pathway analysis of genes upregulated in BIRC2 knockout cells revealed enrichment for inflammatory signaling cascades, antigen presentation pathway, the unfolded protein response and NF-κB signaling (**Figure 6D**). Consistent with the observed changes in mRNA, we found that BIRC2 knockout cells also had increased protein level expression of ICAM1, along with HLA Class I (**Figure 6E**). We also found increased expression of a number of genes previously associated with response to immune checkpoint blockade (Harlin et al., 2009) in BIRC2 knockout cells (**Figure 6F**).

**Figure 6.**
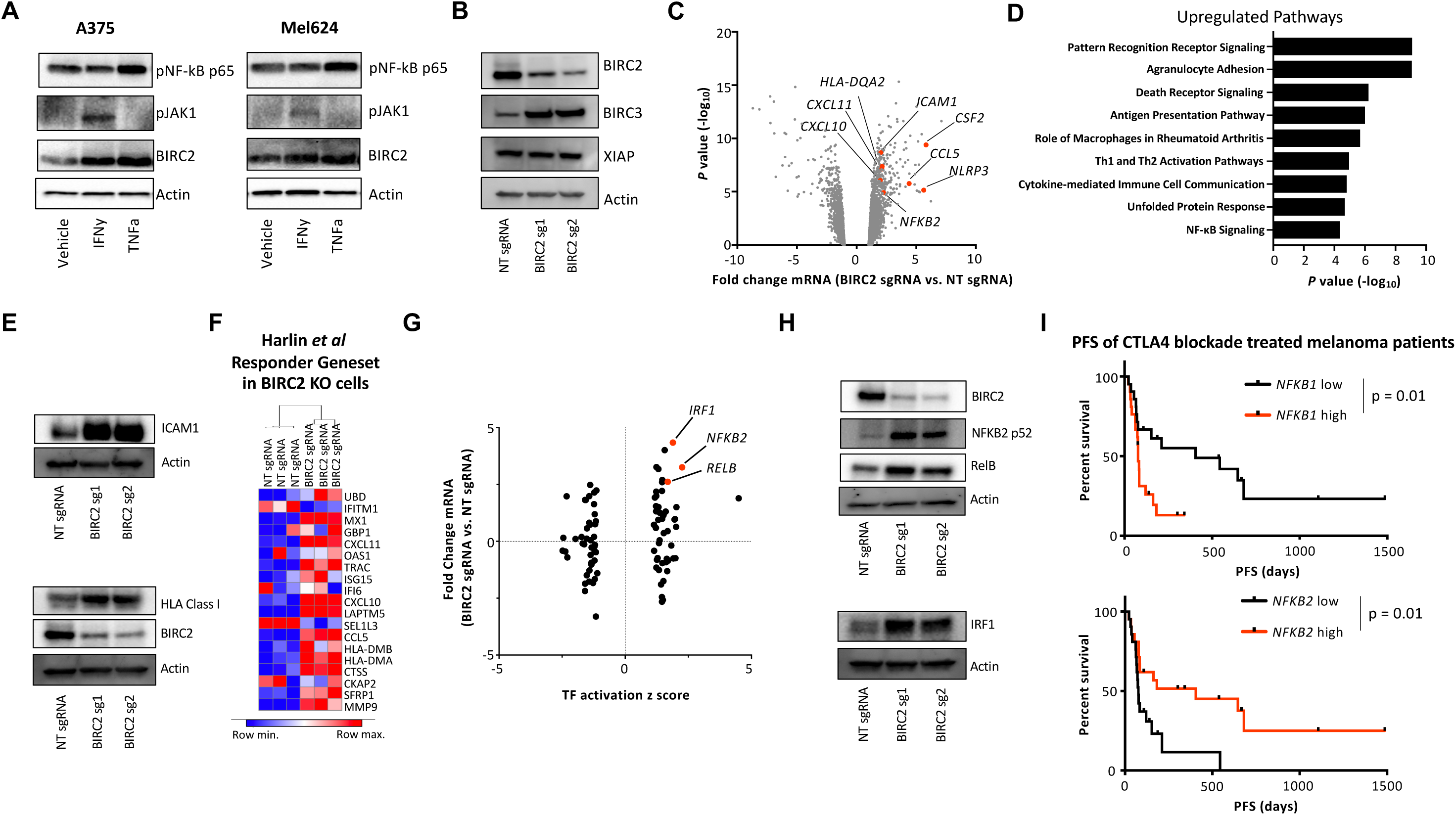
BIRC2 negatively regulates tumor inflammation and non-canonical NF-κB signaling. **(A)** A375 (left panel) or Mel624 (right panel) melanoma cells were treated with IFNγ (100 ng/mL) or TNFα (100 ng/mL) for 8hr and western blot analysis was performed. **(B)** A375 cells were transduced with Cas9 and non-targeting (NT sgRNA) or BIRC2 targeted sgRNAs and western blot analysis was performed. **(C)** Volcano plot representing differentially expressed genes in A375 cells transduced with Cas9 and BIRC2 targeting sgRNA versus NT sgRNA. Expression of genes was analyzed by RNA-seq as in Figure 2 with abundance represented as relative fold change (*x* axis) versus significance (*y* axis). **(D)** Ingenuity pathway analysis of genes significantly upregulated in BIRC2 KO A375 cells. **(E)** A375 cells were transduced with Cas9 and NT sgRNA or BIRC2 targeted sgRNAs and western blot analysis was performed. **(F)** Heatmap quantifying mRNA expression of genes reported to be differentially expressed in patients responding to checkpoint blockade therapy in melanoma (Harlin *et al*., 2009) in A375 cells transduced with Cas9 and NT sgRNA or BIRC2 targeting sgRNA. **(G)** Volcano plot representing transcription factor activation scores in mRNA expression profiles of BIRC2 KO A375 cells. Transcription factor activation was analyzed by IPA with increased transcription factor activity represented as positive *z* scores (*x* axis) and increased mRNA expression of transcription factor represented as positive fold change (*y* axis). **(H)** A375 cells were transduced with Cas9 and NT sgRNA or BIRC2 targeted sgRNAs and western blot analysis was performed. **(I)** Analysis of reported progression free survival of patients treated with CTLA4 blockade (Van Allen *et al*., 2015) based on mRNA expression of NFKB1 or NFKB2. Patients were stratified into groups based on whether they were above or below median mRNA expression for each gene (NFKB1=16.82, *NFKB2=*8.62). Data is representative of three independent experiments **(A, E, H)**, two independent experiment **(B)**, is pooled from a single experiment with three biological replicates per group **(C-D, F-G)** or represents analysis of 21 patients in each group **(I)**. *P* values calculated by log-rank test.

BIRC2 had previously been reported to act as a negative regulator of the non-canonical NF-κB signaling cascade (Yang et al., 2016), and Ingenuity Pathway Analysis of transcription factor activation showed increased activation of key non-canonical NF-κB regulators *NFKB2* and *RELB* in addition to *IRF1* in BIRC2 knockout cells (**Figure 6G**). We found that BIRC2 knockout cells had increased total expression of NFKB2, RELB and IRF1 (**Figure 6H**) and nuclear expression of NFKB2 and RELB was increased **(Figure S3A)** in BIRC2 knockout cells. Importantly, expression of NFKB2 is negatively correlated with the expression of BIRC2 in human melanoma patients **(Figure S3B)** and increased expression of non-canonical NF-κB transcription factor *NFKB2* was associated with increased patient survival, while the expression of canonical NF-κB transcription factor *NFKB1* was associated with decreased survival (**Figure 6I**) in melanoma patients treated with CTLA4 blockade (Van Allen et al., 2015). Additionally, using Tumor Immune Dysfunction and Exclusion (TIDE) computational analysis (Jiang et al., 2018), we determined that combined expression of *NFKB2* and *IRF1* predicts the probability of human patient response to checkpoint blockade across a number of previously published patient cohorts **(Figure S4A)**.

### BIRC2 loss promotes increased T cell migration and recognition of tumor cells

Tumor expression of chemokines including *CCL5* (Dangaj et al., 2019), *CXCL10* (Peng et al., 2015) and the cytokine *CSF2* (Borrello et al., 2009) have been linked with the ability of T cells to infiltrate and control tumor growth, and we found that increased mRNA expression of these genes is positively correlated with *NFKB2* expression in human melanoma patients **(Figure S5A-C)**. Consistent with a role for non-canonical NF-κB in promoting inflammatory chemokine production, we found increased abundance of RelB at the *CXCL10* promoter in BIRC2 KO tumor cells (**Figure 7A**). RelB and NFKB2 were also found to have increased abundance at the promoters of other genes linked with immunotherapy response (Harlin et al., 2009) including *HLA-DMB* **(Figure S5D)**, and *MMP9* **(Figure S5E)** in BIRC2 KO cells.

**Figure 7.**
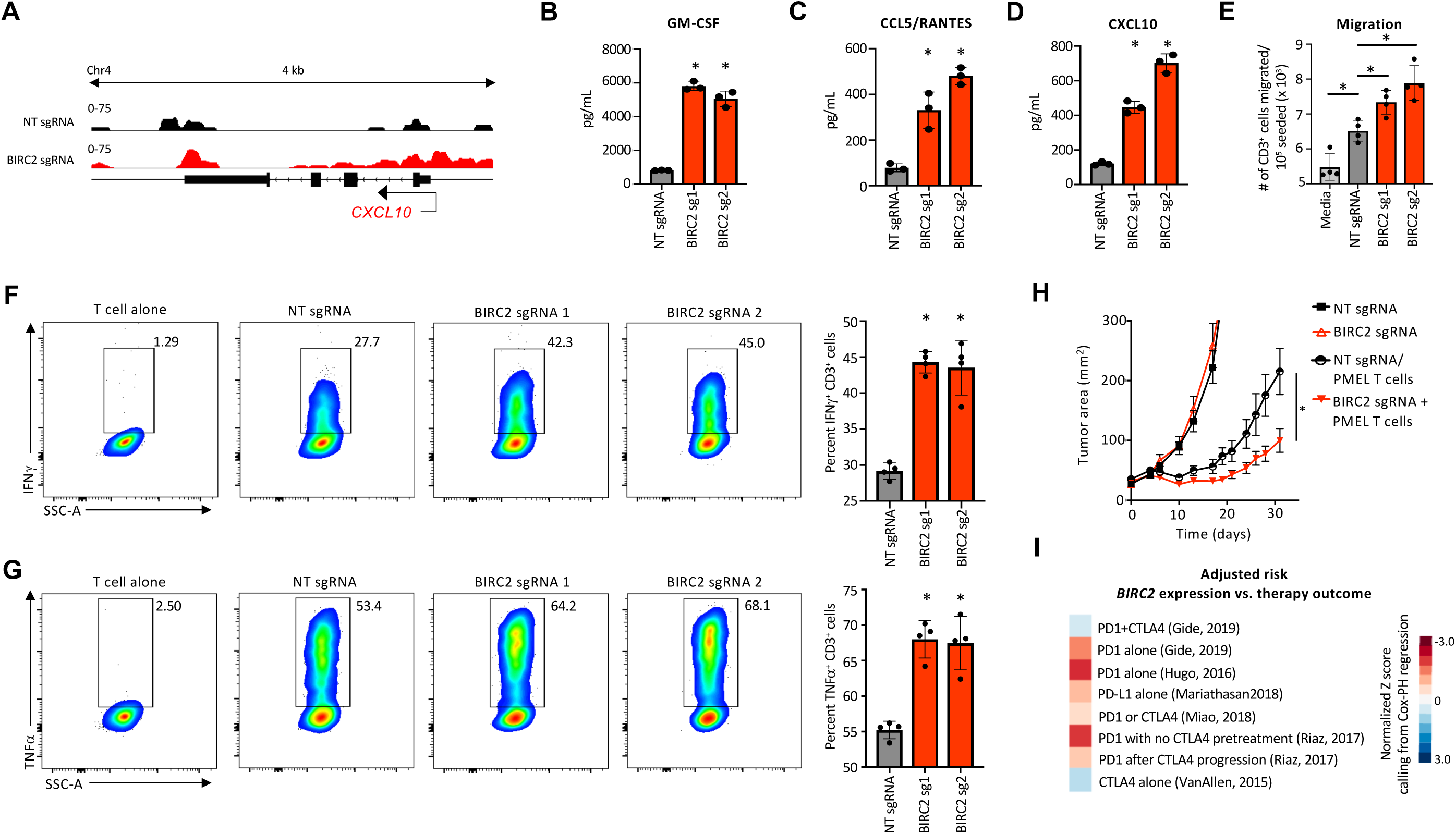
Genetic loss of BIRC2 increases tumor production of inflammatory chemokines and promotes T cell migration. **(A)** ChIP-seq analysis of RELB abundance at *CXCL10* locus in A375 cells transduced with Cas9 and NT sgRNA or BIRC2 targeting sgRNA. **(B-E)** A375 cells transduced with Cas9 and NT sgRNA or BIRC2 targeting sgRNA were assayed for production of **(B)** GM-CSF, **(C)** CCL5/RANTES, **(D)** CXCL10 and **(E)** ability to induce migration of ESO T cells across a transwell membrane. **(F-G)** Intracellular cytokine production of ESO T cells that were co-cultured with A375 cells transduced with Cas9 and NT sgRNA or BIRC2 targeting sgRNAs for 5 hr in the presence of brefeldin and monensin. **(H)** Growth curve of subcutaneous B16 melanoma tumors that were modified with Cas9 and NT sgRNA or BIRC2 targeting sgRNA. Mice were treated with PMEL1 T cells and tumor area was measured over time. **(I)** Heatmap displaying normalized *Z* scores from Cox-PH regression analysis of BIRC2 expression in published human immunotherapy datasets (Jiang et al., 2018). Data is representative of two independent experiments **(A-D),** three independent experiments **(E-G)**, three independent experiments with at least 8 mice per group **(H)**. \**P* < 0.05. *P* values calculated by two tailed Student’s *t* test.

We found that BIRC2 KO cells secreted significantly higher levels of GM-CSF, CCL5 and CXCL10 proteins (**Figure 7B-D**). These unexpected findings suggested that BIRC2 might limit tumor production of these critical factors. Consistent with higher levels of these chemokines, genetic knockout of BIRC2 in tumor cells promoted increased migration of ESO T cells across a transwell membrane (**Figure 7E**). Co-culture of ESO T cells with BIRC2 knockout tumor cells also resulted in increased levels of intracellular IFNγ (**Figure 7F**) and TNFα (**Figure 7G**) in the T cells. We tested the effect of genetic knockout of BIRC2 on tumor susceptibility to ACT *in vivo*. We observed that transducing B16 melanoma cells with BIRC2-targeting sgRNAs along with Cas9 resulted in increased non-canonical NF-κB pathway activity **(Figure S5F)** and increased tumor control after adoptive transfer of syngeneic PMEL TCR transgenic T cells compared with cells transduced with a non-targeting sgRNA (**Figure 7H**). Finally, *BIRC2* expression in human cancer patients treated with immune checkpoint blockade was also found to correlate with risk of adverse patient outcome (Jiang et al., 2018) (**Figure 7I**). Together, these data indicate that BIRC2 inhibition may be a potential opportunity to increase the efficacy of immunotherapy-mediated tumor cell killing across multiple cancer types.

## Discussion

Tumor immunotherapy has reshaped clinical strategies for the treatment of human cancer and has brought about dramatic improvements in patient outcomes with several types of cancer. In spite of these improvements, most patients still receive minimal or non-durable clinical benefit from immunotherapy treatment strategies (Brahmer et al., 2012). Understanding the mechanisms by which tumor cells resist immunotherapy treatments and developing new therapies to overcome resistance mechanisms and improve the therapeutic efficacy of immunotherapy against cancer is an important focus. Using a two-cell type (2CT) whole-genome CRISPR/Cas9 screen, we have profiled tumor genes that are essential for survival in the face of engagement with tumor-specific T cells. Integrating dynamic transcriptional profiling of the tumor response to T cell engagement with the results of the genome-scale CRISPR screen allowed us to identify tumor genes that are both transcriptionally induced by T cell encounter and function to promote tumor cell survival in this context. Genes implicated in tumor resistance to conventional therapies (Jung et al., 2015; Sartorius and Krammer, 2002), including *BCL2* and *BIRC2,* were among the tumor genes we identified that can negatively regulate tumor cell susceptibility to T cell-mediated destruction. In addition, we identified genes with no previously described role in promoting tumor cell survival, including *TOR1AIP1*, *SCAF11* and *LPGAT1*. The role and regulation of these genes in the context of immunotherapy requires further investigation, but our data suggests that, taken together, these genes represent components of an orchestrated tumor cell defense mechanism that limits the cytotoxic effect of T cells on tumors.

In addition to identifying components of the active tumor defense against T cell engagement at a genome scale, we also assessed the therapeutic potential of targeting these and other top genes identified from the 2CT CRISPR to augment the efficacy of immunotherapy. By interrogating top hits from the screen with a database of druggable proteins, we identified a number of potential tumor targets for combination therapy. We found that the inhibition of several of these targets, including BIRC2, ITGAV and DNPEP, increased tumor antigen-specific T cell-mediated destruction of melanoma cells *in vitro* and BIRC2 inhibition could also augment the *in vitro* destruction of epithelial tumor cells by neo-antigen-specific T cells. Inhibition of BIRC2 during ACT treatment of a murine model of established melanoma demonstrated that two separated BIRC2 inhibitors were capable of increasing T cell control of melanoma growth. The pharmacological inhibition of BIRC2 has been proposed as a potential avenue to increase tumor cell susceptibility to TNFα-mediated toxicity (Vredevoogd et al., 2019). Taken together, these data suggest that BIRC2 inhibitors may increase the effects of cancer immunotherapy in human patients. This could be especially beneficial in tumor histologies for which current immunotherapy treatment efficacy is low.

Although a number of functional genomics studies have identified BIRC2 as a negative regulator of susceptibility to immunotherapy (Manguso et al., 2017; Pan et al., 2018; Vredevoogd et al., 2019), mechanistic insight into the role of BIRC2 in promoting tumor cell survival in this context have remained limited. While BIRC2 and other related proteins are often thought to inhibit cellular apoptosis through inhibiting the pro-apoptotic activities of caspases, additional functions have been characterized (Dubrez-Daloz et al., 2008). Further, BIRC2 has been reported to bind, but not inhibit the function of caspases (Eckelman and Salvesen, 2006), suggesting that direct inhibition of cellular apoptosis may not account for the effects of BIRC2 on tumor susceptibility to immunotherapy. Consistent with previous reports in the setting of B cell malignancies (Yang et al., 2016), we found that BIRC2 acts as an important negative regulator of the non-canonical NF-κB signaling cascade. Activation of non-canonical NF-κB signaling by genetic deletion of *BIRC2* promoted increased tumor cell destruction by upregulating tumor cell antigen presentation pathways and the production of pro-inflammatory chemokines that have been linked with clinical responses to immunotherapy (Dangaj et al., 2019; Harlin et al., 2009; Peng et al., 2015). It is notable that the expression of non-canonical NF-κB transcription factor *NFKB2* was positively correlated with survival in patients treated with CTLA blockade (Van Allen et al., 2015) and that *NFKB2* and *IRF1* expression could predict responses across several clinical cohorts of human immunotherapy patients. Further work to study the direct role of the non-canonical NF-κB signaling pathway in regulating the efficacy of immunotherapy will be necessary to establish a role for the pathway in regulating tumor susceptibility apart from BIRC2-mediated effects.

Overall, our work here describes the use of genome-scale functional genomics to rapidly identify targets for combination therapies to increase the efficacy of cancer immunotherapy. Combining functional genomics studies in a two-cell type setting with transcriptional dynamics studies can also be utilized to identify active, acute regulators of resistance and susceptibility to selective pressures. As recent works have demonstrated the possibility of conducting genome-scale functional genomics studies in primary T cells (Shifrut et al., 2018), similar approaches might be utilized to identify T cell-intrinsic negative regulators of anti-tumor effects for targeted inhibition to improve immunotherapy. Integrating functional genomics data from both tumor and T cells settings may help to optimize new immunotherapy strategies to promote increased therapeutic effects and improved patient outcomes.

## Acknowledgments

The authors of this article were supported by the Center for Cell-Based Therapy, NCI, NIH (Bethesda, MD)), NIH Center for Regenerative Medicine, the Milstein Family Foundation, and the Intramural Research Program of the NCI (ZIA BC010763). We would like to thank Erina He for illustrations used in the manuscript. The authors would like to thank members of the Restifo, Rosenberg and Yang labs for helpful discussions and suggestions. N.E.S. is supported by NYU and NYGC startup funds, NIH/NHGRI (R00HG008171, DP2HG010099), NIH/NCI (R01CA218668), DARPA (D18AP00053), the Sidney Kimmel Foundation, the Melanoma Research Alliance, and the Brain and Behavior Foundation.

## Author contributions

R.J.K. and S.J.P. and N.P.R. designed the study and wrote the manuscript. R.J.K., S.J.P., S.K.V., A.E.D., Y.P., M.S., T.N.Y., Z.Y., M.J., W.L., A.P., P.M., D.D.D., A.N.H., N.E.S. performed experiments or provided reagents. K.C.W. and N.E.S. provided reagents and expertise.

## Methods

### Human specimens

Peripheral blood mononuclear cells (PBMCs) were isolated from healthy donors and tumor samples were isolated from patients with melanoma and GI cancers. All human specimens were collected with informed consent and procedures approved by the institutional review board (IRB) of the National Cancer Institute (NCI).

### Mice

All animal experiments were approved by the Institutional Animal Care and Use Committees of the NCI and were performed in accordance with NIH guidelines. C57BL/6NCR mice were obtained from Charles River Laboratories at NCI Frederick. B6.Cg-*Thy1^a^*/Cy Tg(TcraTcrb)8Rest/J (PMEL1) mice were purchased from Jackson Laboratory. All mice were maintained under specific pathogen-free conditions. Female mice aged 6-8 weeks were used for *in vivo* experiments.

### Cell culture

Melanoma cell lines Mel624.38 were isolated from surgically resected metastases as previously described (Robbins et al., 2008) and were cultured in RPMI 1640 (Invitrogen) medium supplemented with 10% fetal bovine serum (FBS, Hyclone, Logan, UT), 2 mM L-glutamine and 1% penicillin-streptomycin. A375 melanoma cells were obtained from the American Type Culture Collection (Manassas, VA) and cultured in RPMI 1640 medium supplemented with 10% FBS, 2 mM L-glutamine and 1% penicillin-streptomycin. Transduced T lymphocytes were cultured in RPMI 1640 supplemented with 10% FBS, 2 mM L-glutamine and 1% penicillin-streptomycin.

### Retroviral transduction of human T cells with TCRs

Retroviral vectors for TCRs recognizing the HLA-A*02-restricted melanoma antigens NY-ESO-1 (NY-ESO-1:157-165 epitope) and MART-1 (MART-1:27-35 epitope, DMF5) were generated as previously described (Johnson et al., 2006; Robbins et al., 2008). Neo-antigen-reactive TCRs were identified and expressed in primary human T cells as previously described (Lo et al., 2019; Malekzadeh et al., 2019). For transduction of human T cells, CD8^+^ T cells seeded at 2 x 10^6^ cells per well in a 24-well plate were stimulated with anti-CD3 antibody OKT3 (5 ug/mL coated) and anti-CD28 antibody (5 ug/mL soluble) along with IL-2 (200 IU/mL) on day 0. Non-tissue culture treated 24-well plates were coated with 0.5 mL per well of 10 μg/mL RetroNectin (Takara) on day 1 and stored overnight at 4°C. Vector supernatant (1 mL per well) was added to plates on day 2 followed by centrifugation at 2000g for 2h at 32°C. 800 μL was aspirated and T cells were added at 1 x 10^6^ cells/mL, centrifuged for 10 min at 1500 rpm and incubated overnight. A second transduction was performed the following day as described above. Cells were subsequently maintained in culture at 1 x 10^6^ cells/mL and expanded until day 10, after which they were used or cryo-preserved for future use.

### Lentiviral production and purification

Lentiviral particles were produced and purified as described previously (Patel et al., 2017). In brief, HEK293FT cells (Invitrogen) were cultured in DMEM supplemented with 10% FBS, 2 mM L-glutamine and 1% penicillin-streptomycin. One day prior to transfection, HEK293FT cells were seeded in T-225 flasks at 60% confluency. One hour prior to transfection, media was aspirated and replaced with 13 mL OptiMEM media (Invitrogen). Each flask of cells were transfected with 100 μL Lipofectamine 200 and 200 μL Plus reagent (Invitrogen) along with 20 μg of lentiCRISPRv2 plasmid or pooled plasmid human GeCKOv.2 (Genome-scale CRISPR knockout) library, 15 μg psPAX2 and 10 μg pMD2.G. 6-8 hr after transfection, media was replaced with 20 mL of DMEM supplemented with 10% FBS and 1% BSA. Media containing viral particles was collected 48 hr post-transfection and titer was assayed with Lenti-X GoStix (Clontech). Viral supernatant was centrifuged at 3,000 r.c.f. at 4°C for 10 min followed by filtration through 0.45 μm low-protein binding membrane. For pooled library plasmids, viral supernatants were concentrated by centrifugation at 4,000 r.c.f at 4°C for 35 min in Amicon Ultra-15 filters (Millipore Ultracel-100K). Concentrated viral supernatants were stored in aliquots at −80°C.

### 2CT T cell and tumor cell co-culture

Two days prior to co-culture, T cells were thawed in T cell media containing 3 U/mL DNase (Genentech Inc.) overnight. Tumor cells were seeded at desired density on this day in T cell media. After 24 hr, T cells were cultured in T cell media supplemented with 300 IU/mL IL-2 for 24 hr. T cells were co-cultured with tumor cells at various effector:target (E:T) ratios for specified durations. After co-culture, T cells were removed by washing tumor cells with PBS and tumor cells were detached using trypsin. Cells were stained with fixable Live/Dead dye (Invitrogen) followed by human anti-CD3 antibody (clone SK7, BD) in FACS staining buffer (PBS + 0.2% BSA). Cell counts were normalized with CountBright Absolute Counting Beads (Invitrogen) by FACS.

### 2CT GeCKOv.2 screens, genomic DNA extractions and screen analysis

2CT genome-wide CRISPR screens were performed as previously described (Patel et al., 2017), with some modifications. In brief, Mel624 cells were transduced independently with both A and B GeCKOv.2 libraries. For each screen, cells were split into two groups of 5×10^7^ transduced Mel624 cells. One group was co-cultured with ESO T cells at an E:T ratio of 1:3 for each library. A second group of transduced Mel624 cells were cultured under the same density and conditions, but without ESO T cells. The co-culture phase was maintained for 6 hr, after which the T cells were removed as described above. The recovery phase was maintained for another 48 hr and surviving cells were frozen to evaluate sgRNA depletion. gDNA was extracted from frozen tumor cells using previously optimized (Chen et al., 2015) ammonium acetate and alcohol precipitation procedure to isolate gDNA with AL buffer (Qiagen) substituted for the initial cell lysis step. sgRNA abundance was detected as previously described (Patel et al., 2017).

### Arrayed validation of targets with CRISPR/Cas9

Selected targets that were identified from the genome-scale CRISPR screen and DTA analysis were further studied using additional sgRNA targeting sequences. We utilized 3 newly designed sgRNA guide sequences for each gene as listed in **Table 6**. We cloned these sgRNAs into the lentiGuide-Puro vector as previously described (Sanjana et al., 2014). A375 cells were transduced with lentiCas9-Blast (Addgene) and selected with 5 μg/mL blasticidin for 10 days, after which cells were transduced with lentiGuide-Puro sgRNA constructs. Cells were then selected with 1 μg/mL puromycin for 7 days. In some experiments (**Figures 6-7**), sgRNA guide sequences were cloned into the lentiCRISPRv2 vector for mechanistic studies as previously described (Patel et al., 2017).

### RNA sequencing and analysis

mRNA-seq was performed on Mel624 melanoma cells and NY-ESO-1 TCR transduced primary human T cells that were co-cultured and separated into purified populations by FACS. Separate sequencing was performed on A375 melanoma cells that were transduced with Cas9 and non-targeting sgRNA or Cas9 and BIRC2-targeting sgRNA. Libraries were prepared using TruSeq RNA sample prep kit (FC-122-1001, Illumina). Approximately 40 million reads were sequenced and aligned to the human genome (hg19 for Mel624/NY-ESO-1 T cell experiment, hg38 for A375 experiments) with TopHat 2.0.11. Uniquely retained mapped reads were used to calculate differentially expressed genes using edgeR or Cuffdiff. Fisher’s exact test or t-tests were used to calculate significance with indicated *P* values and fold-change thresholds. To filter for RNA contamination derived from incomplete FACS-based purification of ESO T cells and Mel624 melanoma cells, we filtered out genes that had 100-fold enrichments in expression in the opposing cell type under basal conditions. RNA-sequencing raw data files are deposited at GEO-GSE137824.

### ChIP-sequencing

Chromatin Immunoprecipitations were performed following manufacturer’s instructions (ChIP-IT Express Shearing Kit, Active Motif). In brief, A375 cells transduced with Cas9 and non-targeting sgRNA or with Cas9 and BIRC2-targeting sgRNA were fixed with formaldehyde for 7 min on a rocking platform and quenched with glycine solution. Cells were pelleted with PMSF and protease inhibitor cocktail and stored at −80°C prior to lysis. Cells were suspended in ice cold lysis buffer to extract nuclear material, which was then sheared by incubating with enzymatic cocktail for 10 min at 37°C. 40 μg of sheared chromatin/sample was incubated with protein G magnetic beads with α-RelB (10544, Cell Signaling) or α-NF-κB2 antibodies (37359 Cell Signaling) overnight. Magnetic beads were washed with buffers to remove unbound immune complexes and chromatin was eluted with 150 μL of elution buffer. Reverse crosslinking was performed on eluted DNA, followed by phenol chloroform extraction. Samples were sequenced on a HiSeq4000 in paired-end mode with approximately 40 million reads/sample. Sequenced reads were trimmed for adapters and aligned to the human genome (hg38) with Bowtie v2 and uniquely mapped reads were retained. The output of Bowtie was converted to BAM files, normalized using RPKM and converted to coverage tracks in big wig format using Deeptools (Command #BamCoverage -b Bam_File –normalizeUsingRPKM – binSize 10 –smoothLength 30 -bl hg38.blacklist.bed –centerReads –minMappingQuality 30 -o Output_File.bw). Tracks generated were viewed using the IGV (Integrative Genomics Viewer).

### Tumor Cell Chemokine and Cytokine Secretion

A375 cell production of chemokine proteins was quantified using a CBA Human Chemokine Kit (BD Biosciences Cat #552990) and GM-CSF production was measured using Human GM-CSF ELISA Kit (Abcam Cat #ab174448). In brief, A375 cells transduced with Cas9 and non-targeting sgRNA or BIRC2-targeting sgRNAs were plated at 4 x 10^5^ in a 6-well dish and cultured for 24 hr. Media supernatant was collected and spun at 1500rpm for 10 min to pellet cells and debris. Supernatant was collected for analysis. In some cases, media was diluted 1:5 using assay diluent to ensure detected values were within standard calibration curves.

### Tumor Pathway Activation Screen

Tumor pathway activation was performed as previously described (Martz et al., 2014), with minor modifications. In brief, lentiviral particles were generated for each pathway activation construct and A375 melanoma cells were transduced by spinfection at an MOI of 0.3. Transduced cells were incubated for 24 hr, followed by puromycin selection. After selection, cells were plated and 2CT assay was conducted with ESO T cells at a 1:3 ratio for 16hr.

### Lentiviral transduction of tumor cells

Lentiviral transduction of tumor cells was performed as previously described (Patel et al., 2017). In brief, tumor cell culture media was replaced with RPMI 1640 media supplemented with 10% FBS and 8μg/mL polybrene. Lentiviral particles were added at an MOI of 0.3, followed by spinfection at 1500 rpm for 30 min. Following this, cells were incubated for 24 hr, after which the culture media was aspirated and replaced with RPMI 1640 media supplemented with 10% FBS and selection antibiotic. For puromycin-resistant constructs, puromycin was added at 1μg/mL. For blasticidin-resistant constructs, blasticidin was added at 5μg/mL. Cells were maintained under antibiotic selection until non-transduced control cells were eliminated by drugs.

### High throughput inhibitor screen

A375 melanoma cells were plated at 1×10^4^ cells/well in 150μL of complete RPMI in a 96-well flat bottom plate and incubated for 24 hr. Inhibitors were then added in 50μL of complete RPMI (concentrated to reach desired concentration after addition), followed by addition of ESO T cells at a 1:3 E:T ratio in complete RPMI to a total of 250μL. Cells were incubated for 16 hr, after which cells were washed twice with PBS, and WST1 reagent (Sigma Aldrich) was added in complete RPMI and cell viability was assessed.

### Inhibitor sources

Inhibitors were purchased from commercial suppliers as listed below, or were provided (Compound 29, (Patch et al., 2011)) by the Donald McDonnell Laboratory (Duke University). ABT199 (Selleckchem #S8048), ABT263 (Selleckchem #S1001), LCL161 (Selleckchem #S7009), Birinapant (Selleckchem #S7015), Fomepizole (Selleckchem #S1717), CHR2797 (Tocris #3595), XCT790 (Sigma Aldrich #X4753), Topiramate (Selleckchem #S1438), ACET (Tocris #2728), Almorexant (Selleckchem #S2160), SB334867 (R&D Systems, #1960), anti-IL1R1 (R&D Systems, #AF269), anti-IL9 (R&D Systems, #AF209), anti-ITGAV (Abcam, #ab16821), Cilengitide (Selleckchem, #S7077), Guanidine HCl (Sigma Aldrich, #G3272), Dalfampridine (Sigma Aldrich, #275875), BEZ235 (Selleckchem, #S1009), Apitolisib (Selleckchem, #S2696), S17092 (Sigma Aldrich, #SML0181), WNK463 (Selleckchem, #S8358).

### FACS-based 2CT inhibitor validations

Tumor cells were plated at 3×10^4^ cells/well in 250μL of complete RPMI in a 48-well flat bottom plate and incubated for 24 hr. Inhibitors were then added in 50μL of complete RPMI, followed by addition of TCR-transduced T cells at a 1:3 E:T ratio in complete RPMI to a total of 400μL. Cells were incubated for 16hr, and analyzed as described for 2CT assay.

### Murine ACT models

For *in vivo* immunotherapy experiments, we utilized previously described (Hanada et al., 2019) B16 melanoma cells expressing a chimeric mouse-human gp100 antigen (B16mhGP100). We generated gene-deleted B16mhGP100 cells using lentiviruses encoding sgRNAs targeting *Birc2* as described above. C57BL/6 mice were subcutaneously implanted with 5×10^5^ B16mhGP100 cells and tumors were allowed to grow for 10 days. Mice were then irradiated (6 Gy) and injected intravenously with 5×10^6^ Pmel CD8^+^ T cells. Mice received intraperitoneal injections of IL-2 in PBS (6×10^4^ IU in 0.5mL) once daily for 3 consecutive days. In certain experiments, mice received intraperitoneal injections of LCL-161 or birinapant in Captisol (30% in H2O acidified to pH 4 with citric acid). Injections of inhibitors was performed every other day for 5 total injections. Tumor measurements were performed in a blinded fashion by an independent investigator approximately every two days after T cell transfer. Tumor area was calculated as length x width of the tumor. Mice with tumors in excess of 400 mm^2^ were euthanized.

### Pathway Enrichment Analysis

For gene pathway analysis, enriched or depleted genes were examined for gene category over-representation using Ingenuity Pathway Analysis (QIAGEN). For analysis of CRISPR depleted genes, the top 250 significant genes by RIGER analysis were selected. For analysis of RNA-sequencing experiments, all differentially expressed genes were analyzed. Fisher’s exact test (*P* < 0.05) was used to compute significance for over-representation of genes in a pathway.

### cBioportal analysis of human melanoma patient datasets

Analysis of human melanoma patient gene expression was performed using the Skin Cutaneous Melanoma geneset (TCGA, PanCancer Atlas). Samples were selected for which mRNA data was available and gene expression correlation analysis was performed.

### Identification of druggable targets for screening

Validated tumor resistance genes (**Figure 3B**) along with the top 250 depleted genes from CRISPR screens as calculated by rank-sum of replicate screens was analyzed by PANTHER (Mi et al., 2019) to map gene list to protein coding genes. 237 identified protein coding genes were analyzed using DGIDB (Griffith et al., 2013) to identify genes that are druggable. Genes were analyzed as ‘druggable gene category results’ and filtered on categories ‘inhibitor, allosteric modulator, antagonist, blocker, channel blocker, desensitize the target, gating inhibitor, incorporation into and destabilization, inhibitor, competitive inhibitor, inhibitory allosteric modulator, inhibitory immune response, intercalation, inverse agonist, negative modulator, neutralizer, partial antagonist, reducer, suppressor’. Results were manually curated for commercial availability of inhibitors.

### T cell migration

Tumor cells were plated at 1×10^4^ cells in complete RPMI in a 96-well receiver plate (Corning #3382) and incubated for 24h. Following this, ESO T cells were added to a 96-well transwell plate (Corning #3387) at 1×10^5^ cells/well in 100μL complete RPMI and the T cell plate was added to the tumor cell plate and incubated for 5 hr. Media supernatant from tumor cell wells was collected, wells were washed with PBS which was also collected, then tumor cells were detached with trypsin and collected. Total contents of each well were stained with fixable viability dye and αCD3e, and migrated T cells were enumerated by FACS counting.

### Flow cytometry

Tumor cells or T cells suspended in FACS staining buffer were stained with fluorochrome-conjugated antibodies against CD3e (SK7, BD) and murine TCRβ (12-5961-82, ThermoFisher). Cell viability was determined using propidium iodide exclusion or fixable Live/Dead kit (Invitrogen). Intracellular staining assay on ESO T cells was conducted after 5-6 hr co-culture with non-targeting sgRNA or BIRC2-targeting sgRNA modified A375 cells in the presence of monensin (BD, 512092KZ) and brefeldin A (BD, 512301KZ). Staining was performed using manufacturer’s instructions using antibodies against IFNγ (25723.11, BD) or TNFα (12-7349-41, ThermoFisher). Flow cytometric data were acquired using either a FACSCanto II or LSRII Fortessa cytometer (BD), and data were analyzed using FlowJo version 10.5.3 software (FlowJo LLC).

### Immunoblotting

Immunoblotting was performed as described previously (Jacobs et al., 2008). Primary antibodies were followed by mouse- or rabbit-conjugated horseradish peroxidase (HRP) secondary antibodies. HRP-conjugated antibodies (anti-mouse or anti-rabbit IgG HRP conjugate, Promega) were detected by enhanced chemiluminescence detections (Thermofisher) using a Biorad ChemiDoc MP. This included the following antibodies: BIRC2 (108361, Abcam), BIRC3 (3130, Cell Signaling), HLA Class I (70328, Abcam), ICAM1 (4915, Cell Signaling), IRF1 (8478 Cell Signaling), phospho-JAK1 (3331, Cell Signaling), Lamin A/C (4777, Cell Signaling), phospho-NF-κB p65 (3033, Cell Signaling), NF-κB2 p100/p52 (4882, Cell Signaling), RelB (10544, Cell Signaling), XIAP (2042, Cell Signaling). Alternatively, primary antibodies were followed by fluorescently labeled anti-mouse or rabbit antibodies and imaged using a Biorad ChemiDoc MP. This included the following antibodies: β-actin (A5441, Sigma).

### Nuclear and cytoplasmic extraction and fractionation

Cells were collected and cytoplasmic and nuclear protein extraction and fractionation protocol was performed as specified in manufacturer protocol (Thermofisher, Cat #78835). Extracted proteins were analyzed by immunoblot. 45 μg protein per lane was loaded for cytoplasmic fractions and 17.5 μg protein per lane was loaded for nuclear fractions.

### Statistical Analysis

Sample sizes were determined by prior experience and estimated power calculations. Statistical analysis was performed with Prism (GraphPad Software Inc.). Data were compared using either a two-tailed Student’s *t* test or one-way ANOVA with multiple comparisons corrected with Dunnett adjustment. *P* values < 0.05 were considered significant. For adoptive transfer experiments, mice were randomized prior to cell transfer and treatments. Tumor treatment graphs were compared by using the Wilcoxon rank sum test and analysis of animal survival was analyzed by a log-rank test.

## Supplemental Figure Titles and Legends

**Figure S1, related to Figure 1.**
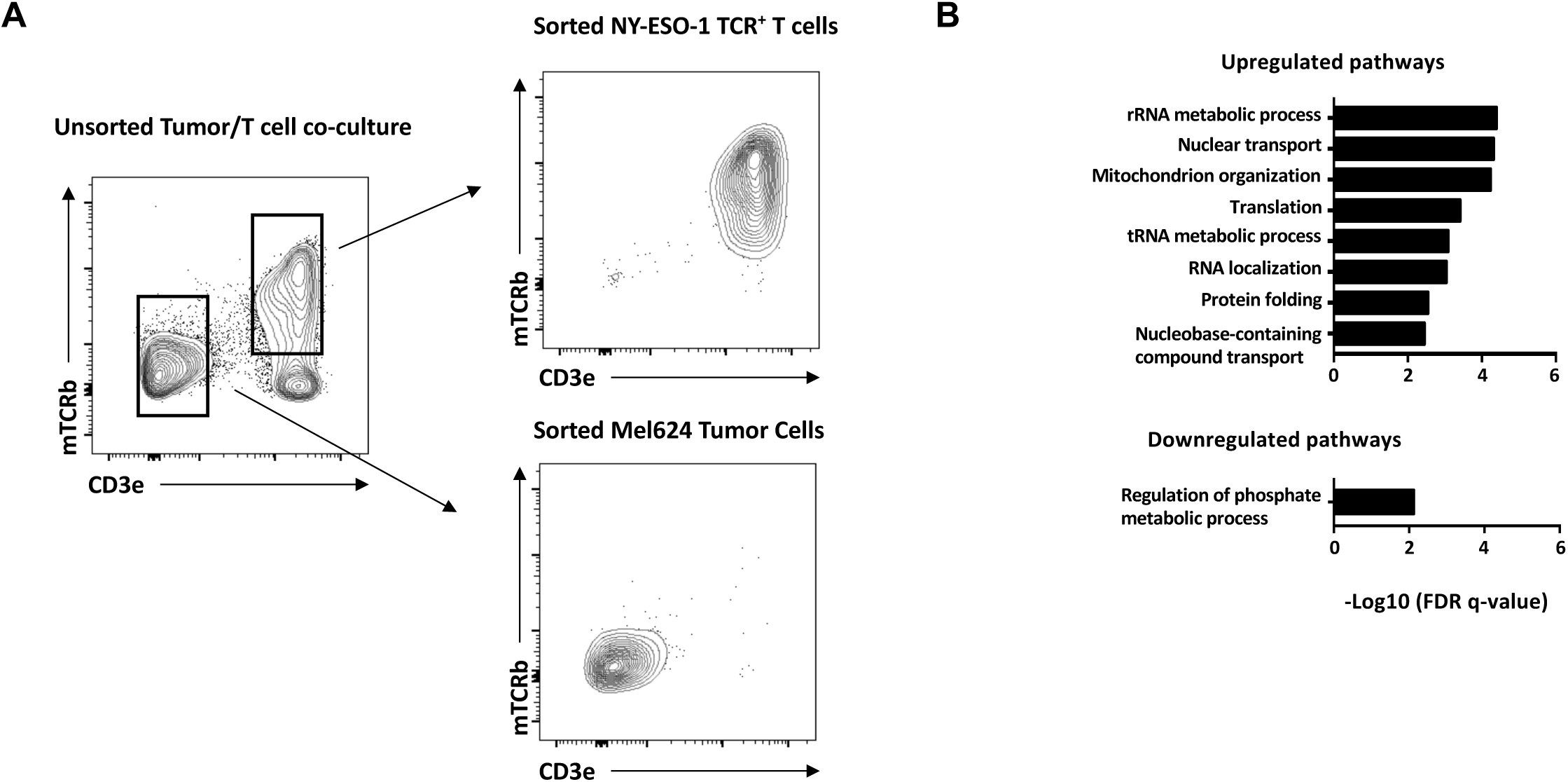
Tumor encounter drives T cell anabolic growth. **(A)** Sorting strategy for isolation of NY-ESO-1 TCR-expressing T cells and Mel624 melanoma cells following co-culture. **(B)** Ingenuity Pathway Analysis of T cell genes that were differentially expressed following co-culture with Mel624 melanoma cells. Data is pooled from three independent experiments **(B)**.

**Figure S2, related to Figure 4.**
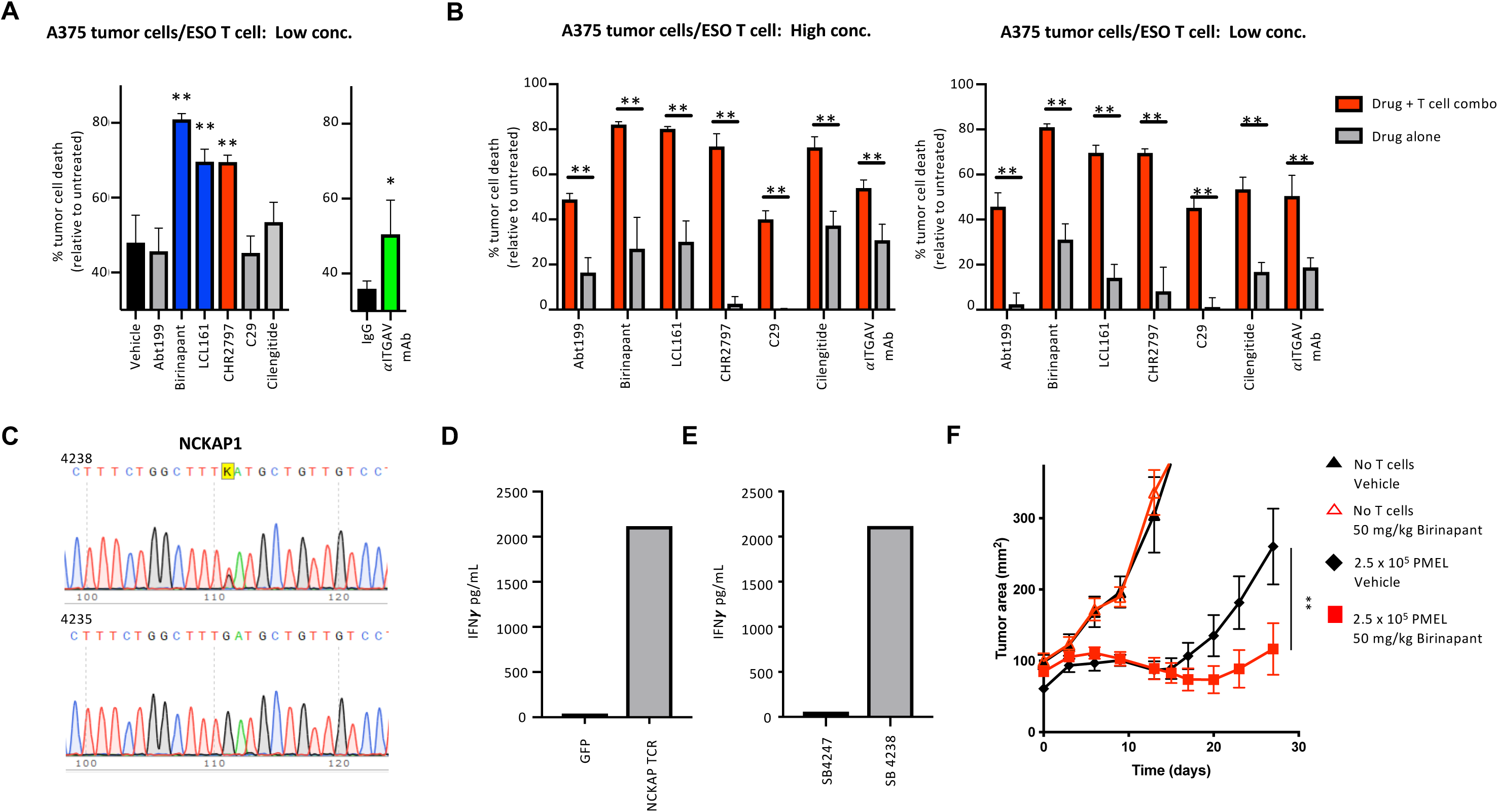
Combining inhibitors with tumor-specific T cells increases tumor cell elimination. **(A)** A375 melanoma cells were co-cultured with ESO T cells along with inhibitors at 500 nM except for α-ITGAV at 0.25 μg/mL and tumor cell elimination was measured by flow cytometry. **(B)** A375 melanoma cells were co-cultured with or without ESO T cells along with inhibitors at (left panel) 5μM except for α-ITGAV at 2.5 μg/mL (left panel) or at 500 nM except for α-ITGAV at 0.25 μg/mL (right panel). Tumor cell elimination was measured by flow cytometry. **(C)** Genomic DNA from SB4238 (NCKAP1 mutant) and SB4235 tumor lines (NCKAP1 wild type) was collected and Sanger Sequencing was performed at the NCKAP1 locus to confirm the presence of a mutation. **(D)** Primary human T cells were transduced with GFP or a NCKAP1 reactive TCR and co-cultured with SB4238 tumor cells for 16h. ELISA was performed to determine IFNγ secretion. **(E)** Primary human T cells were transduced with a NCKAP1 reactive TCR and co-cultured with SB4247 or SB4238 tumor cells for 16h. ELISA was performed to determine IFNγ secretion. **(F)** Subcutaneous tumor growth in mice receiving ACT of PMEL1 T cells along with vehicle or Birinapant (50 mg drug/kg body weight). Mice were treated with Birinapant via IP injection every 48hr beginning 24hr after PMEL1 T cell infusion for a total of 5 doses. Data are representative of four **(A-B)** or two experiments **(D-F).** * *P* < 0.05, ** *P* < 0.01. *P* values for *in vitro* assays **(A-C)** calculated by one-way ANOVA with multiple comparisons corrected with Dunnett adjustment. *P* values for *in vivo* assays **(F)** calculated by Wilcoxon rank sum test.

**Figure S3, related to Figure 6.**
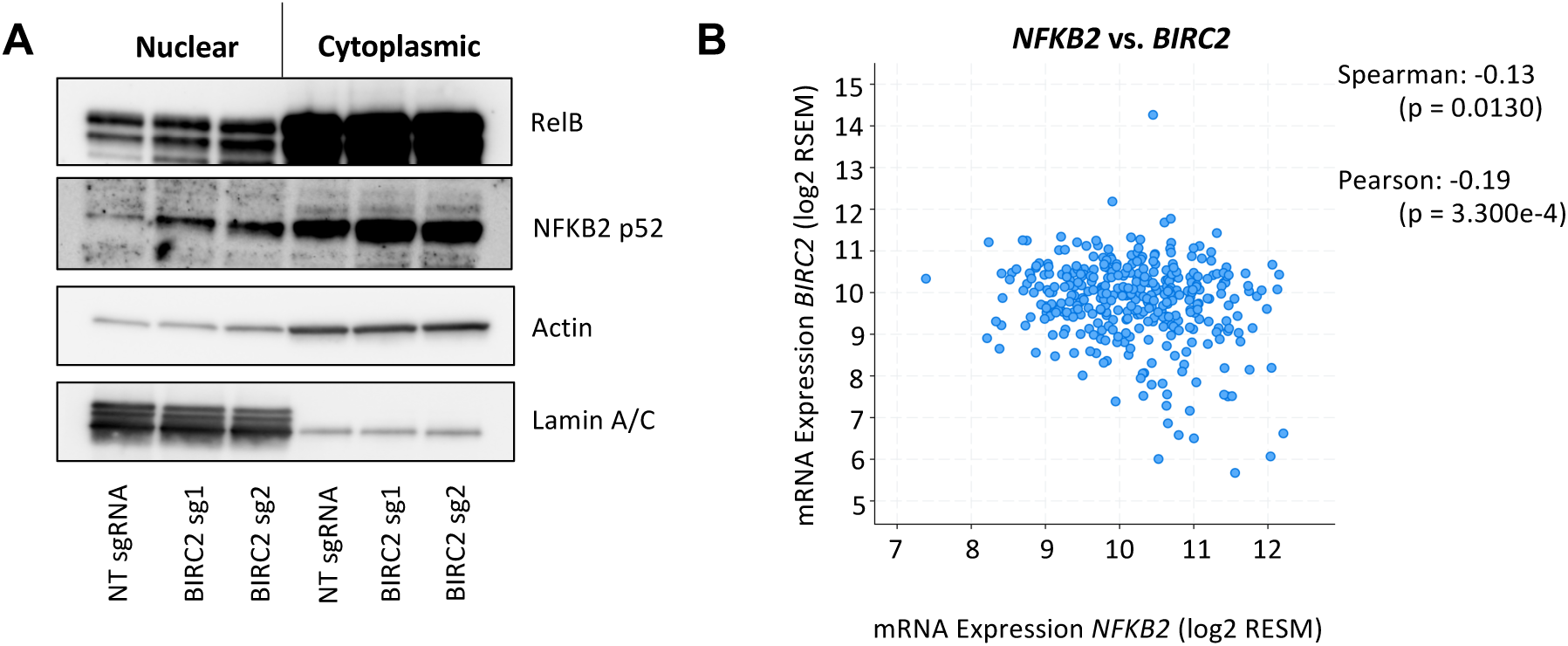
BIRC2 inhibits non-canonical NF-κB activity. **(A)** A375 melanoma cells were transduced with Cas9 and non-targeting (NT) sgRNA or BIRC2-targeting sgRNAs and nuclear and cytoplasmic protein fractions were isolated and western blot analysis was performed. **(B)** Correlation of BIRC2 and NFKB2 expression in human melanoma tumors was assessed using cBioportal (Cerami et al., 2012; Gao et al., 2013). Data are representative of two **(A)** independent experiments.

**Figure S4, related to Figure 6.**
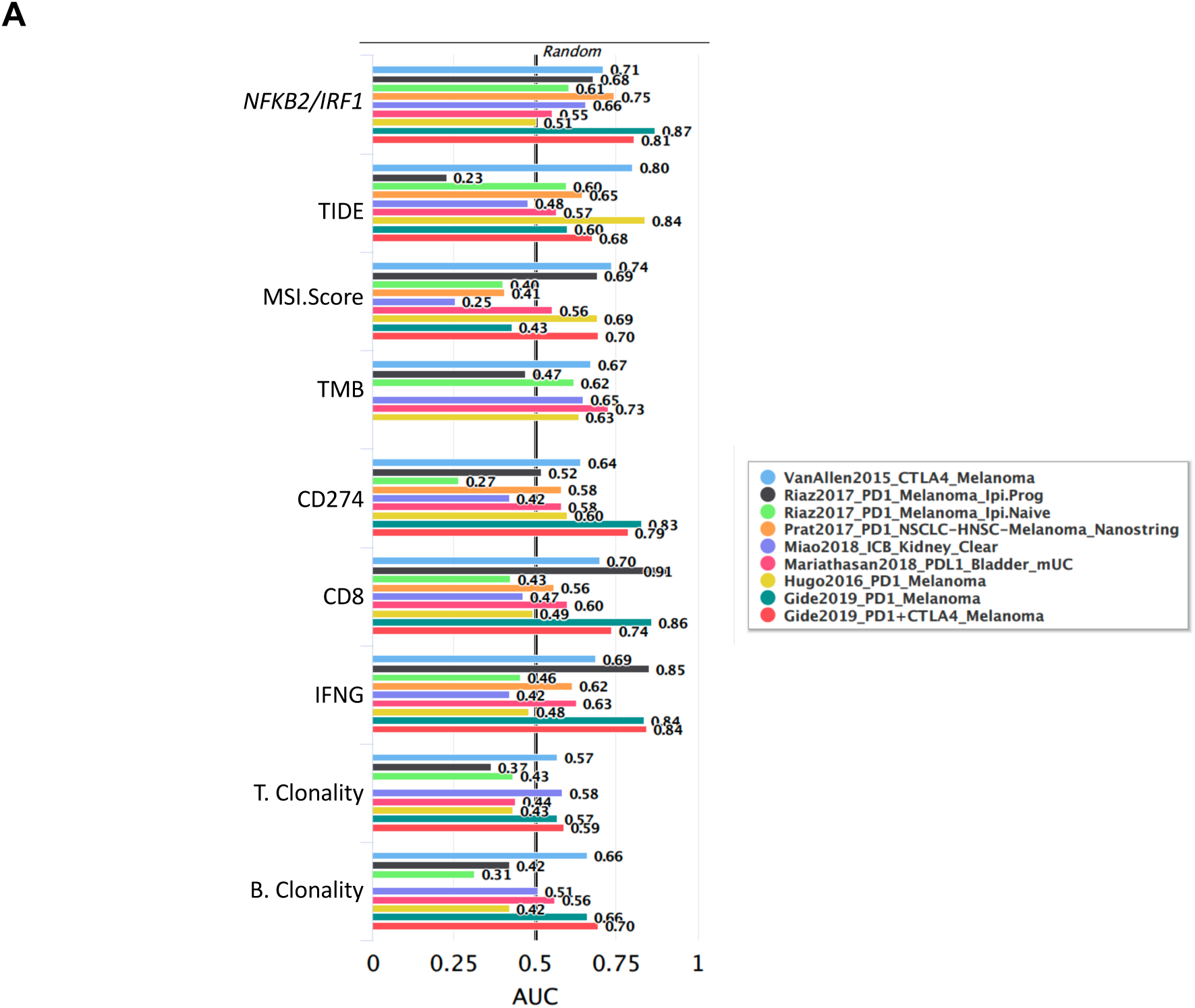
Expression of NFKB2 and IRF1 predicts response to immunotherapy. **(A)** Various biomarkers for response to immunotherapy (y axis), including expression of *NFKB2* + *IRF1* expression, were compared using previously published genesets with TIDE online platform (Jiang et al., 2018).

**Figure S5, related to Figure 7.**
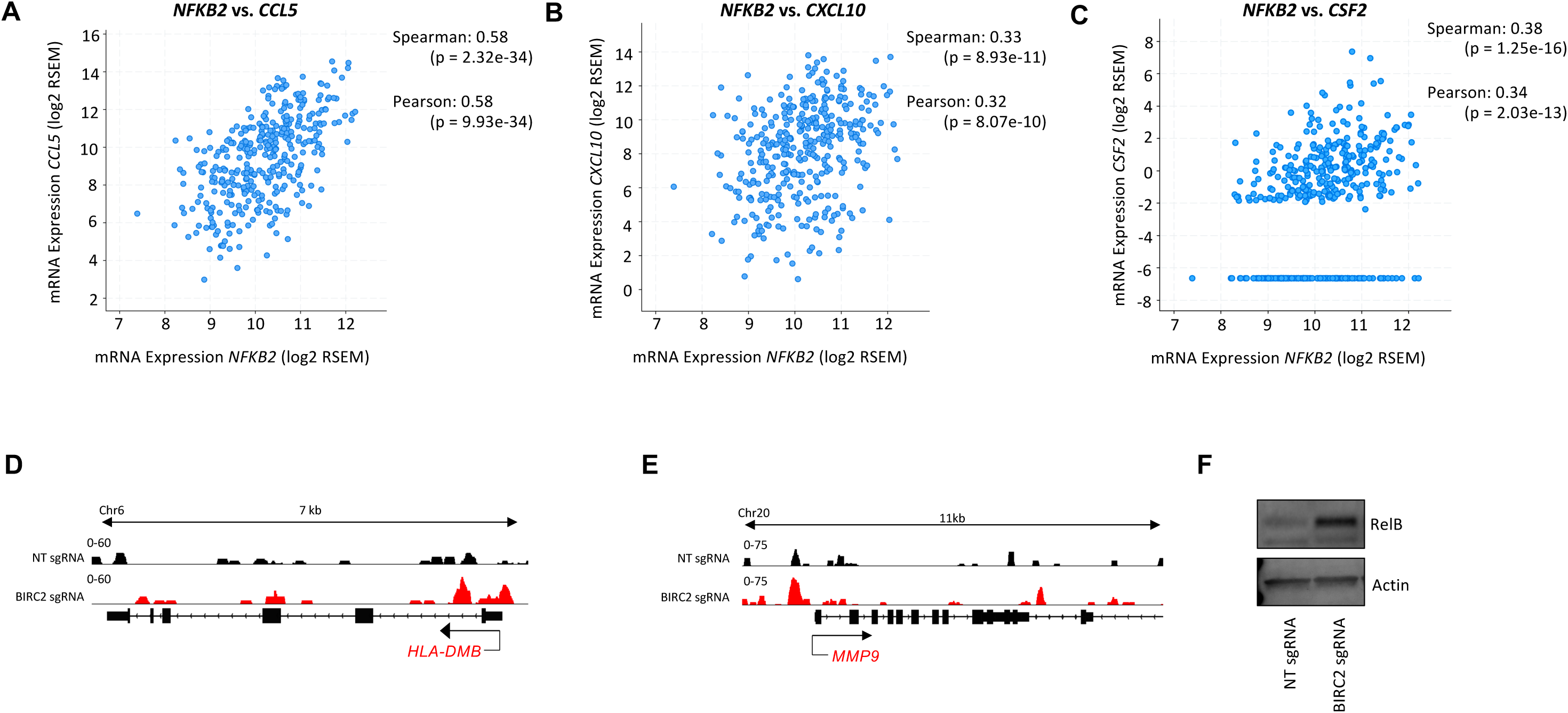
Non-canonical NF-κB signaling is associated with expression of inflammation-related genes. **(A-C)** Correlation of NFKB2 expression with **(A)** CCL5, **(B)** CXCL10 and **(C)** CSF2 expression in human melanoma tumors was assessed using cBioportal. **(D-E)** ChIPseq was performed on A375 cells transduced with Cas9 and non-targeting (NT) sgRNA or BIRC2-targeting sgRNA. **(D)** Association of RelB with HLA-C locus and **(E)** association of NFKB2 with MMP9 locus were assessed. **(F)** Western blot analysis of B16-mhGP100 melanoma cells transduced with Cas9 and non-targeting sgRNA or Cas9 and BIRC2-targeting sgRNA. Data are representative of two independent experiments **(D-F)**.

