## Supplemental Tables for "Genome-wide profiling of druggable active tumor defense mechanisms to enhance cancer immunotherapy"

### Table 1: Tumor genes identified as significantly depleted after genome 2CT screens

| Screen 1 | | | | Screen 2 | | |
|---|---|---|---|---|---|---|
| gene_id | riger | sigLabelSwitchFDR | | gene_id | riger | sigLabelSwitchFDR |
| hsa-mir-1299 | 101.5 | 0.00000 | | AHSG | 197 | 0.00000 |
| ALG11 | 319.75 | 0.00001 | | CHCHD10 | 237 | 0.00000 |
| BIRC2 | 327.75 | 0.00001 | | CBLL1 | 260.25 | 0.00000 |
| FAM32A | 342 | 0.00001 | | KRTAP19-7 | 362.75 | 0.00001 |
| KIF3C | 404.5 | 0.00001 | | PRSS16 | 424.25 | 0.00001 |
| LAPTM4A | 442.75 | 0.00002 | | GOLGA7B | 455 | 0.00001 |
| KRTAP3-2 | 490.5 | 0.00002 | | FAM32A | 480.25 | 0.00001 |
| ARSK | 498.25 | 0.00002 | | ARSD | 489.75 | 0.00001 |
| HHLA3 | 532.25 | 0.00002 | | hsa-mir-1299 | 490.5 | 0.00001 |
| LOC730183 | 543.5 | 0.00002 | | PGM3 | 493.5 | 0.00001 |
| LRR1 | 617.5 | 0.00002 | | APOBEC1 | 509.5 | 0.00001 |
| MED26 | 635.25 | 0.00002 | | SFR1 | 563.75 | 0.00002 |
| hsa-mir-197 | 654.5 | 0.00002 | | ENO4 | 662 | 0.00004 |
| TRAF2 | 703.25 | 0.00002 | | SLC6A18 | 664.75 | 0.00004 |
| EIF3I | 720.5 | 0.00002 | | CXADR | 685 | 0.00004 |
| RBM14 | 730.25 | 0.00003 | | ZFYVE28 | 689.25 | 0.00004 |
| hsa-mir-181b-2 | 782 | 0.00004 | | DNAJC16 | 694.25 | 0.00004 |
| VPS13D | 800.5 | 0.00004 | | SPDYA | 701.25 | 0.00004 |
| SCRT1 | 825.5 | 0.00004 | | hsa-mir-4435-1 | 764.75 | 0.00004 |
| HES4 | 830.25 | 0.00004 | | ARL2BP | 765.75 | 0.00004 |
| DPEP2 | 864.75 | 0.00005 | | HLCS | 769.25 | 0.00004 |
| MUC7 | 883.5 | 0.00005 | | CCDC62 | 780.5 | 0.00004 |
| ASPH | 894 | 0.00006 | | RAD21 | 799.5 | 0.00004 |
| EMC2 | 907.5 | 0.00006 | | hsa-mir-30d | 834.5 | 0.00005 |
| OSTM1 | 939.75 | 0.00006 | | CASKIN2 | 835.5 | 0.00005 |
| ZNF124 | 942.25 | 0.00006 | | EMID1 | 893.25 | 0.00006 |
| DCUN1D1 | 960.75 | 0.00007 | | TWIST2 | 905 | 0.00006 |
| TCEAL7 | 987 | 0.00007 | | SMC5 | 905.5 | 0.00006 |
| RRM1 | 989.75 | 0.00007 | | ECM1 | 944.25 | 0.00006 |
| SLC26A9 | 994.5 | 0.00007 | | APPL1 | 958.75 | 0.00007 |
| TCEB3CL | 997.75 | 0.00007 | | COL11A1 | 966 | 0.00007 |
| ZNF136 | 1038 | 0.00008 | | MRPS24 | 992.75 | 0.00007 |
| TMOD4 | 1038.5 | 0.00008 | | THEMIS2 | 998.75 | 0.00007 |
| PACRG | 1049.75 | 0.00008 | | FOSL2 | 1006.75 | 0.00008 |
| RALGAPA2 | 1078.75 | 0.00008 | | PUS10 | 1012.5 | 0.00008 |
| CLDND1 | 1095.25 | 0.00008 | | GPR12 | 1037.5 | 0.00008 |

| | | | | | | |
|---|---|---|---|---|---|---|
| PROL1 | 1099.25 | 0.00008 | | SCAF11 | 1046.25 | 0.00009 |
| TDO2 | 1133 | 0.00010 | | ISCU | 1049.5 | 0.00009 |
| MT1B | 1171.25 | 0.00010 | | EIF3I | 1052.25 | 0.00009 |
| COG3 | 1177.5 | 0.00011 | | POLR2B | 1069 | 0.00009 |
| CHMP7 | 1200 | 0.00012 | | ACSL1 | 1071.75 | 0.00009 |
| TMX4 | 1202 | 0.00012 | | ZNF232 | 1080.5 | 0.00009 |
| GIMD1 | 1202.75 | 0.00012 | | KAT5 | 1105 | 0.00009 |
| METTL14 | 1241.75 | 0.00012 | | ZGPAT | 1114.25 | 0.00009 |
| ORC1 | 1257 | 0.00013 | | ZBBX | 1123.5 | 0.00009 |
| CWC27 | 1264.25 | 0.00013 | | RLN3 | 1123.75 | 0.00009 |
| DPAGT1 | 1275 | 0.00013 | | PPCS | 1129.5 | 0.00010 |
| GPS2 | 1276.75 | 0.00013 | | TMIGD1 | 1132 | 0.00010 |
| SCEL | 1278.25 | 0.00013 | | RPL6 | 1153.75 | 0.00010 |
| ARMC3 | 1279.5 | 0.00013 | | ST8SIA6 | 1161 | 0.00010 |
| RBM10 | 1294.5 | 0.00013 | | UBE2N | 1177.25 | 0.00011 |
| hsa-mir-6775 | 1354.75 | 0.00016 | | hsa-mir-597 | 1196.25 | 0.00011 |
| RNF31 | 1355.75 | 0.00016 | | MUT | 1200 | 0.00012 |
| SCD5 | 1356 | 0.00016 | | MPG | 1200.75 | 0.00012 |
| MPL | 1376.75 | 0.00016 | | AHSA1 | 1249.25 | 0.00012 |
| NHSL1 | 1383 | 0.00016 | | TOR1AIP1 | 1279.5 | 0.00013 |
| CNGB1 | 1424.5 | 0.00017 | | hsa-mir-3652 | 1289 | 0.00013 |
| C22orf26 | 1425 | 0.00017 | | SNAP25 | 1295.5 | 0.00014 |
| CLEC16A | 1448.25 | 0.00018 | | ITSN2 | 1356.25 | 0.00016 |
| hsa-mir-1207 | 1453 | 0.00018 | | CCR8 | 1391.5 | 0.00016 |
| NLRC5 | 1456.25 | 0.00018 | | PIF1 | 1441 | 0.00017 |
| DDX21 | 1460.5 | 0.00018 | | TAS2R14 | 1451.5 | 0.00017 |
| FBXW7 | 1475 | 0.00018 | | OR2D3 | 1466.5 | 0.00018 |
| MOBP | 1507.25 | 0.00019 | | STK11IP | 1472.25 | 0.00018 |
| SART3 | 1513 | 0.00019 | | DHX35 | 1473 | 0.00018 |
| NKX1-2 | 1536.75 | 0.00019 | | SHOC2 | 1545 | 0.00020 |
| RPL23 | 1540.25 | 0.00019 | | FIGLA | 1589 | 0.00022 |
| YTHDF2 | 1591.5 | 0.00021 | | CRCP | 1605 | 0.00022 |
| hsa-mir-615 | 1592.25 | 0.00021 | | CYB5R3 | 1613.75 | 0.00023 |
| PRAMEF20 | 1604.25 | 0.00022 | | CAT | 1633.75 | 0.00023 |
| IL3 | 1604.75 | 0.00022 | | ATG13 | 1650.25 | 0.00024 |
| GABRR2 | 1620.75 | 0.00022 | | PHAX | 1673 | 0.00024 |
| OR5H14 | 1623 | 0.00022 | | GUF1 | 1685.25 | 0.00024 |
| CHST10 | 1632.25 | 0.00023 | | NDUFS8 | 1714.75 | 0.00025 |
| RPL13 | 1638.75 | 0.00023 | | NAPSA | 1737.5 | 0.00026 |

| | | | | | | |
|---|---|---|---|---|---|---|
| HIST2H2BE | 1643.75 | 0.00023 | | MX2 | 1744.75 | 0.00026 |
| THOP1 | 1649.25 | 0.00023 | | PRTG | 1750.25 | 0.00026 |
| DGAT2L6 | 1669.5 | 0.00024 | | OARD1 | 1754 | 0.00026 |
| KRTAP19-6 | 1697 | 0.00024 | | BTG3 | 1767.25 | 0.00026 |
| SUPT5H | 1704 | 0.00024 | | CHST3 | 1799.5 | 0.00028 |
| NLRP8 | 1706.25 | 0.00024 | | PPIA | 1847.75 | 0.00029 |
| OR4C11 | 1707 | 0.00024 | | SH2B3 | 1853 | 0.00029 |
| SLC25A24 | 1716.75 | 0.00024 | | HIF3A | 1860.25 | 0.00029 |
| VPS29 | 1720.5 | 0.00024 | | ATL2 | 1862 | 0.00029 |
| TMEM47 | 1734 | 0.00025 | | IL9 | 1871.25 | 0.00029 |
| LRRC4 | 1752.75 | 0.00025 | | DESI2 | 1886.5 | 0.00030 |
| ZNF227 | 1783.25 | 0.00026 | | CDC20 | 1886.75 | 0.00030 |
| FBXO36 | 1809.75 | 0.00028 | | ATG16L2 | 1887.5 | 0.00030 |
| GJC2 | 1815.5 | 0.00028 | | BTBD2 | 1890.25 | 0.00030 |
| GPD1L | 1816 | 0.00028 | | KIF21A | 1900.75 | 0.00031 |
| ZPLD1 | 1823 | 0.00028 | | NEURL2 | 1911.5 | 0.00031 |
| P4HA2 | 1825.25 | 0.00028 | | SLC44A1 | 1925 | 0.00031 |
| hsa-mir-4442 | 1852.5 | 0.00030 | | MRPL34 | 1993.5 | 0.00033 |
| ROS1 | 1873.25 | 0.00031 | | TMED2 | 2018 | 0.00034 |
| TTC30A | 1881.5 | 0.00031 | | PDK3 | 2062.25 | 0.00036 |
| POLRMT | 1887.75 | 0.00031 | | FMNL3 | 2076 | 0.00037 |
| RBL2 | 1891.25 | 0.00031 | | OXCT1 | 2084 | 0.00037 |
| SEH1L | 1909.25 | 0.00032 | | C8orf42 | 2101.25 | 0.00038 |
| DDX59 | 1914.75 | 0.00032 | | PCCB | 2124.25 | 0.00039 |
| SERTAD2 | 1929.75 | 0.00032 | | SPATA31A6 | 2126.25 | 0.00039 |
| ABI3BP | 1930.75 | 0.00032 | | NRCAM | 2135.5 | 0.00039 |
| DPT | 1978 | 0.00034 | | hsa-mir-519c | 2152.5 | 0.00040 |
| GRHPR | 2012 | 0.00035 | | GAR1 | 2155.75 | 0.00040 |
| KRTAP4-9 | 2017 | 0.00035 | | TMEM91 | 2179.25 | 0.00041 |
| LCN2 | 2017.75 | 0.00035 | | RBL2 | 2188 | 0.00042 |
| MEX3B | 2029.25 | 0.00035 | | HPS6 | 2228.75 | 0.00043 |
| RLTPR | 2030 | 0.00035 | | EGR2 | 2232.25 | 0.00044 |
| POLR3H | 2037.5 | 0.00035 | | CCNB1IP1 | 2233 | 0.00044 |
| RAD17 | 2048.5 | 0.00036 | | SAA2 | 2237 | 0.00044 |
| CCDC125 | 2081.75 | 0.00037 | | hsa-mir-3687 | 2256 | 0.00044 |
| UQCR11 | 2095.75 | 0.00037 | | ITGA7 | 2289 | 0.00046 |
| LCE2C | 2111.5 | 0.00038 | | DSTN | 2294.75 | 0.00046 |
| hsa-mir-3180-3 | 2158.75 | 0.00041 | | ENY2 | 2308 | 0.00046 |
| FAM3B | 2161.25 | 0.00041 | | PIP5KL1 | 2314.25 | 0.00046 |

| | | | | | | |
|---|---|---|---|---|---|---|
| DNMT3B | 2187.75 | 0.00042 | | VAMP8 | 2330.5 | 0.00047 |
| RPS25 | 2218 | 0.00043 | | KIF3A | 2340 | 0.00047 |
| TRIM43B | 2234.5 | 0.00044 | | ISX | 2344.5 | 0.00047 |
| IFIT1 | 2236.5 | 0.00044 | | GNL3L | 2344.5 | 0.00047 |
| PPP1R3F | 2250.5 | 0.00045 | | RASSF8 | 2367.5 | 0.00048 |
| SNRPD3 | 2270.75 | 0.00045 | | FMO4 | 2399.75 | 0.00050 |
| hsa-mir-6726 | 2315 | 0.00046 | | SYCE1 | 2414.25 | 0.00050 |
| hsa-mir-147a | 2321 | 0.00046 | | IL17D | 2421 | 0.00050 |
| TCEAL6 | 2325.25 | 0.00046 | | CINP | 2423.5 | 0.00050 |
| STRAP | 2334.75 | 0.00046 | | PRPF3 | 2483.5 | 0.00053 |
| DIDO1 | 2397 | 0.00049 | | FBXL13 | 2484 | 0.00053 |
| EIF4G1 | 2413.5 | 0.00050 | | EIF2A | 2484.5 | 0.00053 |
| C2orf49 | 2413.75 | 0.00050 | | ATP8B4 | 2501 | 0.00053 |
| SPRR1B | 2420.25 | 0.00050 | | ALG11 | 2518.5 | 0.00054 |
| TRAF3IP3 | 2462.25 | 0.00053 | | DDIT4L | 2525.5 | 0.00055 |
| C22orf23 | 2466 | 0.00053 | | GATA2 | 2529.75 | 0.00055 |
| TMEM184A | 2493 | 0.00053 | | IGSF9 | 2535 | 0.00056 |
| CDC123 | 2508.75 | 0.00053 | | hsa-mir-146b | 2553.5 | 0.00056 |
| B3GNT8 | 2523.25 | 0.00054 | | DPY19L4 | 2613.5 | 0.00059 |
| SLC6A6 | 2547.75 | 0.00055 | | C15orf61 | 2640.25 | 0.00059 |
| ADM5 | 2548 | 0.00055 | | CDCA8 | 2681.75 | 0.00061 |
| TRA2B | 2565.25 | 0.00056 | | DDX46 | 2689.25 | 0.00062 |
| ATF4 | 2577.75 | 0.00057 | | ZNF469 | 2705 | 0.00063 |
| SLC7A9 | 2590.75 | 0.00058 | | OR52E2 | 2712 | 0.00063 |
| hsa-mir-1289-1 | 2594.5 | 0.00058 | | NPAS2 | 2718.5 | 0.00064 |
| CYB5R3 | 2601.5 | 0.00058 | | P2RX4 | 2724.75 | 0.00064 |
| hsa-mir-4539 | 2623.5 | 0.00059 | | NFKBIL1 | 2780 | 0.00067 |
| SLC5A7 | 2652.75 | 0.00061 | | TLCD2 | 2836.75 | 0.00072 |
| RIOK1 | 2687.75 | 0.00064 | | LILRA4 | 2848 | 0.00072 |
| CCDC58 | 2705.75 | 0.00065 | | SRSF2 | 2902.5 | 0.00075 |
| hsa-mir-4456 | 2714.75 | 0.00066 | | MYH4 | 2909.75 | 0.00076 |
| FKBP6 | 2740.75 | 0.00068 | | KAL1 | 2955.75 | 0.00078 |
| COLEC11 | 2742 | 0.00068 | | ABHD3 | 2970 | 0.00079 |
| C9orf85 | 2752.75 | 0.00069 | | PGD | 2970.25 | 0.00079 |
| SUMO1 | 2791 | 0.00070 | | RELB | 3007.5 | 0.00081 |
| RAB17 | 2811.5 | 0.00071 | | PCNA | 3011.75 | 0.00081 |
| SNRPF | 2815.25 | 0.00071 | | RALB | 3021.25 | 0.00082 |
| PLK2 | 2840.5 | 0.00073 | | COL5A1 | 3035.5 | 0.00082 |
| EML3 | 2841.75 | 0.00073 | | HAL | 3037.25 | 0.00083 |

| | | | | | | |
|---|---|---|---|---|---|---|
| LPPR5 | 2845.25 | 0.00073 | | hsa-mir-4538 | 3042.25 | 0.00083 |
| RPAP2 | 2847.5 | 0.00073 | | OR1L8 | 3050.5 | 0.00083 |
| SLC9C2 | 2862.5 | 0.00074 | | NonTargetingControlGuideForHuman_0561 | 3078.25 | 0.00086 |
| CDC45 | 2877.5 | 0.00074 | | HDDC2 | 3099.5 | 0.00087 |
| hsa-mir-6868 | 2935 | 0.00076 | | hsa-mir-8060 | 3129.25 | 0.00088 |
| SFPQ | 2936.5 | 0.00076 | | IL6 | 3130 | 0.00089 |
| SLC5A9 | 2939.75 | 0.00076 | | PNMA3 | 3161.25 | 0.00090 |
| CHST13 | 2940.5 | 0.00076 | | LOC100507462 | 3239.25 | 0.00095 |
| VAV1 | 2996.25 | 0.00078 | | BIRC2 | 3240.25 | 0.00095 |
| hsa-mir-208a | 3002.25 | 0.00078 | | hsa-mir-1825 | 3241.25 | 0.00095 |
| ENY2 | 3013.75 | 0.00079 | | LRRFIP2 | 3243.75 | 0.00095 |
| GALR3 | 3025 | 0.00079 | | SBNO1 | 3288 | 0.00098 |
| MTA1 | 3028.25 | 0.00080 | | KCNG2 | 3312 | 0.00100 |
| DDX1 | 3049.75 | 0.00081 | | MYO1E | 3312.5 | 0.00100 |
| hsa-mir-4651 | 3050.25 | 0.00081 | | RQCD1 | 3319.25 | 0.00101 |
| TYW5 | 3051.25 | 0.00081 | | MADD | 3359.75 | 0.00104 |
| BAP1 | 3068.25 | 0.00082 | | KRTAP5-11 | 3373.25 | 0.00104 |
| ATP6V1C1 | 3100.75 | 0.00082 | | CAV1 | 3386.75 | 0.00106 |
| ALDH2 | 3119.75 | 0.00083 | | CCNA2 | 3397.75 | 0.00107 |
| hsa-mir-3129 | 3124.5 | 0.00083 | | REG1B | 3415.75 | 0.00108 |
| RRH | 3137.25 | 0.00085 | | IL1R1 | 3469.5 | 0.00111 |
| CHRM3 | 3142.25 | 0.00085 | | PDCL2 | 3473.5 | 0.00111 |
| SF1 | 3183.25 | 0.00087 | | OR1L6 | 3501.5 | 0.00113 |
| OCEL1 | 3211.75 | 0.00089 | | RPS26 | 3502.5 | 0.00113 |
| FFAR3 | 3216.5 | 0.00089 | | hsa-mir-4697 | 3526.75 | 0.00114 |
| ACTC1 | 3216.5 | 0.00089 | | WIBG | 3561 | 0.00117 |
| PPFIBP1 | 3246.25 | 0.00090 | | PSTK | 3562.75 | 0.00117 |
| hsa-mir-6794 | 3262.25 | 0.00091 | | TMED10 | 3564 | 0.00117 |
| MTMR14 | 3265.5 | 0.00091 | | CEBPG | 3593.25 | 0.00119 |
| INSL4 | 3293.5 | 0.00093 | | DEGS2 | 3611.25 | 0.00119 |
| CHRNA3 | 3298.75 | 0.00094 | | C12orf56 | 3620.5 | 0.00120 |
| hsa-mir-663b | 3299.25 | 0.00094 | | DLD | 3685 | 0.00124 |
| WARS2 | 3313.25 | 0.00094 | | TDGF1 | 3693.25 | 0.00124 |
| ATAD5 | 3325.75 | 0.00095 | | PIH1D3 | 3701 | 0.00124 |
| hsa-mir-5787 | 3331.5 | 0.00095 | | BUD31 | 3721.5 | 0.00126 |
| PSMC5 | 3336 | 0.00096 | | TRPV6 | 3721.5 | 0.00126 |
| LOC100507462 | 3336.25 | 0.00096 | | SPC25 | 3724.25 | 0.00126 |
| PRPF40A | 3356.5 | 0.00096 | | DNAJB8 | 3769 | 0.00128 |
| SLC2A8 | 3361.25 | 0.00097 | | NonTargetingControlGuideForHuman_0852 | 3828.25 | 0.00132 |

| | | | | | | |
|---|---|---|---|---|---|---|
| MRPL44 | 3365.25 | 0.00097 | | NonTargetingControlGuideForHuman_0766 | 3832 | 0.00132 |
| RPF1 | 3366.25 | 0.00097 | | KLHL31 | 3832.25 | 0.00132 |
| TTF2 | 3392.5 | 0.00099 | | hsa-mir-1289-1 | 3835 | 0.00133 |
| GLYAT | 3408.75 | 0.00099 | | PSD4 | 3837.5 | 0.00133 |
| ZNF69 | 3437 | 0.00102 | | CUL4A | 3874.25 | 0.00135 |
| BYSL | 3442.5 | 0.00103 | | LCK | 3878 | 0.00136 |
| KAL1 | 3448.25 | 0.00103 | | RLN1 | 3901.25 | 0.00138 |
| hsa-mir-3143 | 3456.75 | 0.00103 | | CSPG5 | 3931.5 | 0.00139 |
| PGS1 | 3471.5 | 0.00105 | | KRTAP5-10 | 3932.5 | 0.00140 |
| WBP2 | 3486.75 | 0.00106 | | hsa-mir-1251 | 3994 | 0.00144 |
| ZNF480 | 3494 | 0.00106 | | ARHGEF11 | 4001.25 | 0.00144 |
| PHYH | 3549.5 | 0.00109 | | ZMYND15 | 4046.5 | 0.00148 |
| COX5B | 3552 | 0.00109 | | AZI1 | 4049 | 0.00148 |
| PTMS | 3554.5 | 0.00109 | | PTPRCAP | 4089 | 0.00152 |
| EEF1B2 | 3558 | 0.00110 | | CCDC172 | 4108.25 | 0.00153 |
| MXD4 | 3580.5 | 0.00110 | | ZNF778 | 4117.75 | 0.00154 |
| HOXB8 | 3593.5 | 0.00111 | | LAMTOR3 | 4157 | 0.00158 |
| TMEM144 | 3609.75 | 0.00112 | | 2-Mar | 4158.5 | 0.00158 |
| SOX18 | 3612 | 0.00112 | | ISCA2 | 4167.5 | 0.00160 |
| RBM14-RBM4 | 3649.75 | 0.00114 | | POGLUT1 | 4169 | 0.00160 |
| hsa-mir-485 | 3674.75 | 0.00117 | | WBP11 | 4171 | 0.00160 |
| BRICD5 | 3702.25 | 0.00119 | | SYDE2 | 4239.75 | 0.00166 |
| PPP1R27 | 3747.75 | 0.00122 | | PDP1 | 4293.75 | 0.00170 |
| hsa-mir-6880 | 3803.25 | 0.00125 | | PGR | 4309.75 | 0.00171 |
| KRTAP4-6 | 3812.5 | 0.00126 | | LOC440563 | 4314.5 | 0.00172 |
| hsa-mir-4425 | 3813 | 0.00126 | | CHGA | 4316.25 | 0.00172 |
| AKIRIN1 | 3841.75 | 0.00127 | | ANKRD42 | 4331.25 | 0.00173 |
| FAM114A2 | 3845.5 | 0.00127 | | SHARPIN | 4346.75 | 0.00174 |
| IL36RN | 3861 | 0.00128 | | SRGAP2 | 4355.5 | 0.00174 |
| PRPF3 | 3902.25 | 0.00132 | | TMEM37 | 4364.75 | 0.00175 |
| hsa-mir-548w | 3941.75 | 0.00135 | | RPL12 | 4374 | 0.00176 |
| THAP5 | 3951.5 | 0.00136 | | ELTD1 | 4390.25 | 0.00176 |
| DPM2 | 3955.5 | 0.00136 | | MT3 | 4394.25 | 0.00176 |
| CDR2L | 3964.25 | 0.00136 | | GART | 4396.5 | 0.00177 |
| WDR45B | 4002.5 | 0.00137 | | TRIM37 | 4405.5 | 0.00177 |
| LYRM1 | 4044.5 | 0.00140 | | KRTAP5-1 | 4411.5 | 0.00178 |
| PTPLAD1 | 4053.5 | 0.00140 | | SNX1 | 4428.5 | 0.00180 |
| NLRC4 | 4055.75 | 0.00140 | | USP8 | 4429.75 | 0.00181 |
| OTOR | 4095.25 | 0.00143 | | KITLG | 4443.25 | 0.00181 |

| | | | | | | |
|---|---|---|---|---|---|---|
| DYDC1 | 4109.75 | 0.00145 | | FAM216A | 4447.75 | 0.00182 |
| BARHL2 | 4139.25 | 0.00147 | | CDH7 | 4468.25 | 0.00183 |
| hsa-mir-3197 | 4151.5 | 0.00148 | | EPB41 | 4487.25 | 0.00185 |
| PARP10 | 4173 | 0.00149 | | TOM1L1 | 4487.75 | 0.00185 |
| TNFRSF10B | 4226.5 | 0.00156 | | PIP5K1A | 4506.25 | 0.00187 |
| WFDC6 | 4236 | 0.00156 | | FOLR1 | 4523.75 | 0.00188 |
| FAM126A | 4238.5 | 0.00157 | | MRPL45 | 4524.75 | 0.00189 |
| TAF4 | 4252.75 | 0.00158 | | UQCRFS1 | 4551.25 | 0.00192 |
| TMIGD1 | 4256 | 0.00158 | | RPLP0 | 4567.25 | 0.00193 |
| hsa-mir-19a | 4283.25 | 0.00159 | | ADAMTS4 | 4570.75 | 0.00193 |
| LIN7A | 4283.5 | 0.00159 | | PDCD11 | 4659.5 | 0.00200 |
| POLR3F | 4311.5 | 0.00163 | | COPE | 4700.5 | 0.00202 |
| RNPS1 | 4325.75 | 0.00164 | | FAM50A | 4719.5 | 0.00203 |
| STUB1 | 4333.25 | 0.00165 | | DDX51 | 4785 | 0.00210 |
| CST4 | 4351.5 | 0.00167 | | FOXD4L5 | 4811.75 | 0.00214 |
| BCL2 | 4373 | 0.00168 | | NAALAD2 | 4818.25 | 0.00215 |
| POLD2 | 4428.5 | 0.00172 | | RPAP2 | 4818.75 | 0.00215 |
| PUS7 | 4470.75 | 0.00176 | | TAC1 | 4843.25 | 0.00217 |
| OR2B11 | 4497 | 0.00177 | | DAOA | 4852.75 | 0.00218 |
| COL7A1 | 4497.25 | 0.00177 | | hsa-mir-2276 | 4871.5 | 0.00219 |
| CCDC151 | 4498.75 | 0.00177 | | C14orf119 | 4877.25 | 0.00220 |
| HMGB4 | 4565.75 | 0.00182 | | NME8 | 4883.5 | 0.00221 |
| TOP1 | 4568.5 | 0.00182 | | TBCD | 4890.75 | 0.00221 |
| UBE2I | 4579.5 | 0.00183 | | DLGAP4 | 4962 | 0.00229 |
| ABCG8 | 4616.25 | 0.00185 | | hsa-mir-3185 | 4990.75 | 0.00231 |
| SLA2 | 4661.5 | 0.00189 | | ZNF350 | 5005.75 | 0.00232 |
| TREML1 | 4665.5 | 0.00190 | | PRRG1 | 5008.25 | 0.00232 |
| CHMP4B | 4666.5 | 0.00190 | | MAGEA3 | 5032 | 0.00233 |
| MAT2A | 4670 | 0.00190 | | MAL | 5051.75 | 0.00235 |
| ZBTB41 | 4678.75 | 0.00190 | | KISS1R | 5101.75 | 0.00239 |
| CASP6 | 4683.5 | 0.00191 | | MGA | 5164.5 | 0.00244 |
| METTL2B | 4721.75 | 0.00195 | | CELSR2 | 5180.25 | 0.00246 |
| RHO | 4734.5 | 0.00196 | | ERO1L | 5184.75 | 0.00246 |
| HOXC8 | 4744.25 | 0.00196 | | BCL2 | 5231.25 | 0.00250 |
| hsa-mir-320c-1 | 4750 | 0.00197 | | RPH3AL | 5243 | 0.00250 |
| RPS16 | 4755 | 0.00197 | | TXNDC16 | 5261 | 0.00252 |
| WSB2 | 4760.5 | 0.00198 | | SMIM8 | 5297 | 0.00256 |
| CINP | 4770.75 | 0.00198 | | hsa-mir-101-1 | 5315.5 | 0.00258 |
| SLC16A3 | 4771.5 | 0.00198 | | BRPF1 | 5317 | 0.00258 |

| | | | | | | |
|---|---|---|---|---|---|---|
| MS4A14 | 4801.25 | 0.00200 | | ICA1L | 5325.75 | 0.00259 |
| C1orf194 | 4818.5 | 0.00201 | | PRKRIP1 | 5331.5 | 0.00260 |
| CCDC93 | 4898.5 | 0.00207 | | hsa-mir-4690 | 5346 | 0.00262 |
| OR5H6 | 4912.5 | 0.00208 | | ROBO4 | 5349.25 | 0.00262 |
| STXBP4 | 4940 | 0.00210 | | FOCAD | 5426 | 0.00267 |
| CWF19L2 | 4942 | 0.00210 | | DSG3 | 5446 | 0.00269 |
| PHF8 | 4944 | 0.00210 | | CC2D1A | 5450.5 | 0.00270 |
| DDHD2 | 4951.25 | 0.00211 | | ZNF134 | 5455.25 | 0.00271 |
| ZNF665 | 4955.5 | 0.00212 | | ATP11B | 5460.75 | 0.00272 |
| MYRIP | 4958.25 | 0.00212 | | TMEM238 | 5515 | 0.00277 |
| ATP6V0C | 4971.25 | 0.00213 | | MARCKSL1 | 5533.25 | 0.00277 |
| MNS1 | 4990.5 | 0.00214 | | WNT7A | 5538.25 | 0.00278 |
| GATAD2A | 4994.75 | 0.00214 | | ZNF98 | 5589.5 | 0.00283 |
| C1orf111 | 5005.75 | 0.00216 | | HAX1 | 5599.25 | 0.00285 |
| hsa-mir-6777 | 5024.75 | 0.00217 | | PRKX | 5607.25 | 0.00285 |
| PNMAL1 | 5095 | 0.00223 | | SNX14 | 5691.75 | 0.00295 |
| OR1N2 | 5100 | 0.00224 | | ZNF607 | 5719.75 | 0.00298 |
| IL12A | 5131 | 0.00226 | | FAHD1 | 5781.5 | 0.00306 |
| KRT24 | 5163.5 | 0.00228 | | ZNF575 | 5787.5 | 0.00306 |
| CSF1 | 5193.75 | 0.00231 | | CFP | 5852.75 | 0.00312 |
| SMIM15 | 5199.25 | 0.00231 | | ARHGAP40 | 5876.25 | 0.00314 |
| RNF135 | 5200.5 | 0.00231 | | TSPYL6 | 5884.5 | 0.00314 |
| hsa-mir-142 | 5234.5 | 0.00234 | | INSIG2 | 5924 | 0.00317 |
| C19orf67 | 5252.75 | 0.00235 | | EFNB1 | 5935.5 | 0.00318 |
| NDNL2 | 5254.5 | 0.00235 | | CCSAP | 5977 | 0.00323 |
| PAK1IP1 | 5258.5 | 0.00235 | | PKP4 | 5998.5 | 0.00325 |
| RPL7 | 5262.25 | 0.00236 | | CNGA1 | 6005.75 | 0.00326 |
| GAGE10 | 5285.5 | 0.00237 | | RAG2 | 6087.75 | 0.00334 |
| IDE | 5287.75 | 0.00237 | | KCNJ9 | 6121.5 | 0.00337 |
| CCDC101 | 5310.25 | 0.00240 | | C9orf153 | 6127 | 0.00338 |
| TUBA3D | 5315.5 | 0.00240 | | EAF2 | 6142 | 0.00339 |
| DOK6 | 5356.25 | 0.00244 | | AUH | 6165.5 | 0.00343 |
| SPAST | 5369.25 | 0.00245 | | SLFN5 | 6166.25 | 0.00343 |
| ZNF175 | 5375.25 | 0.00245 | | RNF130 | 6192.5 | 0.00346 |
| OR5M1 | 5415.25 | 0.00249 | | FIZ1 | 6204 | 0.00348 |
| MYOCD | 5450.5 | 0.00254 | | HSCB | 6212.75 | 0.00349 |
| CETN2 | 5521 | 0.00260 | | SLC34A3 | 6227.5 | 0.00350 |
| NME6 | 5521.5 | 0.00260 | | EOGT | 6228.5 | 0.00351 |
| BUB1 | 5540.75 | 0.00261 | | LDHA | 6251 | 0.00352 |

| | | | | | | |
|---|---|---|---|---|---|---|
| hsa-mir-3923 | 5555 | 0.00263 | | ZC3H12C | 6257 | 0.00353 |
| FASN | 5593.25 | 0.00265 | | AKIRIN1 | 6262 | 0.00354 |
| PITHD1 | 5595 | 0.00265 | | IWS1 | 6288 | 0.00357 |
| RPS19 | 5622.25 | 0.00266 | | TRAF5 | 6356 | 0.00366 |
| SYNGR3 | 5687.25 | 0.00272 | | TMEM17 | 6371.5 | 0.00367 |
| hsa-mir-4421 | 5749.75 | 0.00278 | | ARFIP2 | 6400.5 | 0.00370 |
| PPAPDC1A | 5762.75 | 0.00280 | | TOMM7 | 6401.5 | 0.00370 |
| LRMP | 5770.25 | 0.00281 | | hsa-mir-3689d-2 | 6415.25 | 0.00372 |
| ECI1 | 5785.75 | 0.00282 | | PLEKHF1 | 6504.5 | 0.00382 |
| TCFL5 | 5790.25 | 0.00282 | | SENP5 | 6525 | 0.00384 |
| ECHDC3 | 5811.75 | 0.00285 | | MANSC1 | 6545 | 0.00386 |
| HCRTR1 | 5812.75 | 0.00285 | | SERTAD2 | 6550.75 | 0.00387 |
| IST1 | 5838 | 0.00289 | | MEX3D | 6564 | 0.00388 |
| hsa-mir-6820 | 5860.5 | 0.00291 | | CCDC42B | 6579.75 | 0.00389 |
| hsa-mir-4448 | 5884 | 0.00294 | | FAM104A | 6580 | 0.00389 |
| TAF10 | 5896.5 | 0.00295 | | POMP | 6623.75 | 0.00395 |
| ATP5H | 5975.75 | 0.00303 | | LPGAT1 | 6642 | 0.00397 |
| SLC27A5 | 5982.25 | 0.00304 | | SLC12A1 | 6649.5 | 0.00397 |
| TCP11X2 | 6038.75 | 0.00309 | | hsa-mir-6749 | 6670.75 | 0.00399 |
| SOX10 | 6047.75 | 0.00310 | | TBPL2 | 6676.75 | 0.00401 |
| SAC3D1 | 6055.75 | 0.00310 | | SC5D | 6690.5 | 0.00402 |
| TWF1 | 6063 | 0.00310 | | NGEF | 6695.5 | 0.00403 |
| DNAJC1 | 6099 | 0.00313 | | MED22 | 6701.5 | 0.00404 |
| CAPN13 | 6116.5 | 0.00315 | | MIDN | 6701.5 | 0.00404 |
| AKAP1 | 6151.75 | 0.00320 | | MPZL1 | 6706.25 | 0.00404 |
| CTGF | 6206.5 | 0.00326 | | WDR60 | 6738 | 0.00408 |
| SAP30BP | 6221.5 | 0.00328 | | FBXL15 | 6757.75 | 0.00409 |
| LBX1 | 6244.5 | 0.00329 | | VRK3 | 6780 | 0.00413 |
| SYBU | 6272.75 | 0.00332 | | ITGA10 | 6786.5 | 0.00413 |
| LALBA | 6309 | 0.00337 | | MIER1 | 6806 | 0.00416 |
| NCAPH2 | 6311.75 | 0.00338 | | DBI | 6812.75 | 0.00417 |
| PBX1 | 6315.75 | 0.00339 | | COG8 | 6826 | 0.00419 |
| HMGB1 | 6361 | 0.00344 | | OR13H1 | 6840 | 0.00419 |
| hsa-mir-1249 | 6398 | 0.00350 | | hsa-mir-5687 | 6869.5 | 0.00424 |
| hsa-mir-3173 | 6438.75 | 0.00353 | | RPL22L1 | 6886.5 | 0.00426 |
| GIPC2 | 6450 | 0.00354 | | KCNIP1 | 6887.75 | 0.00426 |
| TRIML1 | 6466.25 | 0.00357 | | COPA | 6897.75 | 0.00428 |
| YIPF6 | 6476.5 | 0.00358 | | ZNF549 | 6916.75 | 0.00430 |
| C9orf3 | 6513 | 0.00361 | | NKX2-2 | 7036 | 0.00448 |

| | | | | | | |
|---|---|---|---|---|---|---|
| hsa-mir-215 | 6532.75 | 0.00364 | | WDR45B | 7064.25 | 0.00450 |
| ATPAF1 | 6537.5 | 0.00365 | | CYP4F2 | 7078.25 | 0.00453 |
| hsa-mir-5591 | 6585.25 | 0.00371 | | hsa-mir-6165 | 7106.75 | 0.00456 |
| hsa-mir-4443 | 6595.25 | 0.00372 | | BSCL2 | 7107.25 | 0.00456 |
| ACADSB | 6607.5 | 0.00373 | | SLC25A25 | 7179.75 | 0.00464 |
| C11orf48 | 6650.75 | 0.00379 | | LARP6 | 7231.75 | 0.00471 |
| C11orf71 | 6699.25 | 0.00383 | | CTNND2 | 7241.25 | 0.00473 |
| hsa-mir-583 | 6735.75 | 0.00386 | | CAV2 | 7244.5 | 0.00473 |
| BAG3 | 6743 | 0.00387 | | PIGM | 7273.75 | 0.00475 |
| hsa-mir-661 | 6754.5 | 0.00388 | | ESD | 7284.25 | 0.00477 |
| TPM4 | 6768.75 | 0.00389 | | JAM2 | 7310.5 | 0.00479 |
| TMPRSS11D | 6801 | 0.00394 | | MECOM | 7321.25 | 0.00481 |
| C1orf185 | 6806.25 | 0.00394 | | GRIA2 | 7335.75 | 0.00482 |
| hsa-mir-6743 | 6824 | 0.00396 | | FAM63B | 7339.75 | 0.00482 |
| C16orf70 | 6841 | 0.00400 | | WDR18 | 7346.75 | 0.00483 |
| ZNF24 | 6843.25 | 0.00400 | | SRSF10 | 7362.75 | 0.00485 |
| ARHGAP24 | 6850.25 | 0.00400 | | BIK | 7364.75 | 0.00486 |
| HECW1 | 6869.75 | 0.00402 | | CAPN12 | 7384.75 | 0.00489 |
| SLC39A12 | 6894.75 | 0.00405 | | WDR44 | 7405.5 | 0.00492 |
| OSBPL3 | 6949 | 0.00411 | | C11orf54 | 7444.5 | 0.00496 |
| hsa-mir-3163 | 6961.75 | 0.00413 | | FARSB | 7446 | 0.00496 |
| PIGX | 6969.25 | 0.00414 | | hsa-mir-4419a | 7473 | 0.00501 |
| ERCC4 | 6982.75 | 0.00416 | | SEMA7A | 7488.75 | 0.00502 |
| PITPNM1 | 7029.25 | 0.00421 | | ZNF416 | 7600.5 | 0.00520 |
| MILR1 | 7114 | 0.00433 | | WNT6 | 7607.75 | 0.00520 |
| AAK1 | 7194 | 0.00442 | | NonTargetingControlGuideForHuman_0811 | 7615.75 | 0.00522 |
| PRSS55 | 7202 | 0.00444 | | ELOVL4 | 7634.75 | 0.00525 |
| MBP | 7217 | 0.00446 | | MEIG1 | 7648.75 | 0.00527 |
| TEX37 | 7221.75 | 0.00446 | | OLA1 | 7678.75 | 0.00529 |
| LARS2 | 7242.75 | 0.00447 | | HSBP1L1 | 7695 | 0.00531 |
| LYSMD4 | 7270.5 | 0.00449 | | ZMAT3 | 7709.25 | 0.00532 |
| NonTargetingControlGuideForHuman_0625 | 7285.25 | 0.00451 | | ATRNL1 | 7734.5 | 0.00536 |
| BIRC6 | 7287.5 | 0.00451 | | FAM3B | 7768.5 | 0.00540 |
| hsa-mir-378d-1 | 7300.75 | 0.00453 | | CLEC2B | 7777.75 | 0.00541 |
| F10 | 7317.25 | 0.00455 | | CCL4L2 | 7781.75 | 0.00542 |
| SPATA31A6 | 7334 | 0.00457 | | SFTA3 | 7789.5 | 0.00544 |
| FXYD2 | 7335 | 0.00457 | | ANXA1 | 7845 | 0.00550 |
| hsa-mir-744 | 7353.25 | 0.00460 | | PLK5 | 7871 | 0.00553 |
| EGR2 | 7373.25 | 0.00462 | | ARPP19 | 7877.75 | 0.00554 |

| | | | | | | |
|---|---|---|---|---|---|---|
| FRS2 | 7430.25 | 0.00470 | | ZNF556 | 7888.25 | 0.00555 |
| ZNF556 | 7474 | 0.00476 | | CHMP2B | 7932.5 | 0.00561 |
| CLMN | 7507 | 0.00481 | | ARHGAP20 | 7952.5 | 0.00564 |
| KLHL36 | 7512 | 0.00482 | | KBTBD13 | 8036 | 0.00577 |
| OR2F2 | 7551.25 | 0.00489 | | CREB3L2 | 8058.5 | 0.00581 |
| NAA25 | 7614 | 0.00498 | | ZNF654 | 8067 | 0.00582 |
| GRK7 | 7624.5 | 0.00500 | | SPACA1 | 8085 | 0.00584 |
| DOCK1 | 7626.5 | 0.00500 | | CXXC1 | 8101.75 | 0.00587 |
| USP37 | 7635.75 | 0.00500 | | ZNF322 | 8121.5 | 0.00590 |
| ZNF329 | 7637.5 | 0.00501 | | SKA3 | 8184.5 | 0.00599 |
| LDHB | 7638.25 | 0.00501 | | FSD1 | 8239.75 | 0.00605 |
| ARL4A | 7680.75 | 0.00506 | | TSG101 | 8264.5 | 0.00608 |
| DVL2 | 7756.75 | 0.00516 | | hsa-mir-346 | 8269.25 | 0.00608 |
| HMX1 | 7762.75 | 0.00517 | | SLC44A5 | 8294 | 0.00613 |
| RNF186 | 7781.25 | 0.00520 | | SERHL2 | 8295 | 0.00613 |
| hsa-mir-6845 | 7802.75 | 0.00523 | | DUS3L | 8301.75 | 0.00613 |
| NPPB | 7803.75 | 0.00524 | | HOXA11 | 8362.5 | 0.00621 |
| NonTargetingControlGuideForHuman_0582 | 7827.25 | 0.00526 | | PCMT1 | 8368.75 | 0.00622 |
| ADAM33 | 7846.25 | 0.00528 | | PTER | 8397.5 | 0.00626 |
| SETD9 | 7857.25 | 0.00529 | | XRCC2 | 8462.25 | 0.00639 |
| YPEL1 | 7864.25 | 0.00531 | | MOSPD2 | 8493.25 | 0.00642 |
| COPS8 | 7924.75 | 0.00538 | | SI | 8494.75 | 0.00642 |
| DYDC2 | 7957 | 0.00540 | | WIZ | 8511.5 | 0.00644 |
| RNF13 | 8064.5 | 0.00556 | | CISD3 | 8549.75 | 0.00649 |
| SLC25A25 | 8076.25 | 0.00558 | | DEFB125 | 8567.5 | 0.00651 |
| OTUD6A | 8087 | 0.00560 | | TMEM185B | 8571.5 | 0.00651 |
| ITGAV | 8093 | 0.00560 | | SLC23A3 | 8603.25 | 0.00655 |
| CNTFR | 8126 | 0.00564 | | TDG | 8653 | 0.00661 |
| SNRPD2 | 8135 | 0.00566 | | TSGA13 | 8655 | 0.00661 |
| NPS | 8148.5 | 0.00568 | | NDUFA3 | 8679.25 | 0.00664 |
| LAMTOR1 | 8189.5 | 0.00575 | | PRCP | 8751.25 | 0.00675 |
| CD200R1L | 8203.5 | 0.00576 | | ZDHHC16 | 8774 | 0.00679 |
| XPO4 | 8223.5 | 0.00578 | | AKTIP | 8792.75 | 0.00681 |
| AXIN1 | 8229.75 | 0.00580 | | SLC30A3 | 8840.25 | 0.00690 |
| PLA2G7 | 8273.75 | 0.00586 | | FSIP2 | 8859 | 0.00692 |
| RPP40 | 8311.5 | 0.00592 | | SP6 | 8909.5 | 0.00700 |
| KRTAP5-7 | 8392.25 | 0.00604 | | SRSF3 | 8910.25 | 0.00700 |
| CD160 | 8425.5 | 0.00608 | | CDC37 | 8921.75 | 0.00701 |
| SRPX2 | 8480 | 0.00616 | | HGD | 8922 | 0.00701 |

| | | | | | | |
|---|---|---|---|---|---|---|
| ZNF586 | 8492 | 0.00617 | | SLC7A6OS | 8946 | 0.00706 |
| HTATIP2 | 8501 | 0.00620 | | WFDC12 | 9044.75 | 0.00721 |
| SLC25A38 | 8548.25 | 0.00627 | | GPRC5B | 9050.75 | 0.00722 |
| CCDC66 | 8565.25 | 0.00630 | | ECM2 | 9091.75 | 0.00729 |
| TSR2 | 8594.25 | 0.00633 | | IL1R2 | 9118.75 | 0.00733 |
| ASGR1 | 8604.25 | 0.00634 | | FEN1 | 9131.25 | 0.00733 |
| MDN1 | 8667.25 | 0.00644 | | BAG5 | 9217 | 0.00745 |
| POLR3C | 8685 | 0.00644 | | MSANTD1 | 9222.5 | 0.00745 |
| GPHB5 | 8697.25 | 0.00646 | | GTF2H1 | 9223.25 | 0.00745 |
| TSPYL2 | 8714 | 0.00647 | | hsa-mir-4292 | 9233.75 | 0.00747 |
| PPA2 | 8740 | 0.00651 | | RWDD2B | 9247.5 | 0.00750 |
| NonTargetingControlGuideForHuman_0258 | 8741.5 | 0.00651 | | SLC25A28 | 9271.75 | 0.00754 |
| RNF5 | 8759.75 | 0.00654 | | MAPK13 | 9278.25 | 0.00755 |
| OR10J5 | 8809.5 | 0.00663 | | hsa-mir-2861 | 9335.75 | 0.00764 |
| hsa-mir-4697 | 8810.25 | 0.00663 | | APAF1 | 9339.75 | 0.00765 |
| TRAPPC2L | 8843.5 | 0.00670 | | OR2A14 | 9339.75 | 0.00765 |
| SPATS2 | 8851.5 | 0.00670 | | BOD1 | 9341 | 0.00765 |
| EPRS | 8874.75 | 0.00672 | | GLI1 | 9401 | 0.00774 |
| CTSK | 8886.5 | 0.00674 | | KRTAP5-3 | 9401.5 | 0.00774 |
| ISCU | 8926.25 | 0.00680 | | FKBPL | 9403.5 | 0.00775 |
| SLC30A6 | 8941.25 | 0.00681 | | TMEM57 | 9428.25 | 0.00778 |
| SEMA3E | 8986.5 | 0.00688 | | NPAT | 9437.5 | 0.00780 |
| GUCY1A3 | 9024.75 | 0.00695 | | NEK8 | 9494.25 | 0.00789 |
| NonTargetingControlGuideForHuman_0042 | 9069.5 | 0.00703 | | TROAP | 9505.75 | 0.00791 |
| hsa-mir-4755 | 9184.25 | 0.00722 | | RPS16 | 9529.5 | 0.00794 |
| OR52H1 | 9206 | 0.00727 | | NUDT8 | 9556 | 0.00799 |
| FAM206A | 9206.5 | 0.00727 | | CD300LD | 9559 | 0.00799 |
| APCDD1 | 9233.25 | 0.00732 | | C10orf99 | 9561.5 | 0.00800 |
| hsa-mir-3915 | 9233.5 | 0.00732 | | TXLNB | 9576 | 0.00803 |
| DNAJB5 | 9313.75 | 0.00744 | | TUBB4B | 9588.25 | 0.00805 |
| HENMT1 | 9345.75 | 0.00749 | | CRYBA4 | 9646.75 | 0.00816 |
| NUDC | 9406.5 | 0.00759 | | GIMAP2 | 9703 | 0.00824 |
| CHMP6 | 9427.25 | 0.00762 | | OR10J3 | 9765.5 | 0.00834 |
| MTMR6 | 9429 | 0.00763 | | CLDN2 | 9773.25 | 0.00835 |
| TACC1 | 9499.75 | 0.00772 | | SEBOX | 9781 | 0.00837 |
| KARS | 9516 | 0.00774 | | MTRNR2L2 | 9809.25 | 0.00843 |
| FTSJD1 | 9522.25 | 0.00775 | | C1orf86 | 9861.5 | 0.00853 |
| MEGF10 | 9523 | 0.00775 | | RNF180 | 9883.25 | 0.00856 |
| IGLON5 | 9556.5 | 0.00781 | | KIAA1429 | 9886.25 | 0.00856 |

| | | | | | | |
|---|---|---|---|---|---|---|
| ATP6V0A1 | 9624.25 | 0.00790 | | NUP85 | 9913.5 | 0.00860 |
| hsa-mir-5586 | 9640.75 | 0.00794 | | hsa-mir-1285-2 | 9915.75 | 0.00860 |
| MTRR | 9743.5 | 0.00810 | | USP25 | 9940.5 | 0.00865 |
| PLEKHM3 | 9757.75 | 0.00811 | | CCDC88A | 9941.5 | 0.00865 |
| TAS2R40 | 9764.25 | 0.00813 | | TBCA | 9942.25 | 0.00865 |
| CDK12 | 9773.25 | 0.00815 | | DNAJB6 | 10016 | 0.00881 |
| PPME1 | 9839 | 0.00828 | | CUL3 | 10043.75 | 0.00886 |
| GATAD2B | 9852.75 | 0.00830 | | ACTR10 | 10070.25 | 0.00889 |
| OTUD7B | 9864.25 | 0.00832 | | NonTargetingControlGuideForHuman_0491 | 10094 | 0.00893 |
| PPARD | 9867.25 | 0.00832 | | MTIF3 | 10096.75 | 0.00893 |
| EFCAB6 | 9869 | 0.00832 | | HDHD2 | 10115.75 | 0.00895 |
| TMEM127 | 9875.25 | 0.00834 | | LMAN1 | 10121 | 0.00896 |
| ANAPC16 | 9888.25 | 0.00836 | | CHUK | 10188 | 0.00908 |
| KRTAP9-9 | 9935.5 | 0.00847 | | DDO | 10205 | 0.00912 |
| PSMG1 | 9940.5 | 0.00848 | | ZNF726 | 10229 | 0.00917 |
| ARRDC1 | 9993.75 | 0.00858 | | SERPINB13 | 10256 | 0.00922 |
| hsa-mir-381 | 10037.75 | 0.00866 | | MYH14 | 10355.5 | 0.00941 |
| hsa-mir-501 | 10044.25 | 0.00867 | | ZHX2 | 10360.75 | 0.00942 |
| KDM2A | 10099.25 | 0.00875 | | MROH9 | 10371.5 | 0.00943 |
| RPL37 | 10103.5 | 0.00876 | | ZNF639 | 10392.25 | 0.00949 |
| STOX1 | 10109.75 | 0.00877 | | NR5A2 | 10413.75 | 0.00952 |
| PLAU | 10121.5 | 0.00879 | | FEZF1 | 10422.5 | 0.00954 |
| GADD45G | 10137.5 | 0.00883 | | BTG4 | 10429 | 0.00956 |
| SYF2 | 10152.75 | 0.00886 | | FN1 | 10461.25 | 0.00961 |
| TAS2R4 | 10236.25 | 0.00899 | | CKMT2 | 10488 | 0.00966 |
| XPO6 | 10241.25 | 0.00899 | | PARN | 10541.75 | 0.00977 |
| HOXB4 | 10264.75 | 0.00904 | | VDAC3 | 10548.25 | 0.00978 |
| CCNT1 | 10327 | 0.00916 | | ZNF595 | 10582 | 0.00984 |
| ITGA3 | 10410.25 | 0.00930 | | hsa-mir-150 | 10594.25 | 0.00985 |
| ZNF689 | 10438 | 0.00934 | | WDR37 | 10596.25 | 0.00986 |
| COL4A3BP | 10498.5 | 0.00945 | | RBP5 | 10711.75 | 0.01004 |
| ARGLU1 | 10498.75 | 0.00945 | | CKLF | 10734.5 | 0.01010 |
| NPM3 | 10539 | 0.00952 | | DHFR | 10814.25 | 0.01025 |
| PAH | 10543.75 | 0.00952 | | HERC2 | 10886.75 | 0.01037 |
| hsa-mir-4436b-2 | 10566 | 0.00956 | | TMEM216 | 10903 | 0.01040 |
| QSOX1 | 10572.75 | 0.00958 | | CHERP | 10936.75 | 0.01046 |
| CCDC127 | 10607.25 | 0.00965 | | DPM1 | 10940 | 0.01047 |

| | | | | | | |
|---|---|---|---|---|---|---|
| SPATA3 | 10630.75 | 0.00969 | | PKDREJ | 10948 | 0.01049 |
| F11R | 10657.5 | 0.00974 | | EN2 | 10958.5 | 0.01050 |
| MUC15 | 10696.25 | 0.00981 | | ZNF776 | 10960 | 0.01051 |
| hsa-mir-6884 | 10701.25 | 0.00982 | | SPRY1 | 10971 | 0.01053 |
| DDRGK1 | 10718.25 | 0.00985 | | KCTD14 | 11012.5 | 0.01061 |
| ST8SIA3 | 10731 | 0.00988 | | BCL3 | 11056 | 0.01068 |
| AR | 10738 | 0.00988 | | TNFSF12-TNFSF13 | 11104.75 | 0.01077 |
| VPS37A | 10811 | 0.01003 | | YIPF4 | 11110.25 | 0.01078 |
| CFH | 10813.25 | 0.01003 | | ZNF431 | 11112.75 | 0.01078 |
| ZRANB2 | 10842.25 | 0.01008 | | NIP7 | 11210.5 | 0.01093 |
| VSTM1 | 10858.25 | 0.01011 | | C3orf36 | 11239.75 | 0.01098 |
| PSG3 | 10873.5 | 0.01013 | | KRT13 | 11240.25 | 0.01098 |
| BEST3 | 10896.25 | 0.01017 | | NUDT13 | 11241.25 | 0.01098 |
| BANK1 | 10912.75 | 0.01020 | | KIAA1211L | 11259.5 | 0.01102 |
| TAL1 | 10915.75 | 0.01021 | | KCNT2 | 11269.25 | 0.01103 |
| PET117 | 10943.5 | 0.01026 | | FEV | 11296.5 | 0.01109 |
| COMMD7 | 10988 | 0.01034 | | TIMP1 | 11324.75 | 0.01115 |
| PVR | 11021 | 0.01039 | | DGKE | 11376.75 | 0.01123 |
| LOC284385 | 11040.25 | 0.01042 | | CHRNG | 11430.5 | 0.01134 |
| hsa-mir-6127 | 11047.25 | 0.01044 | | ACSM4 | 11439.5 | 0.01136 |
| ATP6AP1 | 11097.75 | 0.01053 | | POLR1A | 11486.75 | 0.01145 |
| RPL30 | 11114.75 | 0.01058 | | C1orf172 | 11486.75 | 0.01145 |
| GRIA2 | 11127.25 | 0.01060 | | CLCC1 | 11501.75 | 0.01149 |
| SGCB | 11152.25 | 0.01065 | | IQCF3 | 11507.75 | 0.01150 |
| TMTC1 | 11199.5 | 0.01075 | | MBOAT4 | 11512.75 | 0.01151 |
| SKA2 | 11218.25 | 0.01078 | | ATF5 | 11518.75 | 0.01152 |
| RPLP2 | 11239.75 | 0.01083 | | FSTL4 | 11552.25 | 0.01158 |
| BCAS1 | 11293.25 | 0.01094 | | TMX4 | 11624.5 | 0.01173 |
| ACSM2A | 11302.5 | 0.01096 | | ANKS4B | 11731.25 | 0.01194 |
| CENPF | 11306.5 | 0.01097 | | TRIM52 | 11736.5 | 0.01194 |
| ESD | 11331.75 | 0.01103 | | GABRQ | 11738.75 | 0.01194 |
| THRAP3 | 11344 | 0.01105 | | HRH4 | 11755.5 | 0.01197 |
| GARNL3 | 11375 | 0.01111 | | LOC100506422 | 11784 | 0.01203 |
| EAF2 | 11511.75 | 0.01136 | | LMOD3 | 11801.5 | 0.01208 |
| NDUFA8 | 11542.25 | 0.01141 | | COG3 | 11887.25 | 0.01225 |
| CD180 | 11546.75 | 0.01141 | | PPP2R5D | 11930 | 0.01233 |
| KBTBD3 | 11560.5 | 0.01145 | | SAE1 | 11937.5 | 0.01234 |

| | | | | | | |
|---|---|---|---|---|---|---|
| CLCN7 | 11566.5 | 0.01146 | | CLDN10 | 11939.5 | 0.01235 |
| PAX8 | 11585.75 | 0.01150 | | SF3B4 | 11948.75 | 0.01238 |
| SLC16A9 | 11590 | 0.01152 | | CFHR5 | 11960.25 | 0.01241 |
| hsa-mir-6863 | 11603.75 | 0.01155 | | NGB | 12013.5 | 0.01251 |
| UBE2V1 | 11612.25 | 0.01156 | | TK2 | 12047.25 | 0.01258 |
| CRYBB3 | 11619.25 | 0.01157 | | TMEM69 | 12099.25 | 0.01271 |
| DZIP1 | 11676.25 | 0.01168 | | PDE1B | 12113.5 | 0.01273 |
| NXPE3 | 11694.5 | 0.01171 | | STRN4 | 12242.75 | 0.01302 |
| MINA | 11719 | 0.01177 | | ADARB2 | 12270.25 | 0.01309 |
| GDF10 | 11774.25 | 0.01191 | | KRTAP9-4 | 12343.75 | 0.01326 |
| SPATA31A4 | 11804 | 0.01195 | | HDGF | 12365.75 | 0.01332 |
| DDX24 | 11812.5 | 0.01197 | | RAB11FIP1 | 12380.5 | 0.01335 |
| TMEM251 | 11831 | 0.01201 | | NR2F6 | 12403 | 0.01341 |
| hsa-mir-653 | 11900.75 | 0.01219 | | SCGB1D1 | 12410 | 0.01342 |
| HKDC1 | 11949.25 | 0.01229 | | LOC158434 | 12432 | 0.01345 |
| PPP2CA | 11952.25 | 0.01229 | | CHST1 | 12441 | 0.01346 |
| ATP1B4 | 11966.5 | 0.01233 | | GDF11 | 12442.75 | 0.01346 |
| NEXN | 11981.5 | 0.01237 | | HHLA3 | 12467.25 | 0.01351 |
| CA7 | 12013.75 | 0.01242 | | SRP54 | 12468.25 | 0.01352 |
| AAMP | 12038.5 | 0.01248 | | CWC25 | 12570.75 | 0.01370 |
| C12orf71 | 12204.75 | 0.01281 | | COX6C | 12604.5 | 0.01377 |
| CDC23 | 12222.25 | 0.01285 | | ARHGAP19 | 12607.25 | 0.01379 |
| CTSO | 12283.5 | 0.01300 | | HSD11B2 | 12617.75 | 0.01381 |
| HGD | 12288.25 | 0.01301 | | AFMID | 12631.75 | 0.01383 |
| hsa-mir-6516 | 12308.5 | 0.01306 | | ACTR1A | 12672.25 | 0.01390 |
| FITM1 | 12312.25 | 0.01307 | | IQCH | 12701.25 | 0.01396 |
| CRISP1 | 12326 | 0.01309 | | CSNK1G1 | 12743.75 | 0.01403 |
| GCSAML | 12333.25 | 0.01311 | | SLC12A2 | 12843.5 | 0.01429 |
| PDDC1 | 12344.75 | 0.01312 | | DUT | 12849.25 | 0.01431 |
| VSIG8 | 12354.5 | 0.01314 | | hsa-mir-4780 | 12850.25 | 0.01431 |
| GLTSCR1 | 12370.75 | 0.01317 | | hsa-mir-3907 | 12905.5 | 0.01446 |
| DLG1 | 12372.25 | 0.01317 | | KPNA3 | 12907.75 | 0.01446 |
| CCL24 | 12380.5 | 0.01319 | | TBX20 | 12916.75 | 0.01449 |
| REN | 12400.5 | 0.01322 | | CFH | 12922.25 | 0.01450 |
| hsa-mir-548a-1 | 12470.75 | 0.01337 | | CDK3 | 12949.75 | 0.01456 |
| hsa-mir-4297 | 12478 | 0.01339 | | COL10A1 | 12974.25 | 0.01463 |

| | | | | | | |
|---|---|---|---|---|---|---|
| ADCY2 | 12520.25 | 0.01350 | | TMEM27 | 12997.25 | 0.01470 |
| HRASLS2 | 12522.75 | 0.01350 | | OCIAD1 | 13001.25 | 0.01471 |
| hsa-mir-3908 | 12532.5 | 0.01353 | | SERTAD1 | 13012.75 | 0.01473 |
| MANSC1 | 12550.25 | 0.01355 | | INTS7 | 13023 | 0.01476 |
| PDF | 12551.5 | 0.01356 | | TOP3A | 13109.75 | 0.01496 |
| POM121C | 12560.75 | 0.01359 | | KRT28 | 13123 | 0.01499 |
| BTBD17 | 12584.5 | 0.01363 | | TMEM72 | 13151.25 | 0.01506 |
| NUDT11 | 12616.25 | 0.01373 | | ANXA6 | 13218.75 | 0.01522 |
| IL13RA1 | 12640.75 | 0.01377 | | UNC45B | 13242 | 0.01527 |
| CCDC74A | 12680.5 | 0.01385 | | HLA-DRB1 | 13259.25 | 0.01531 |
| GPATCH1 | 12723.5 | 0.01395 | | MSL1 | 13319.5 | 0.01547 |
| PSMD6 | 12753.25 | 0.01405 | | ACTA2 | 13322.25 | 0.01548 |
| MYOC | 12758.75 | 0.01406 | | ANO7 | 13326.25 | 0.01549 |
| MOB1A | 12793.75 | 0.01413 | | CXCL10 | 13376.25 | 0.01560 |
| PCDHGB3 | 12818.75 | 0.01419 | | TNFRSF9 | 13417.75 | 0.01571 |
| ATP8A1 | 12842 | 0.01426 | | FIBIN | 13423.5 | 0.01573 |
| hsa-mir-600 | 12843.75 | 0.01426 | | CHIA | 13435 | 0.01576 |
| SCAF11 | 12852.25 | 0.01428 | | CRADD | 13440.5 | 0.01578 |
| ZFP90 | 12874.5 | 0.01433 | | KRT33A | 13442.75 | 0.01578 |
| MRPL35 | 12881.75 | 0.01435 | | DKC1 | 13462.5 | 0.01581 |
| hsa-mir-4499 | 12931.5 | 0.01446 | | NonTargetingControlGuideForHuman_0147 | 13495.5 | 0.01588 |
| ABCA8 | 12939.5 | 0.01449 | | CDHR5 | 13510.5 | 0.01592 |
| SOX14 | 12949.25 | 0.01451 | | KCNAB2 | 13528.75 | 0.01598 |
| FBXL15 | 12965.75 | 0.01456 | | HSF2 | 13539.25 | 0.01600 |
| IFI27L2 | 13067 | 0.01480 | | CBX4 | 13586.5 | 0.01612 |
| SRFBP1 | 13082.75 | 0.01484 | | hsa-mir-378c | 13593.75 | 0.01614 |
| FAM76B | 13099.75 | 0.01487 | | KRTAP9-9 | 13602.75 | 0.01616 |
| TMIE | 13137 | 0.01496 | | DMRTB1 | 13615.75 | 0.01620 |
| FKBP4 | 13192.75 | 0.01509 | | GK5 | 13643 | 0.01626 |
| CSRP2BP | 13199.5 | 0.01511 | | ODF2L | 13668.75 | 0.01630 |
| TNRC6C | 13202 | 0.01512 | | EIF3B | 13674.75 | 0.01631 |
| RPL24 | 13319 | 0.01539 | | hsa-mir-23a | 13677 | 0.01631 |
| VASP | 13340.75 | 0.01542 | | SNPH | 13679.25 | 0.01632 |
| RAI2 | 13345.25 | 0.01543 | | hsa-mir-8076 | 13692.75 | 0.01634 |
| ARSD | 13357.75 | 0.01546 | | HARS2 | 13694.25 | 0.01634 |
| RPS8 | 13362.5 | 0.01547 | | POLI | 13742.5 | 0.01645 |
| hsa-mir-548s | 13389.5 | 0.01553 | | C7orf73 | 13753 | 0.01646 |

| | | | | | | |
|---|---|---|---|---|---|---|
| PINLYP | 13390.75 | 0.01553 | | YLPM1 | 13862.25 | 0.01673 |
| FTMT | 13394.25 | 0.01554 | | FGF23 | 13876.75 | 0.01676 |
| KIAA1211L | 13443.25 | 0.01564 | | TAS2R46 | 13877 | 0.01676 |
| ATG4A | 13488 | 0.01573 | | MFN2 | 13923.75 | 0.01689 |
| SDAD1 | 13511.25 | 0.01579 | | TOX2 | 13955 | 0.01697 |
| FAM131A | 13607.75 | 0.01599 | | PJA2 | 13958.75 | 0.01698 |
| COPS6 | 13624 | 0.01601 | | SART3 | 13976.25 | 0.01701 |
| hsa-mir-1277 | 13646.5 | 0.01607 | | SOX21 | 13981.25 | 0.01703 |
| LOC100996485 | 13674.5 | 0.01611 | | TCEB1 | 13992.75 | 0.01704 |
| PPP2R5D | 13705 | 0.01619 | | hsa-mir-4686 | 14031.75 | 0.01714 |
| PSMB3 | 13829.25 | 0.01648 | | CCDC92 | 14062.25 | 0.01721 |
| SNX1 | 13864.25 | 0.01655 | | TRIM27 | 14086.75 | 0.01727 |
| TXNL4A | 13873 | 0.01657 | | FTSJD1 | 14107.75 | 0.01732 |
| ROPN1 | 13901.75 | 0.01665 | | OSBPL8 | 14135.75 | 0.01741 |
| C14orf119 | 13958.25 | 0.01679 | | OR52I1 | 14144 | 0.01743 |
| C15orf61 | 13996.75 | 0.01689 | | NUDC | 14149.5 | 0.01745 |
| ADAMTS5 | 14040.25 | 0.01699 | | DKK1 | 14185 | 0.01754 |
| HIBADH | 14051.5 | 0.01701 | | BLNK | 14220.75 | 0.01763 |
| SLITRK2 | 14061.5 | 0.01704 | | CCDC71L | 14252.25 | 0.01771 |
| CLDN5 | 14076.5 | 0.01707 | | FAM46C | 14256.75 | 0.01772 |
| C17orf97 | 14161.75 | 0.01726 | | TMCO5A | 14263.25 | 0.01774 |
| DNAJB3 | 14172.75 | 0.01728 | | RPS6KA2 | 14264.5 | 0.01774 |
| AMER3 | 14246.25 | 0.01745 | | SLC26A1 | 14321.5 | 0.01789 |
| PRPF4B | 14255.5 | 0.01748 | | MYBL2 | 14353.5 | 0.01795 |
| AK5 | 14265.5 | 0.01750 | | C7orf61 | 14423.5 | 0.01812 |
| IDS | 14287.5 | 0.01756 | | RPS2 | 14424 | 0.01812 |
| MAOB | 14307.25 | 0.01761 | | HPS3 | 14496.5 | 0.01832 |
| ALKBH5 | 14326.5 | 0.01764 | | CYP17A1 | 14505.75 | 0.01834 |
| STAT1 | 14345.75 | 0.01769 | | XAGE2 | 14605.25 | 0.01856 |
| hsa-mir-6727 | 14362.25 | 0.01773 | | hsa-mir-198 | 14621 | 0.01859 |
| IPMK | 14366.5 | 0.01775 | | OR2T5 | 14634.25 | 0.01864 |
| YBX3 | 14375.75 | 0.01777 | | RBM20 | 14647.5 | 0.01869 |
| PLCL1 | 14419 | 0.01790 | | GULP1 | 14681.25 | 0.01876 |
| hsa-mir-4280 | 14443 | 0.01796 | | SETX | 14708.25 | 0.01881 |
| CSNK2B | 14446.5 | 0.01797 | | hsa-mir-18b | 14708.75 | 0.01881 |
| DHDH | 14462 | 0.01799 | | CLGN | 14768 | 0.01896 |
| ATP10B | 14467.75 | 0.01800 | | TBK1 | 14789.5 | 0.01901 |

| | | | | | | |
|---|---|---|---|---|---|---|
| MTX1 | 14483.25 | 0.01804 | | CACTIN | 14830.25 | 0.01910 |
| BPIFA2 | 14518.5 | 0.01812 | | ZNF280C | 14876.25 | 0.01922 |
| hsa-mir-8086 | 14546 | 0.01820 | | PRDX1 | 14937.5 | 0.01941 |
| TEAD2 | 14561.25 | 0.01824 | | ARHGEF5 | 14966.75 | 0.01949 |
| NKX2-1 | 14584.5 | 0.01831 | | TMC1 | 15052.5 | 0.01974 |
| GPR110 | 14605.5 | 0.01837 | | POLM | 15073.25 | 0.01979 |
| GCGR | 14606 | 0.01837 | | SH3BP1 | 15080.5 | 0.01981 |
| hsa-mir-580 | 14618.5 | 0.01841 | | PHF20 | 15129 | 0.01994 |
| CSTF2 | 14660.5 | 0.01850 | | DPM2 | 15139.75 | 0.01997 |
| ADD3 | 14682.75 | 0.01857 | | SCGB1D2 | 15154 | 0.02000 |
| OTUD4 | 14688.5 | 0.01858 | | FXN | 15175.75 | 0.02007 |
| ZNF365 | 14719 | 0.01869 | | WDR92 | 15196 | 0.02013 |
| LPGAT1 | 14738 | 0.01872 | | KIAA0825 | 15242.5 | 0.02026 |
| FOLR3 | 14776.5 | 0.01883 | | CD300A | 15347.5 | 0.02050 |
| CNKSR3 | 14785.5 | 0.01885 | | GBX1 | 15399 | 0.02063 |
| AKT1S1 | 14846 | 0.01901 | | hsa-mir-621 | 15408.25 | 0.02065 |
| GUCA2A | 14859.25 | 0.01904 | | NCF2 | 15464.25 | 0.02078 |
| CD5L | 14946 | 0.01927 | | CDH11 | 15609 | 0.02118 |
| DOCK5 | 14948.75 | 0.01928 | | TMEM52 | 15644.5 | 0.02130 |
| NTF3 | 14982.25 | 0.01939 | | DPH2 | 15651.5 | 0.02131 |
| NUDCD2 | 14989.5 | 0.01940 | | DIRAS2 | 15651.5 | 0.02131 |
| ZNF202 | 15075 | 0.01965 | | THAP1 | 15660.25 | 0.02133 |
| C9orf153 | 15078.25 | 0.01966 | | POLR2E | 15702.75 | 0.02144 |
| MC5R | 15136.5 | 0.01980 | | C16orf97 | 15702.75 | 0.02144 |
| OR10A3 | 15232 | 0.02006 | | SMCHD1 | 15704.75 | 0.02144 |
| MRPL1 | 15247.75 | 0.02009 | | SLC4A8 | 15764.25 | 0.02160 |
| CLEC4E | 15261 | 0.02012 | | STX17 | 15817.75 | 0.02176 |
| ATF3 | 15262 | 0.02013 | | HEATR5B | 15858.75 | 0.02188 |
| hsa-mir-5096 | 15378.25 | 0.02047 | | RPS23 | 15859 | 0.02188 |
| MAB21L1 | 15415.75 | 0.02058 | | PTN | 15879.75 | 0.02194 |
| EPS8 | 15417.25 | 0.02058 | | NDUFAF6 | 15907 | 0.02201 |
| CHSY1 | 15429.25 | 0.02061 | | MYLK4 | 15914.5 | 0.02203 |
| CATSPERG | 15454.5 | 0.02068 | | CDC42EP3 | 15944.25 | 0.02211 |
| PYROXD2 | 15457 | 0.02069 | | TFCP2L1 | 15975.75 | 0.02220 |
| MYADML2 | 15459 | 0.02069 | | NDUFC1 | 16023 | 0.02234 |
| hsa-mir-6129 | 15514.5 | 0.02084 | | MEMO1 | 16064.5 | 0.02249 |
| USP7 | 15520.75 | 0.02086 | | RBM48 | 16072 | 0.02250 |

| | | | | | | |
|---|---|---|---|---|---|---|
| OTC | 15526.5 | 0.02088 | | BCCIP | 16093.5 | 0.02255 |
| NOC2L | 15607.5 | 0.02111 | | EXOSC8 | 16110.75 | 0.02260 |
| C17orf80 | 15619 | 0.02114 | | NonTargetingControlGuideForHuman_0717 | 16135 | 0.02266 |
| TIMM21 | 15672.25 | 0.02130 | | OR8U1 | 16144.75 | 0.02268 |
| IFNA8 | 15761.75 | 0.02158 | | EAF1 | 16212.75 | 0.02292 |
| ALDH3B1 | 15820 | 0.02174 | | MOBP | 16275.25 | 0.02307 |
| OR7A10 | 15863.5 | 0.02187 | | GABRE | 16283.25 | 0.02310 |
| NT5E | 15869.75 | 0.02188 | | TPM2 | 16300 | 0.02315 |
| CNOT3 | 15913.75 | 0.02198 | | THRAP3 | 16373 | 0.02336 |
| SIX5 | 15920.25 | 0.02200 | | TMF1 | 16388 | 0.02338 |
| CALCR | 15927 | 0.02201 | | MDH1 | 16480.25 | 0.02367 |
| NAALAD2 | 15936 | 0.02204 | | FLYWCH1 | 16509.25 | 0.02374 |
| ZNF430 | 16020.5 | 0.02226 | | KCNJ4 | 16571.5 | 0.02389 |
| hsa-mir-626 | 16027.75 | 0.02229 | | TMPRSS11D | 16625 | 0.02402 |
| ZNF225 | 16055 | 0.02235 | | CCL18 | 16635.5 | 0.02407 |
| CELA3A | 16067.75 | 0.02239 | | DEDD2 | 16747 | 0.02436 |
| C11orf35 | 16087.25 | 0.02243 | | PDK1 | 16774.25 | 0.02444 |
| PRKAR1A | 16119.5 | 0.02251 | | ZNF704 | 16806 | 0.02454 |
| POLR2K | 16126 | 0.02253 | | PIK3R6 | 16860.75 | 0.02470 |
| ULK4 | 16134.25 | 0.02255 | | hsa-mir-633 | 16869 | 0.02472 |
| C5orf47 | 16135.75 | 0.02255 | | ENDOD1 | 16873.25 | 0.02473 |
| ZFP36 | 16140.75 | 0.02257 | | hsa-mir-4766 | 16890 | 0.02478 |
| PROKR1 | 16151.75 | 0.02259 | | PXK | 16920 | 0.02488 |
| EMCN | 16162 | 0.02261 | | FAM178A | 16954.5 | 0.02500 |
| MTRNR2L5 | 16171.75 | 0.02266 | | ABCC4 | 16970 | 0.02506 |
| NFX1 | 16191.5 | 0.02272 | | NADK | 16982 | 0.02509 |
| OR10J1 | 16223.5 | 0.02280 | | CPEB3 | 16983 | 0.02509 |
| CAPNS2 | 16326.5 | 0.02313 | | MAGEH1 | 16986.25 | 0.02510 |
| EPHA6 | 16337 | 0.02316 | | ALG10B | 16987.25 | 0.02510 |
| OAF | 16383.25 | 0.02330 | | ANAPC4 | 16996.25 | 0.02512 |
| GSTM1 | 16389 | 0.02332 | | CTDSPL | 17011.5 | 0.02516 |
| ENDOD1 | 16411.25 | 0.02339 | | TNFSF12 | 17054.25 | 0.02528 |
| hsa-mir-125a | 16411.5 | 0.02339 | | NFIC | 17123.75 | 0.02552 |
| MTF1 | 16411.5 | 0.02339 | | OR6K2 | 17138.25 | 0.02556 |
| SIAH2 | 16489.5 | 0.02362 | | GP6 | 17169.5 | 0.02565 |
| MEST | 16584 | 0.02392 | | FAM9A | 17236.75 | 0.02583 |
| KIAA0319L | 16588 | 0.02394 | | hsa-mir-330 | 17253.25 | 0.02586 |

| | | | | | | |
|---|---|---|---|---|---|---|
| TSPAN5 | 16612.5 | 0.02401 | | UBE2D2 | 17286 | 0.02591 |
| LGALS8 | 16630.75 | 0.02408 | | TMEM74 | 17297 | 0.02595 |
| OR6C3 | 16657.25 | 0.02416 | | BTF3 | 17416.75 | 0.02635 |
| hsa-mir-2467 | 16672.5 | 0.02421 | | TUBA1B | 17455.25 | 0.02647 |
| MTSS1L | 16705.5 | 0.02432 | | TYK2 | 17483 | 0.02656 |
| LRRC59 | 16755.5 | 0.02446 | | CLEC1A | 17505.25 | 0.02662 |
| FGF22 | 16907.5 | 0.02492 | | ZNF730 | 17539.75 | 0.02674 |
| APOBEC3B | 16950.25 | 0.02503 | | SLC5A2 | 17554 | 0.02677 |
| EMR1 | 16951.25 | 0.02504 | | ENHO | 17563.25 | 0.02680 |
| ZSCAN25 | 16972.25 | 0.02509 | | HIST1H2BH | 17598.25 | 0.02690 |
| ZNF514 | 17025 | 0.02524 | | SCEL | 17622.5 | 0.02696 |
| MUCL1 | 17105 | 0.02549 | | HOXB8 | 17684.75 | 0.02717 |
| RIMKLB | 17145.25 | 0.02560 | | FGD3 | 17706.25 | 0.02723 |
| FAM229B | 17199.75 | 0.02576 | | WDR45 | 17709.25 | 0.02724 |
| OR10H5 | 17238 | 0.02587 | | PSD3 | 17777.75 | 0.02745 |
| OR4C45 | 17264.75 | 0.02597 | | BRD4 | 17785.5 | 0.02748 |
| AK7 | 17264.75 | 0.02597 | | POU5F2 | 17859.5 | 0.02770 |
| PLEKHG7 | 17268.75 | 0.02598 | | KRTAP4-6 | 17865.5 | 0.02771 |
| DHCR7 | 17332 | 0.02617 | | SERPINB2 | 17961.75 | 0.02801 |
| GNB2L1 | 17379 | 0.02631 | | KCTD11 | 18012.25 | 0.02812 |
| ENO4 | 17440.5 | 0.02652 | | SGCE | 18043.75 | 0.02823 |
| hsa-mir-125b-1 | 17483.25 | 0.02665 | | ERH | 18045.25 | 0.02823 |
| IMPDH1 | 17502.75 | 0.02673 | | CAPN13 | 18051.25 | 0.02825 |
| LIG4 | 17640.5 | 0.02715 | | KIF9 | 18121 | 0.02849 |
| LIN54 | 17658.5 | 0.02721 | | PRPSAP1 | 18150.5 | 0.02858 |
| hsa-mir-4512 | 17665.75 | 0.02724 | | POLG2 | 18196 | 0.02872 |
| hsa-mir-1302-7 | 17676.25 | 0.02727 | | APCDD1L | 18244.25 | 0.02886 |
| hsa-mir-3196 | 17770.5 | 0.02758 | | HLA-DQA1 | 18259.5 | 0.02890 |
| ASB4 | 17774.75 | 0.02760 | | NUB1 | 18331 | 0.02912 |
| GINS2 | 17783 | 0.02763 | | CCDC80 | 18355.75 | 0.02921 |
| GNAQ | 17800.5 | 0.02768 | | NEK10 | 18376.25 | 0.02926 |
| DIO3 | 17817 | 0.02774 | | TRA2B | 18379 | 0.02927 |
| SPATA7 | 17823.25 | 0.02777 | | PHC1 | 18394.25 | 0.02932 |
| COX20 | 17886.25 | 0.02797 | | DOCK7 | 18405 | 0.02936 |
| SOX5 | 17887 | 0.02797 | | RBBP8 | 18432.5 | 0.02945 |
| SIGLEC7 | 17916.5 | 0.02804 | | LRSAM1 | 18492.25 | 0.02965 |
| C5orf52 | 17928.75 | 0.02807 | | GRINA | 18542.25 | 0.02983 |

| | | | | | | |
|---|---|---|---|---|---|---|
| ZCCHC14 | 17993.25 | 0.02831 | | hsa-mir-3193 | 18569.25 | 0.02994 |
| KIAA0922 | 18008 | 0.02836 | | PACS1 | 18585.5 | 0.02999 |
| FOXF2 | 18030.75 | 0.02845 | | SMC3 | 18675.75 | 0.03028 |
| hsa-mir-6885 | 18124.75 | 0.02870 | | HIST1H2AE | 18720.5 | 0.03040 |
| FAM110A | 18172.25 | 0.02887 | | WDR62 | 18791.25 | 0.03062 |
| SF3A2 | 18176 | 0.02888 | | PLEKHM2 | 18857.5 | 0.03083 |
| HMBOX1 | 18239.25 | 0.02906 | | TMBIM1 | 18952.25 | 0.03114 |
| SIT1 | 18252.5 | 0.02908 | | PTBP2 | 19014.25 | 0.03134 |
| OR2T8 | 18254.5 | 0.02909 | | GRSF1 | 19014.5 | 0.03134 |
| CDH1 | 18260.5 | 0.02911 | | HDGFRP2 | 19122.75 | 0.03166 |
| NonTargetingControlGuideForHuman_0820 | 18324.75 | 0.02934 | | CEP135 | 19151.5 | 0.03172 |
| HERC1 | 18343 | 0.02940 | | ACVR2B | 19232.25 | 0.03196 |
| NEK8 | 18368.75 | 0.02951 | | MSANTD4 | 19251.25 | 0.03200 |
| BIK | 18453.75 | 0.02979 | | C9orf89 | 19442 | 0.03267 |
| MIB2 | 18473 | 0.02984 | | HAUS7 | 19495 | 0.03282 |
| C16orf55 | 18492 | 0.02989 | | SUPT20HL1 | 19504.75 | 0.03285 |
| LTBP2 | 18513.25 | 0.02997 | | PDLIM1 | 19514.75 | 0.03289 |
| GOLGA7 | 18533 | 0.03001 | | GCM1 | 19565 | 0.03306 |
| MRPS18B | 18550.5 | 0.03006 | | hsa-mir-3124 | 19678 | 0.03346 |
| KDM5C | 18559.25 | 0.03008 | | P2RX3 | 19680.75 | 0.03347 |
| OTX1 | 18579 | 0.03015 | | hsa-mir-4682 | 19721 | 0.03361 |
| SLX4 | 18596 | 0.03020 | | NARS2 | 19723.75 | 0.03362 |
| GBP5 | 18634.5 | 0.03033 | | hsa-mir-4475 | 19727 | 0.03363 |
| TM4SF1 | 18657.5 | 0.03041 | | OR2AE1 | 19768.5 | 0.03377 |
| EIF4E1B | 18690 | 0.03052 | | KRTAP12-3 | 19770.25 | 0.03377 |
| C1orf173 | 18761 | 0.03072 | | PPIF | 19796.5 | 0.03387 |
| DYNC1LI2 | 18814.5 | 0.03088 | | POLR2J | 19894.25 | 0.03417 |
| CCDC61 | 18874.25 | 0.03109 | | NDUFA4 | 19926.75 | 0.03430 |
| REPS1 | 18925.5 | 0.03126 | | HOXD11 | 19977.75 | 0.03449 |
| KCTD12 | 18979.5 | 0.03142 | | SLC4A7 | 20002.75 | 0.03459 |
| PRR25 | 18989.75 | 0.03144 | | CCL25 | 20147.25 | 0.03509 |
| COX17 | 19076 | 0.03169 | | RNF182 | 20202.5 | 0.03525 |
| PUSL1 | 19102 | 0.03178 | | SRA1 | 20222.75 | 0.03533 |
| GDPD4 | 19158 | 0.03196 | | HDAC9 | 20236.25 | 0.03539 |
| ZNF28 | 19232.5 | 0.03220 | | RECK | 20267.75 | 0.03551 |
| METRN | 19246.25 | 0.03224 | | FAM177B | 20328.25 | 0.03573 |
| EIF3J | 19249.5 | 0.03225 | | GNAZ | 20342.75 | 0.03576 |

| | | | | | | |
|---|---|---|---|---|---|---|
| SERTAD4 | 19254.25 | 0.03227 | | TWF1 | 20350.75 | 0.03579 |
| HECTD1 | 19263.25 | 0.03230 | | FGGY | 20362.5 | 0.03584 |
| NAIF1 | 19311.75 | 0.03244 | | SPRED3 | 20380.5 | 0.03591 |
| HIF1A | 19339 | 0.03253 | | CCL8 | 20406.75 | 0.03602 |
| TMEM209 | 19381.75 | 0.03270 | | FUT8 | 20443.75 | 0.03615 |
| hsa-mir-346 | 19488.5 | 0.03305 | | KCNIP4 | 20541.75 | 0.03648 |
| FZD5 | 19557 | 0.03327 | | NonTargetingControlGuideForHuman_0418 | 20587.5 | 0.03662 |
| ST7L | 19604.25 | 0.03344 | | DSG4 | 20725.75 | 0.03711 |
| XAB2 | 19630 | 0.03352 | | CDKN2C | 20728.25 | 0.03712 |
| PER2 | 19666 | 0.03367 | | hsa-mir-4660 | 20798 | 0.03734 |
| PCDHGB4 | 19702.5 | 0.03378 | | FAM161B | 20806.75 | 0.03739 |
| RORB | 19748 | 0.03395 | | hsa-mir-125b-1 | 20810.5 | 0.03739 |
| ATRNL1 | 19786 | 0.03404 | | BET1 | 20890.5 | 0.03772 |
| SLBP | 19829.5 | 0.03420 | | MTRNR2L7 | 20933 | 0.03789 |
| SATL1 | 19861.75 | 0.03431 | | C14orf28 | 20986.5 | 0.03809 |
| MEF2D | 19875 | 0.03434 | | BEND4 | 21000.5 | 0.03814 |
| TOR1AIP1 | 19881.5 | 0.03437 | | UVRAG | 21002 | 0.03815 |
| CYB561A3 | 19982 | 0.03468 | | DDX4 | 21021 | 0.03823 |
| hsa-mir-4283-2 | 19988.25 | 0.03470 | | KRT77 | 21208.75 | 0.03891 |
| CCDC91 | 20079.25 | 0.03504 | | P2RY10 | 21225.25 | 0.03897 |
| PKN2 | 20123.25 | 0.03520 | | MYRIP | 21240.25 | 0.03905 |
| hsa-mir-5688 | 20253.25 | 0.03571 | | MYH1 | 21271.5 | 0.03915 |
| NOL7 | 20271.75 | 0.03578 | | C2CD4B | 21283.75 | 0.03919 |
| ZKSCAN5 | 20279.5 | 0.03582 | | TIMM10B | 21328.5 | 0.03938 |
| CCDC28A | 20341 | 0.03601 | | TSPAN2 | 21371 | 0.03951 |
| TNRC18 | 20415 | 0.03625 | | SMC6 | 21491.5 | 0.03999 |
| AMOTL2 | 20457 | 0.03636 | | FAM129B | 21511.5 | 0.04006 |
| SNRNP35 | 20503 | 0.03652 | | SCUBE1 | 21604.25 | 0.04044 |
| RAP1GDS1 | 20511.25 | 0.03656 | | RABEP1 | 21611.25 | 0.04046 |
| hsa-mir-939 | 20548.25 | 0.03668 | | hsa-mir-3195 | 21620.75 | 0.04048 |
| CYTH1 | 20569.75 | 0.03680 | | hsa-mir-606 | 21747.75 | 0.04099 |
| AHSG | 20594 | 0.03688 | | TAB3 | 21780.25 | 0.04113 |
| CHCHD10 | 20673.5 | 0.03713 | | MATN3 | 21805.25 | 0.04119 |
| CBLL1 | 20749.25 | 0.03738 | | C19orf59 | 21854.5 | 0.04137 |
| KRTAP19-7 | 20780 | 0.03748 | | CAPN3 | 21887.5 | 0.04150 |
| PRSS16 | 20823.25 | 0.03765 | | SEC61A1 | 21890 | 0.04152 |
| GOLGA7B | 20832.25 | 0.03768 | | MGST3 | 21900 | 0.04155 |

| | | | | | | |
|---|---|---|---|---|---|---|
| FAM32A | 20853.25 | 0.03777 | | GCNT2 | 21951 | 0.04172 |
| ARSD | 20950.75 | 0.03813 | | STXBP1 | 22009.5 | 0.04193 |
| hsa-mir-1299 | 20984.25 | 0.03827 | | CRIM1 | 22037 | 0.04204 |
| PGM3 | 21001.25 | 0.03835 | | NonTargetingControlGuideForHuman_0820 | 22058.75 | 0.04213 |
| APOBEC1 | 21027.75 | 0.03844 | | FYB | 22138.25 | 0.04243 |
| SFR1 | 21044.75 | 0.03853 | | TMEM130 | 22140 | 0.04244 |
| ENO4 | 21216.25 | 0.03922 | | VPS28 | 22170.25 | 0.04258 |
| SLC6A18 | 21248.25 | 0.03934 | | ANKFY1 | 22229.5 | 0.04281 |
| CXADR | 21280.5 | 0.03945 | | LARGE | 22260.5 | 0.04294 |
| ZFYVE28 | 21305 | 0.03953 | | EPB41L1 | 22260.75 | 0.04294 |
| DNAJC16 | 21322.5 | 0.03959 | | UBP1 | 22327 | 0.04320 |
| SPDYA | 21374.25 | 0.03981 | | ZCCHC2 | 22348.25 | 0.04327 |
| hsa-mir-4435-1 | 21389 | 0.03986 | | IER2 | 22350.5 | 0.04328 |
| ARL2BP | 21435.75 | 0.04007 | | SRPK2 | 22466 | 0.04369 |
| HLCS | 21550.5 | 0.04049 | | OR11A1 | 22511.5 | 0.04389 |
| CCDC62 | 21574.5 | 0.04058 | | SLC39A7 | 22531 | 0.04398 |
| RAD21 | 21602 | 0.04069 | | MORF4L2 | 22544 | 0.04402 |
| hsa-mir-30d | 21621.5 | 0.04077 | | FBLN5 | 22597.75 | 0.04425 |
| CASKIN2 | 21659 | 0.04089 | | ZDHHC2 | 22612.75 | 0.04430 |
| EMID1 | 21703.75 | 0.04106 | | BTF3L4 | 22643.25 | 0.04443 |
| TWIST2 | 21747 | 0.04123 | | CSNK1A1L | 22782.5 | 0.04491 |
| SMC5 | 21985.5 | 0.04216 | | POLE2 | 22843.5 | 0.04517 |
| ECM1 | 21987.75 | 0.04217 | | IMPACT | 22942.5 | 0.04553 |
| APPL1 | 21996.25 | 0.04220 | | CXorf61 | 23000.25 | 0.04577 |
| COL11A1 | 22032.75 | 0.04234 | | LRRFIP1 | 23009.25 | 0.04581 |
| MRPS24 | 22057.75 | 0.04244 | | AAED1 | 23017.25 | 0.04584 |
| THEMIS2 | 22069.25 | 0.04248 | | XKRX | 23111.25 | 0.04621 |
| FOSL2 | 22149.5 | 0.04278 | | ASB16 | 23134.75 | 0.04630 |
| PUS10 | 22188 | 0.04294 | | TMEM51 | 23146 | 0.04634 |
| GPR12 | 22200 | 0.04298 | | KCNT1 | 23225.5 | 0.04668 |
| SCAF11 | 22342 | 0.04349 | | INSL4 | 23258.5 | 0.04682 |
| ISCU | 22406 | 0.04376 | | ANGPTL3 | 23277.25 | 0.04686 |
| EIF3I | 22408 | 0.04377 | | RNF133 | 23305 | 0.04698 |
| POLR2B | 22472.75 | 0.04404 | | RAI2 | 23426.5 | 0.04752 |
| ACSL1 | 22531.25 | 0.04429 | | PEX11A | 23452.5 | 0.04760 |
| ZNF232 | 22575 | 0.04448 | | RBM15 | 23495.5 | 0.04781 |
| KAT5 | 22613.75 | 0.04463 | | FAM150B | 23531.25 | 0.04796 |

| | | | | | | |
|---|---|---|---|---|---|---|
| ZGPAT | 22639.75 | 0.04472 | | FTH1 | 23536.75 | 0.04799 |
| ZBBX | 22667 | 0.04484 | | FGB | 23657 | 0.04853 |
| RLN3 | 22675 | 0.04486 | | NCAPG2 | 23726 | 0.04880 |
| PPCS | 22745.75 | 0.04516 | | MAML2 | 23741.5 | 0.04886 |
| TMIGD1 | 22894.5 | 0.04571 | | CCDC151 | 23765.5 | 0.04895 |
| RPL6 | 22926.25 | 0.04584 | | ZNF592 | 23786.5 | 0.04904 |
| ST8SIA6 | 22995.25 | 0.04613 | | ZNF804A | 23851.5 | 0.04932 |
| MRFAP1 | 22998.5 | 0.04615 | | GPR87 | 23872.75 | 0.04943 |
| POLR3G | 23053.5 | 0.04640 | | FAM117B | 23911 | 0.04957 |
| PRKCI | 23093.5 | 0.04657 | | CARHSP1 | 23931.5 | 0.04966 |
| BTBD2 | 23193.75 | 0.04696 | | GDNF | 23938.25 | 0.04969 |
| UCP3 | 23207.5 | 0.04702 | | | | |
| NPTX1 | 23211.5 | 0.04704 | | | | |
| LRSAM1 | 23371.75 | 0.04772 | | | | |
| hsa-mir-3684 | 23532.75 | 0.04836 | | | | |
| RHOXF2B | 23593.75 | 0.04859 | | | | |
| E2F6 | 23621.5 | 0.04869 | | | | |
| TOR3A | 23716.5 | 0.04906 | | | | |
| RNF113B | 23819.75 | 0.04951 | | | | |
| CNTNAP2 | 23868.5 | 0.04971 | | | | |
| LHX1 | 23919.25 | 0.04995 | | | | |

#### Table 2: Transcriptional dynamics of genes depleted in replicate 2CT genome-scale CRISPR screens

| Gene name | Fold change mRNA 6hr coculture/0hr | pvalue (-log10) |
|---|---|---|
| *RPAP2* | 3.83 | 1.386153 |
| *BIRC2* | 3.49 | 2.095161 |
| *ESD* | 2.78 | 1.655263 |
| *TWF1* | 2.15 | 1.127761 |
| *EGR2* | 2.13 | 1.422591 |
| *SCAF11* | 1.92 | 2.493371 |
| *TOR1AIP1* | 1.73 | 1.654647 |
| *LPGAT1* | 1.67 | 1.594682 |
| *TXLNB* | 1.65 | 0.736541 |
| *SERTAD2* | 1.62 | 1.964377 |
| *COG3* | 1.55 | 0.951055 |
| *ALG11* | 1.53 | 0.630745 |
| *RBL2* | 1.42 | 1.730386 |
| *BCL2* | 1.39 | 1.333402 |
| *ENDOD1* | 1.35 | 0.880535 |
| *AKIRIN1* | 1.31 | 0.846618 |
| *EAF2* | 1.3 | 0.313509 |
| *C14orf119* | 1.27 | 0.752152 |
| *PRPF3* | 1.22 | 0.822365 |
| *ENY2* | 1.21 | 0.719565 |
| *TRA2B* | 1.18 | 0.809857 |
| *SNX1* | 1.17 | 0.933405 |
| *THRAP3* | 1.17 | 1.068386 |
| *ISCU* | 1.09 | 0.277723 |
| *WDR45B* | 1.03 | 0.136798 |
| *EIF3I* | 1.02 | 0.092091 |
| *TMX4* | -1.03 | 0.052917 |
| *ARSD* | -1.05 | 0.069726 |
| *HHLA3* | -1.05 | 0.042312 |
| *PPP2R5D* | -1.06 | 0.239111 |
| *SART3* | -1.06 | 0.201343 |
| *CINP* | -1.13 | 0.501461 |
| *MANSC1* | -1.13 | 0.079621 |
| *NUDC* | -1.29 | 0.689154 |
| *SLC25A25* | -1.31 | 1.128394 |
| *RPS16* | -1.36 | 1.127576 |

| | | |
|---|---|---|
| *FAM32A* | -1.4 | 1.213536 |
| *DPM2* | -1.62 | 2.076065 |
| *C15orf61* | -1.72 | 1.713913 |
| *CYB5R3* | -1.76 | 2.226299 |
| *NEK8* | -1.81 | 2.116487 |
| *FBXL15* | -2.15 | 2.270763 |

#### Table 3: Mapping top 2CT CRISPR depleted genes to protein-coding genes

| Gene ID | Mapped ID | Gene name/Gene Symbol |
|---|---|---|
| HUMAN\|HGNC=13921\|UniProtKB=O95870 | ABHD16A | Protein ABHD16A;ABHD16A;ortholog |
| HUMAN\|HGNC=19195\|UniProtKB=Q6H8Q1 | ABLIM2 | Actin-binding LIM protein 2;ABLIM2;ortholog |
| HUMAN\|HGNC=18075\|UniProtKB=Q9NZD4 | AHSP | Alpha-hemoglobin-stabilizing protein;AHSP;ortholog |
| HUMAN\|HGNC=25744\|UniProtKB=Q9H9L7 | AKIRIN1 | Akirin-1;AKIRIN1;ortholog |
| HUMAN\|HGNC=32456\|UniProtKB=Q2TAA5 | ALG11 | GDP-Man:Man(3)GlcNAc(2)-PP-Dol alpha-1,2-mannosyltransferase;ALG11;ortholog |
| HUMAN\|HGNC=28658\|UniProtKB=Q6DCA0 | AMMECR1L | AMMECR1-like protein;AMMECR1L;ortholog |
| HUMAN\|HGNC=29135\|UniProtKB=Q6UB98 | ANKRD12 | Ankyrin repeat domain-containing protein 12;ANKRD12;ortholog |
| HUMAN\|HGNC=23725\|UniProtKB=Q5T5U3 | ARHGAP21 | Rho GTPase-activating protein 21;ARHGAP21;ortholog |
| HUMAN\|HGNC=25361\|UniProtKB=Q8N264 | ARHGAP24 | Rho GTPase-activating protein 24;ARHGAP24;ortholog |
| HUMAN\|HGNC=721\|UniProtKB=P54793 | ARSF | Arylsulfatase F;ARSF;ortholog |
| HUMAN\|HGNC=588\|UniProtKB=O94817 | ATG12 | Ubiquitin-like protein ATG12;ATG12;ortholog |
| HUMAN\|HGNC=855\|UniProtKB=P27449 | ATP6V0C | V-type proton ATPase 16 kDa proteolipid subunit;ATP6V0C;ortholog |
| HUMAN\|HGNC=24137\|UniProtKB=Q6L9W6 | B4GALNT3 | Beta-1,4-N-acetylgalactosaminyltransferase 3;B4GALNT3;ortholog |
| HUMAN\|HGNC=950\|UniProtKB=Q92560 | BAP1 | Ubiquitin carboxyl-terminal hydrolase BAP1;BAP1;ortholog |
| HUMAN\|HGNC=990\|UniProtKB=P10415 | BCL2 | Apoptosis regulator Bcl-2;BCL2;ortholog |
| HUMAN\|HGNC=590\|UniProtKB=Q13490 | BIRC2 | Baculoviral IAP repeat-containing protein 2;BIRC2;ortholog |
| HUMAN\|HGNC=15519\|UniProtKB=Q14137 | BOP1 | Ribosome biogenesis protein BOP1;BOP1;ortholog |
| HUMAN\|HGNC=27441\|UniProtKB=Q3KP22 | C11orf85 | Membrane-anchored junction protein;C11orf85;ortholog |
| HUMAN\|HGNC=32331\|UniProtKB=Q5T5A4 | C1orf194 | Uncharacterized protein C1orf194;C1orf194;ortholog |
| HUMAN\|HGNC=28250\|UniProtKB=Q6P1W5 | C1orf94 | Uncharacterized protein C1orf94;C1orf94;ortholog |
| HUMAN\|HGNC=28772\|UniProtKB=Q9BVC5 | C2orf49 | Ashwin;C2orf49;ortholog |
| HUMAN\|HGNC=26951\|UniProtKB=Q86YL5 | C8orf42 | Testis development-related protein;TDRP;ortholog |
| HUMAN\|HGNC=18200\|UniProtKB=Q8WXE0 | CASKIN2 | Caskin-2;CASKIN2;ortholog |
| HUMAN\|HGNC=1516\|UniProtKB=P04040 | CAT | Catalase;CAT;ortholog |
| HUMAN\|HGNC=29588\|UniProtKB=Q5BJE1 | CCDC178 | Coiled-coil domain-containing protein 178;CCDC178;ortholog |
| HUMAN\|HGNC=30723\|UniProtKB=Q6P9F0 | CCDC62 | Coiled-coil domain-containing protein 62;CCDC62;ortholog |

| | | |
|---|---|---|
| HUMAN\|HGNC=1578\|UniProtKB=P20248 | CCNA2 | Cyclin-A2;CCNA2;ortholog |
| HUMAN\|HGNC=1766\|UniProtKB=Q9ULB5 | CDH7 | Cadherin-7;CDH7;ortholog |
| HUMAN\|HGNC=1837\|UniProtKB=P53567 | CEBPG | CCAAT/enhancer-binding protein gamma;CEBPG;ortholog |
| HUMAN\|HGNC=1876\|UniProtKB=O15519 | CFLAR | CASP8 and FADD-like apoptosis regulator;CFLAR;ortholog |
| HUMAN\|HGNC=23789\|UniProtKB=Q9BW66 | CINP | Cyclin-dependent kinase 2-interacting protein;CINP;ortholog |
| HUMAN\|HGNC=2080\|UniProtKB=P54105 | CLNS1A | Methylosome subunit pICln;CLNS1A;ortholog |
| HUMAN\|HGNC=16999\|UniProtKB=Q92989 | CLP1 | Polyribonucleotide 5'-hydroxyl-kinase Clp1;CLP1;ortholog |
| HUMAN\|HGNC=18623\|UniProtKB=Q96MW5 | COG8 | Conserved oligomeric Golgi complex subunit 8;COG8;ortholog |
| HUMAN\|HGNC=17888\|UniProtKB=O75575 | CRCP | DNA-directed RNA polymerase III subunit RPC9;CRCP;ortholog |
| HUMAN\|HGNC=2396\|UniProtKB=P53673 | CRYBA4 | Beta-crystallin A4;CRYBA4;ortholog |
| HUMAN\|HGNC=15982\|UniProtKB=Q8IWT3 | CUL9 | Cullin-9;CUL9;ortholog |
| HUMAN\|HGNC=10664\|UniProtKB=Q6UX04 | CWC27 | Peptidyl-prolyl cis-trans isomerase CWC27 homolog;CWC27;ortholog |
| HUMAN\|HGNC=2873\|UniProtKB=P00387 | CYB5R3 | NADH-cytochrome b5 reductase 3;CYB5R3;ortholog |
| HUMAN\|HGNC=21191\|UniProtKB=P59103 | DAOA | D-amino acid oxidase activator;DAOA;ortholog |
| HUMAN\|HGNC=17347\|UniProtKB=Q9BUQ8 | DDX23 | Probable ATP-dependent RNA helicase DDX23;DDX23;ortholog |
| HUMAN\|HGNC=25360\|UniProtKB=Q5T1V6 | DDX59 | Probable ATP-dependent RNA helicase DDX59;DDX59;ortholog |
| HUMAN\|HGNC=15861\|UniProtKB=Q9H5Z1 | DHX35 | Probable ATP-dependent RNA helicase DHX35;DHX35;ortholog |
| HUMAN\|HGNC=2680\|UniProtKB=Q9BTC0 | DIDO1 | Death-inducer obliterator 1;DIDO1;ortholog |
| HUMAN\|HGNC=2902\|UniProtKB=Q92796 | DLG3 | Disks large homolog 3;DLG3;ortholog |
| HUMAN\|HGNC=29157\|UniProtKB=Q9Y2G8 | DNAJC16 | DnaJ homolog subfamily C member 16;DNAJC16;ortholog |
| HUMAN\|HGNC=2981\|UniProtKB=Q9ULA0 | DNPEP | Aspartyl aminopeptidase;DNPEP;ortholog |
| HUMAN\|HGNC=23028\|UniProtKB=Q9H4A9 | DPEP2 | Dipeptidase 2;DPEP2;ortholog |
| HUMAN\|HGNC=3011\|UniProtKB=Q07507 | DPT | Dermatopontin;DPT;ortholog |
| HUMAN\|HGNC=3208\|UniProtKB=P24534 | EEF1B2 | Elongation factor 1-beta;EEF1B2;ortholog |
| HUMAN\|HGNC=3226\|UniProtKB=P98172 | EFNB1 | Ephrin-B1;EFNB1;ortholog |
| HUMAN\|HGNC=3239\|UniProtKB=P11161 | EGR2 | E3 SUMO-protein ligase EGR2;EGR2;ortholog |
| HUMAN\|HGNC=3272\|UniProtKB=Q13347 | EIF3I | Eukaryotic translation initiation factor 3 subunit I;EIF3I;ortholog |
| HUMAN\|HGNC=3296\|UniProtKB=Q04637 | EIF4G1 | Eukaryotic translation initiation factor 4 gamma 1;EIF4G1;ortholog |
| HUMAN\|HGNC=3297\|UniProtKB=P78344 | EIF4G2 | Eukaryotic translation initiation factor 4 gamma 2;EIF4G2;ortholog |
| HUMAN\|HGNC=28430\|UniProtKB=Q9BV81 | EMC6 | ER membrane protein complex subunit 6;EMC6;ortholog |
| HUMAN\|HGNC=1316\|UniProtKB=Q9HC35 | EML4 | Echinoderm microtubule-associated protein-like 4;EML4;ortholog |
| HUMAN\|HGNC=24449\|UniProtKB=Q9NPA8 | ENY2 | Transcription and mRNA export factor ENY2;ENY2;ortholog |
| HUMAN\|HGNC=3415\|UniProtKB=P01588 | EPO | Erythropoietin;EPO;ortholog |
| HUMAN\|HGNC=3471\|UniProtKB=P11474 | ESRRA | Steroid hormone receptor ERR1;ESRRA;ortholog |

| | | |
|---|---|---|
| HUMAN\|HGNC=3522\|UniProtKB=O9567 7 | EYA4 | Eyes absent homolog 4;EYA4;ortholog |
| HUMAN\|HGNC=24563\|UniProtKB=Q9Y4 21 | FAM32A | Protein FAM32A;FAM32A;ortholog |
| HUMAN\|HGNC=1253\|UniProtKB=P5849 9 | FAM3B | Protein FAM3B;FAM3B;ortholog |
| HUMAN\|HGNC=18786\|UniProtKB=Q143 20 | FAM50A | Protein FAM50A;FAM50A;ortholog |
| HUMAN\|HGNC=17800\|UniProtKB=Q9NS D9 | FARSB | Phenylalanine--tRNA ligase beta subunit;FARSB;ortholog |
| HUMAN\|HGNC=3595\|UniProtKB=Q1451 7 | FAT1 | Protocadherin Fat 1;FAT1;ortholog |
| HUMAN\|HGNC=3600\|UniProtKB=P2314 2 | FBLN1 | Fibulin-1;FBLN1;ortholog |
| HUMAN\|HGNC=16712\|UniProtKB=Q969 H0 | FBXW7 | F-box/WD repeat-containing protein 7;FBXW7;ortholog |
| HUMAN\|HGNC=18522\|UniProtKB=Q5VV 16 | FOXD4L5 | Forkhead box protein D4-like 5;FOXD4L5;ortholog |
| HUMAN\|HGNC=30968\|UniProtKB=A6NG K3 | GAGE10 | G antigen 10;GAGE10;ortholog |
| HUMAN\|HGNC=14264\|UniProtKB=Q9NY 12 | GAR1 | H/ACA ribonucleoprotein complex subunit 1;GAR1;ortholog |
| HUMAN\|HGNC=11960\|UniProtKB=Q6Y7 W6 | GIGYF2 | PERQ amino acid-rich with GYF domain-containing protein 2;GIGYF2;ortholog |
| HUMAN\|HGNC=28980\|UniProtKB=Q146 91 | GINS1 | DNA replication complex GINS protein PSF1;GINS1;ortholog |
| HUMAN\|HGNC=4374\|UniProtKB=O6023 4 | GMFG | Glia maturation factor gamma;GMFG;ortholog |
| HUMAN\|HGNC=18707\|UniProtKB=Q7Z6 J2 | GRASP | General receptor for phosphoinositides 1-associated scaffold protein;GRASP;ortholog |
| HUMAN\|HGNC=4579\|UniProtKB=P3908 6 | GRIK1 | Glutamate receptor ionotropic, kainate 1;GRIK1;ortholog |
| HUMAN\|HGNC=33383\|UniProtKB=A0PJZ 3 | GXYLT2 | Glucoside xylosyltransferase 2;GXYLT2;ortholog |
| HUMAN\|HGNC=4801\|UniProtKB=P4093 9 | HADH | Trifunctional enzyme subunit alpha, mitochondrial;HADHA;ortholog |
| HUMAN\|HGNC=4803\|UniProtKB=P5508 4 | HADHB | Trifunctional enzyme subunit beta, mitochondrial;HADHB;ortholog |
| HUMAN\|HGNC=4848\|UniProtKB=O4361 3 | HCRTR1 | Orexin receptor type 1;HCRTR1;ortholog |
| HUMAN\|HGNC=28982\|UniProtKB=Q127 66 | HMGXB3 | HMG domain-containing protein 3;HMGXB3;ortholog |
| HUMAN\|HGNC=25451\|UniProtKB=Q1K MD3 | HNRNPUL 2 | Heterogeneous nuclear ribonucleoprotein U-like protein 2;HNRNPUL2;ortholog |
| HUMAN\|HGNC=25155\|UniProtKB=Q86X E5 | HOGA1 | 4-hydroxy-2-oxoglutarate aldolase, mitochondrial;HOGA1;ortholog |
| HUMAN\|HGNC=5129\|UniProtKB=P3127 3 | HOXC8 | Homeobox protein Hox-C8;HOXC8;ortholog |
| HUMAN\|HGNC=5961\|UniProtKB=Q9Y6K 9 | IKBKG | NF-kappa-B essential modulator;IKBKG;ortholog |
| HUMAN\|HGNC=5993\|UniProtKB=P1477 8 | IL1R1 | Interleukin-1 receptor type 1;IL1R1;ortholog |
| HUMAN\|HGNC=15561\|UniProtKB=Q9UB H0 | IL36RN | Interleukin-36 receptor antagonist protein;IL36RN;ortholog |
| HUMAN\|HGNC=6029\|UniProtKB=P1524 8 | IL9 | Interleukin-9;IL9;ortholog |
| HUMAN\|HGNC=29882\|UniProtKB=Q9H1 K1 | ISCU | Iron-sulfur cluster assembly enzyme ISCU, mitochondrial;ISCU;ortholog |
| HUMAN\|HGNC=28977\|UniProtKB=P539 90 | IST1 | IST1 homolog;IST1;ortholog |
| HUMAN\|HGNC=6150\|UniProtKB=P0675 6 | ITGAV | Integrin alpha-V;ITGAV;ortholog |
| HUMAN\|HGNC=6173\|UniProtKB=O4373 6 | ITM2A | Integral membrane protein 2A;ITM2A;ortholog |
| HUMAN\|HGNC=21071\|UniProtKB=Q6PH W0 | IYD | Iodotyrosine deiodinase 1;IYD;ortholog |
| HUMAN\|HGNC=6211\|UniProtKB=P2335 2 | KAL1 | Anosmin-1;ANOS1;ortholog |

| | | |
|---|---|---|
| HUMAN\|HGNC=6217\|UniProtKB=Q9BVA0 | KATNB1 | Katanin p80 WD40 repeat-containing subunit B1;KATNB1;ortholog |
| HUMAN\|HGNC=6246\|UniProtKB=Q9H3M0 | KCNF1 | Potassium voltage-gated channel subfamily F member 1;KCNF1;ortholog |
| HUMAN\|HGNC=6249\|UniProtKB=Q9UJ96 | KCNG2 | Potassium voltage-gated channel subfamily G member 2;KCNG2;ortholog |
| HUMAN\|HGNC=6304\|UniProtKB=P24390 | KDELR1 | ER lumen protein-retaining receptor 1;KDELR1;ortholog |
| HUMAN\|HGNC=29266\|UniProtKB=Q9P2G3 | KLHL14 | Kelch-like protein 14;KLHL14;ortholog |
| HUMAN\|HGNC=16905\|UniProtKB=O60662 | KLHL41 | Kelch-like protein 41;KLHL41;ortholog |
| HUMAN\|HGNC=6356\|UniProtKB=Q96PQ7 | KLHL5 | Kelch-like protein 5;KLHL5;ortholog |
| HUMAN\|HGNC=6359\|UniProtKB=Q9UBX7 | KLK11 | Kallikrein-11;KLK11;ortholog |
| HUMAN\|HGNC=20523\|UniProtKB=P60371 | KRTAP10-6 | Keratin-associated protein 10-6;KRTAP10-6;ortholog |
| HUMAN\|HGNC=23598\|UniProtKB=Q6L8H2 | KRTAP5-3 | Keratin-associated protein 5-3;KRTAP5-3;ortholog |
| HUMAN\|HGNC=17095\|UniProtKB=Q15031 | LARS2 | Probable leucine--tRNA ligase, mitochondrial;LARS2;ortholog |
| HUMAN\|HGNC=29460\|UniProtKB=Q5TA81 | LCE2C | Late cornified envelope protein 2C;LCE2C;ortholog |
| HUMAN\|HGNC=17787\|UniProtKB=O14910 | LIN7A | Protein lin-7 homolog A;LIN7A;ortholog |
| HUMAN\|Gene=LPPR5\|UniProtKB=Q32ZL2 | LPPR5 | Phospholipid phosphatase-related protein type 5;PLPPR5;ortholog |
| HUMAN\|HGNC=6690\|UniProtKB=Q12912 | LRMP | Lymphoid-restricted membrane protein;LRMP;ortholog |
| HUMAN\|HGNC=19742\|UniProtKB=Q96L50 | LRR1 | Leucine-rich repeat protein 1;LRR1;ortholog |
| HUMAN\|HGNC=19410\|UniProtKB=Q86VH5 | LRRTM3 | Leucine-rich repeat transmembrane neuronal protein 3;LRRTM3;ortholog |
| HUMAN\|HGNC=13940\|UniProtKB=Q9Y333 | LSM2 | U6 snRNA-associated Sm-like protein LSm2;LSM2;ortholog |
| HUMAN\|HGNC=1968\|UniProtKB=Q99698 | LYST | Lysosomal-trafficking regulator;LYST;ortholog |
| HUMAN\|HGNC=25183\|UniProtKB=Q8TC57 | M1AP | Meiosis 1 arrest protein;M1AP;ortholog |
| HUMAN\|HGNC=6780\|UniProtKB=Q9ULX9 | MAFF | Transcription factor MafF;MAFF;ortholog |
| HUMAN\|HGNC=6823\|UniProtKB=Q9UKM7 | MAN1B1 | Endoplasmic reticulum mannosyl-oligosaccharide 1,2-alpha-mannosidase;MAN1B1;ortholog |
| HUMAN\|HGNC=13961\|UniProtKB=P35410 | MAS1L | Mas-related G-protein coupled receptor MRG;MAS1L;ortholog |
| HUMAN\|HGNC=6904\|UniProtKB=P31153 | MAT2A | S-adenosylmethionine synthase isoform type-2;MAT2A;ortholog |
| HUMAN\|HGNC=19866\|UniProtKB=Q05BQ5 | MBTD1 | MBT domain-containing protein 1;MBTD1;ortholog |
| HUMAN\|HGNC=7095\|UniProtKB=O15344 | MID1 | E3 ubiquitin-protein ligase Midline-1;MID1;ortholog |
| HUMAN\|HGNC=21460\|UniProtKB=Q8TD10 | MIPOL1 | Mirror-image polydactyly gene 1 protein;MIPOL1;ortholog |
| HUMAN\|HGNC=29636\|UniProtKB=Q8NEH6 | MNS1 | Meiosis-specific nuclear structural protein 1;MNS1;ortholog |
| HUMAN\|HGNC=17617\|UniProtKB=Q96LA9 | MRGPRX4 | Mas-related G-protein coupled receptor member X4;MRGPRX4;ortholog |
| HUMAN\|HGNC=16635\|UniProtKB=P82673 | MRPS35 | 28S ribosomal protein S35, mitochondrial;MRPS35;ortholog |
| HUMAN\|HGNC=7394\|UniProtKB=P07438 | MT1B | Metallothionein-1B;MT1B;ortholog |
| HUMAN\|HGNC=7411\|UniProtKB=O94776 | MTA2 | Metastasis-associated protein MTA2;MTA2;ortholog |
| HUMAN\|HGNC=7748\|UniProtKB=Q6P3R8 | NEK5 | Serine/threonine-protein kinase Nek5;NEK5;ortholog |
| HUMAN\|HGNC=29933\|UniProtKB=Q86WI3 | NLRC5 | Protein NLRC5;NLRC5;ortholog |

| | | |
|---|---|---|
| HUMAN\|HGNC=17877\|UniProtKB=Q9HAN9 | NMNAT1 | Nicotinamide/nicotinic acid mononucleotide adenylyltransferase 1;NMNAT1;ortholog |
| HUMAN\|HGNC=32203\|UniProtKB=Q5H8A3 | NMS | Neuromedin-S;NMS;ortholog |
| HUMAN\|HGNC=16821\|UniProtKB=P78316 | NOP14 | Nucleolar protein 14;NOP14;ortholog |
| HUMAN\|HGNC=7895\|UniProtKB=Q99743 | NPAS2 | Neuronal PAS domain-containing protein 2;NPAS2;ortholog |
| HUMAN\|HGNC=14124\|UniProtKB=Q12980 | NPRL3 | Nitrogen permease regulator 3-like protein;NPRL3;ortholog |
| HUMAN\|HGNC=8266\|UniProtKB=Q8NH16 | OR2L2 | Olfactory receptor 2L2;OR2L2;ortholog |
| HUMAN\|HGNC=8268\|UniProtKB=Q96R28 | OR2M2 | Olfactory receptor 2M2;OR2M2;ortholog |
| HUMAN\|HGNC=8510\|UniProtKB=Q92882 | OSTF1 | Osteoclast-stimulating factor 1;OSTF1;ortholog |
| HUMAN\|HGNC=20882\|UniProtKB=Q9NWT1 | PAK1IP1 | p21-activated protein kinase-interacting protein 1;PAK1IP1;ortholog |
| HUMAN\|HGNC=8621\|UniProtKB=P23759 | PAX7 | Paired box protein Pax-7;PAX7;ortholog |
| HUMAN\|HGNC=9719\|UniProtKB=P50542 | PEX5 | Peroxisomal targeting signal 1 receptor;PEX5;ortholog |
| HUMAN\|HGNC=8940\|UniProtKB=O14832 | PHYH | Phytanoyl-CoA dioxygenase, peroxisomal;PHYH;ortholog |
| HUMAN\|HGNC=8971\|UniProtKB=O00443 | PIK3C2A | Phosphatidylinositol 4-phosphate 3-kinase C2 domain-containing subunit alpha;PIK3C2A;ortholog |
| HUMAN\|HGNC=20764\|UniProtKB=Q96S99 | PLEKHF1 | Pleckstrin homology domain-containing family F member 1;PLEKHF1;ortholog |
| HUMAN\|HGNC=9200\|UniProtKB=O00411 | POLRMT | DNA-directed RNA polymerase, mitochondrial;POLRMT;ortholog |
| HUMAN\|HGNC=23531\|UniProtKB=Q5VZY2 | PPAPDC1A | Phospholipid phosphatase 4;PLPP4;ortholog |
| HUMAN\|HGNC=9358\|UniProtKB=P48147 | PREP | Prolyl endopeptidase;PREP;ortholog |
| HUMAN\|HGNC=17348\|UniProtKB=O43395 | PRPF3 | U4/U6 small nuclear ribonucleoprotein Prp3;PRPF3;ortholog |
| HUMAN\|HGNC=16463\|UniProtKB=O75400 | PRPF40A | Pre-mRNA-processing factor 40 homolog A;PRPF40A;ortholog |
| HUMAN\|HGNC=9480\|UniProtKB=Q9NQE7 | PRSS16 | Thymus-specific serine protease;PRSS16;ortholog |
| HUMAN\|HGNC=19096\|UniProtKB=Q8NDX1 | PSD4 | PH and SEC7 domain-containing protein 4;PSD4;ortholog |
| HUMAN\|HGNC=17822\|UniProtKB=Q9H7Z7 | PTGES2 | Prostaglandin E synthase 2;PTGES2;ortholog |
| HUMAN\|HGNC=9650\|UniProtKB=P17706 | PTPN2 | Tyrosine-protein phosphatase non-receptor type 2;PTPN2;ortholog |
| HUMAN\|HGNC=9692\|UniProtKB=P26022 | PTX3 | Pentraxin-related protein PTX3;PTX3;ortholog |
| HUMAN\|HGNC=26505\|UniProtKB=Q3MIT2 | PUS10 | Putative tRNA pseudouridine synthase Pus10;PUS10;ortholog |
| HUMAN\|HGNC=29982\|UniProtKB=P83859 | QRFP | Orexigenic neuropeptide QRFP;QRFP;ortholog |
| HUMAN\|HGNC=9795\|UniProtKB=Q92696 | RABGGTA | Geranylgeranyl transferase type-2 subunit alpha;RABGGTA;ortholog |
| HUMAN\|HGNC=9807\|UniProtKB=O75943 | RAD17 | Cell cycle checkpoint protein RAD17;RAD17;ortholog |
| HUMAN\|HGNC=9832\|UniProtKB=P55895 | RAG2 | V(D)J recombination-activating protein 2;RAG2;ortholog |
| HUMAN\|HGNC=9840\|UniProtKB=P11234 | RALB | Ras-related protein Ral-B;RALB;ortholog |
| HUMAN\|HGNC=15864\|UniProtKB=Q9BYM8 | RBCK1 | RanBP-type and C3HC4-type zinc finger-containing protein 1;RBCK1;ortholog |
| HUMAN\|HGNC=9894\|UniProtKB=Q08999 | RBL2 | Retinoblastoma-like protein 2;RBL2;ortholog |
| HUMAN\|HGNC=16502\|UniProtKB=P58872 | RHBDL3 | Rhomboid-related protein 3;RHBDL3;ortholog |
| HUMAN\|HGNC=21158\|UniProtKB=Q8IUD6 | RNF135 | E3 ubiquitin-protein ligase RNF135;RNF135;ortholog |

| | | |
|---|---|---|
| HUMAN\|HGNC=13432\|UniProtKB=Q9NV58 | RNF19A | E3 ubiquitin-protein ligase RNF19A;RNF19A;ortholog |
| HUMAN\|HGNC=17985\|UniProtKB=Q8WZ75 | ROBO4 | Roundabout homolog 4;ROBO4;ortholog |
| HUMAN\|HGNC=25791\|UniProtKB=Q8IXW5 | RPAP2 | Putative RNA polymerase II subunit B1 CTD phosphatase RPAP2;RPAP2;ortholog |
| HUMAN\|HGNC=30350\|UniProtKB=Q9H9Y2 | RPF1 | Ribosome production factor 1;RPF1;ortholog |
| HUMAN\|HGNC=10302\|UniProtKB=P30050 | RPL12 | 60S ribosomal protein L12;RPL12;ortholog |
| HUMAN\|HGNC=10371\|UniProtKB=P05388 | RPLP0 | 60S acidic ribosomal protein P0;RPLP0;ortholog |
| HUMAN\|HGNC=10402\|UniProtKB=P39019 | RPS19 | 40S ribosomal protein S19;RPS19;ortholog |
| HUMAN\|HGNC=6502\|UniProtKB=P08865 | RPSA | 40S ribosomal protein SA;RPSA;ortholog |
| HUMAN\|HGNC=10450\|UniProtKB=O14718 | RRH | Visual pigment-like receptor peropsin;RRH;ortholog |
| HUMAN\|HGNC=13081\|UniProtKB=Q9BY12 | SCAPER | S phase cyclin A-associated protein in the endoplasmic reticulum;SCAPER;ortholog |
| HUMAN\|HGNC=21088\|UniProtKB=Q86SK9 | SCD5 | Stearoyl-CoA desaturase 5;SCD5;ortholog |
| HUMAN\|HGNC=15950\|UniProtKB=Q9BWW7 | SCRT1 | Transcriptional repressor scratch 1;SCRT1;ortholog |
| HUMAN\|HGNC=8951\|UniProtKB=P07093 | SERPINE2 | Glia-derived nexin;SERPINE2;ortholog |
| HUMAN\|HGNC=30784\|UniProtKB=Q14140 | SERTAD2 | SERTA domain-containing protein 2;SERTAD2;ortholog |
| HUMAN\|HGNC=12950\|UniProtKB=Q15637 | SF1 | Splicing factor 1;SF1;ortholog |
| HUMAN\|HGNC=10809\|UniProtKB=Q13326 | SGCG | Gamma-sarcoglycan;SGCG;ortholog |
| HUMAN\|HGNC=25321\|UniProtKB=Q9H0F6 | SHARPIN | Sharpin;SHARPIN;ortholog |
| HUMAN\|HGNC=11014\|UniProtKB=Q99726 | SLC30A3 | Zinc transporter 3;SLC30A3;ortholog |
| HUMAN\|HGNC=11047\|UniProtKB=Q9UN76 | SLC6A14 | Sodium- and chloride-dependent neutral and basic amino acid transporter B(0+);SLC6A14;ortholog |
| HUMAN\|HGNC=23092\|UniProtKB=Q8TCU3 | SLC7A13 | Solute carrier family 7 member 13;SLC7A13;ortholog |
| HUMAN\|HGNC=14013\|UniProtKB=Q9NTJ3 | SMC4 | Structural maintenance of chromosomes protein 4;SMC4;ortholog |
| HUMAN\|HGNC=20465\|UniProtKB=Q8IY18 | SMC5 | Structural maintenance of chromosomes protein 5;SMC5;ortholog |
| HUMAN\|HGNC=18122\|UniProtKB=Q9H6I2 | SOX17 | Transcription factor SOX-17;SOX17;ortholog |
| HUMAN\|HGNC=32006\|UniProtKB=Q5VVP1 | SPATA31A6 | Spermatogenesis-associated protein 31A6;SPATA31A6;ortholog |
| HUMAN\|HGNC=24508\|UniProtKB=B4DYI2 | SPATA31C2 | Spermatogenesis-associated protein 31C2;SPATA31C2;ortholog |
| HUMAN\|HGNC=24031\|UniProtKB=Q9HBM1 | SPC25 | Kinetochore protein Spc25;SPC25;ortholog |
| HUMAN\|HGNC=11337\|UniProtKB=Q99909 | SSX3 | Protein SSX3;SSX3;ortholog |
| HUMAN\|HGNC=10863\|UniProtKB=Q16842 | ST3GAL2 | CMP-N-acetylneuraminate-beta-galactosamide-alpha-2,3-sialyltransferase 2;ST3GAL2;ortholog |
| HUMAN\|HGNC=23317\|UniProtKB=P61647 | ST8SIA6 | Alpha-2,8-sialyltransferase 8F;ST8SIA6;ortholog |
| HUMAN\|HGNC=11389\|UniProtKB=Q15831 | STK11 | Serine/threonine-protein kinase STK11;STK11;ortholog |
| HUMAN\|HGNC=30796\|UniProtKB=Q9Y3F4 | STRAP | Serine-threonine kinase receptor-associated protein;STRAP;ortholog |
| HUMAN\|HGNC=11427\|UniProtKB=Q9UNE7 | STUB1 | E3 ubiquitin-protein ligase CHIP;STUB1;ortholog |
| HUMAN\|HGNC=19694\|UniProtKB=Q6ZWJ1 | STXBP4 | Syntaxin-binding protein 4;STXBP4;ortholog |
| HUMAN\|HGNC=27411\|UniProtKB=Q6PIF2 | SYCE2 | Synaptonemal complex central element protein 2;SYCE2;ortholog |

| | | |
|---|---|---|
| HUMAN\|HGNC=11523\|UniProtKB=O953 59 | TACC2 | Transforming acidic coiled-coil-containing protein 2;TACC2;ortholog |
| HUMAN\|HGNC=11585\|UniProtKB=O609 07 | TBL1X | F-box-like/WD repeat-containing protein TBL1X;TBL1X;ortholog |
| HUMAN\|HGNC=20854\|UniProtKB=Q9H0 W7 | THAP2 | THAP domain-containing protein 2;THAP2;ortholog |
| HUMAN\|HGNC=11793\|UniProtKB=P528 88 | THOP1 | Thimet oligopeptidase;THOP1;ortholog |
| HUMAN\|HGNC=11803\|UniProtKB=O954 11 | TIAF1 | TGFB1-induced anti-apoptotic factor 1;TIAF1;ortholog |
| HUMAN\|HGNC=14523\|UniProtKB=Q96 MW7 | TIGD1 | Tigger transposable element-derived protein 1;TIGD1;ortholog |
| HUMAN\|HGNC=33522\|UniProtKB=A6NG C4 | TLCD2 | TLC domain-containing protein 2;TLCD2;ortholog |
| HUMAN\|HGNC=24257\|UniProtKB=Q9HC 24 | TMBIM4 | Protein lifeguard 4;TMBIM4;ortholog |
| HUMAN\|HGNC=16996\|UniProtKB=Q153 63 | TMED2 | Transmembrane emp24 domain-containing protein 2;TMED2;ortholog |
| HUMAN\|HGNC=16823\|UniProtKB=P171 52 | TMEM11 | Transmembrane protein 11, mitochondrial;TMEM11;ortholog |
| HUMAN\|HGNC=30366\|UniProtKB=Q925 45 | TMEM131 | Transmembrane protein 131;TMEM131;ortholog |
| HUMAN\|HGNC=31723\|UniProtKB=Q24J Q0 | TMEM241 | Transmembrane protein 241;TMEM241;ortholog |
| HUMAN\|HGNC=32393\|UniProtKB=Q6ZN R0 | TMEM91 | Transmembrane protein 91;TMEM91;ortholog |
| HUMAN\|HGNC=32431\|UniProtKB=Q6UX Z0 | TMIGD1 | Transmembrane and immunoglobulin domain-containing protein 1;TMIGD1;ortholog |
| HUMAN\|HGNC=11905\|UniProtKB=O147 63 | TNFRSF10 B | Tumor necrosis factor receptor superfamily member 10B;TNFRSF10B;ortholog |
| HUMAN\|HGNC=11983\|UniProtKB=O756 74 | TOM1L1 | TOM1-like protein 1;TOM1L1;ortholog |
| HUMAN\|HGNC=12032\|UniProtKB=Q129 33 | TRAF2 | TNF receptor-associated factor 2;TRAF2;ortholog |
| HUMAN\|HGNC=30832\|UniProtKB=Q96Q 05 | TRAPPC9 | Trafficking protein particle complex subunit 9;TRAPPC9;ortholog |
| HUMAN\|HGNC=31454\|UniProtKB=Q6A5 55 | TXNDC8 | Thioredoxin domain-containing protein 8;TXNDC8;ortholog |
| HUMAN\|HGNC=12477\|UniProtKB=P519 65 | UBE2E1 | Ubiquitin-conjugating enzyme E2 E1;UBE2E1;ortholog |
| HUMAN\|HGNC=12492\|UniProtKB=P610 88 | UBE2N | Ubiquitin-conjugating enzyme E2 N;UBE2N;ortholog |
| HUMAN\|HGNC=15664\|UniProtKB=Q9NY U1 | UGGT2 | UDP-glucose:glycoprotein glucosyltransferase 2;UGGT2;ortholog |
| HUMAN\|HGNC=20332\|UniProtKB=Q9H1 J1 | UPF3A | Regulator of nonsense transcripts 3A;UPF3A;ortholog |
| HUMAN\|HGNC=20485\|UniProtKB=O753 17 | USP12 | Ubiquitin carboxyl-terminal hydrolase 12;USP12;ortholog |
| HUMAN\|HGNC=14340\|UniProtKB=Q9UB Q0 | VPS29 | Vacuolar protein sorting-associated protein 29;VPS29;ortholog |
| HUMAN\|HGNC=25072\|UniProtKB=Q5M NZ6 | WDR45B | WD repeat domain phosphoinositide-interacting protein 3;WDR45B;ortholog |
| HUMAN\|HGNC=14540\|UniProtKB=Q9H4 A3 | WNK1 | Serine/threonine-protein kinase WNK1;WNK1;ortholog |
| HUMAN\|HGNC=12786\|UniProtKB=O007 55 | WNT7A | Protein Wnt-7a;WNT7A;ortholog |
| HUMAN\|HGNC=28304\|UniProtKB=Q96E C8 | YIPF6 | Protein YIPF6;YIPF6;ortholog |
| HUMAN\|HGNC=31675\|UniProtKB=Q9Y5 A9 | YTHDF2 | YTH domain-containing family protein 2;YTHDF2;ortholog |
| HUMAN\|HGNC=17908\|UniProtKB=O432 98 | ZBTB43 | Zinc finger and BTB domain-containing protein 43;ZBTB43;ortholog |
| HUMAN\|HGNC=29362\|UniProtKB=Q9C0 D7 | ZC3H12C | Probable ribonuclease ZC3H12C;ZC3H12C;ortholog |
| HUMAN\|HGNC=20368\|UniProtKB=Q5T2 00 | ZC3H13 | Zinc finger CCCH domain-containing protein 13;ZC3H13;ortholog |
| HUMAN\|HGNC=14881\|UniProtKB=O603 15 | ZEB2 | Zinc finger E-box-binding homeobox 2;ZEB2;ortholog |

| | | |
|---|---|---|
| HUMAN\|HGNC=12863\|UniProtKB=Q9Y6Q3 | ZFP37 | Zinc finger protein 37 homolog;ZFP37;ortholog |
| HUMAN\|HGNC=20997\|UniProtKB=Q9H091 | ZMYND15 | Zinc finger MYND domain-containing protein 15;ZMYND15;ortholog |
| HUMAN\|HGNC=12907\|UniProtKB=Q15973 | ZNF124 | Zinc finger protein 124;ZNF124;ortholog |
| HUMAN\|HGNC=12920\|UniProtKB=P52737 | ZNF136 | Zinc finger protein 136;ZNF136;ortholog |
| HUMAN\|HGNC=23708\|UniProtKB=Q6ZR52 | ZNF493 | Zinc finger protein 493;ZNF493;ortholog |
| HUMAN\|HGNC=13168\|UniProtKB=Q03936 | ZNF92 | Zinc finger protein 92;ZNF92;ortholog |
| HUMAN\|HGNC=25820\|UniProtKB=Q9C0D3 | ZYG11B | Protein zyg-11 homolog B;ZYG11B;ortholog |

#### Table 4: Preliminary Identification of tumor targets for combination immunotherapy

| Druggable Gene Category | Matching Gene Count | Matching Gene(s) |
|---|---|---|
| DRUGGABLE GENOME | 68 | ABHD16A, ARSF, BAP1, BCL2, BIRC2, CAT, CCNA2, CFLAR, CWC27, CYB5R3, DNAJC16, DNPEP, DPEP2, DPT, EFNB1, EPO, ESRRA, FAM3B, FAT1, FBLN1, GRIK1, HCRTR1, HNRNPUL2, IKBKG, IL1R1, IL36RN, IL9, ITGAV, ANOS1, KCNF1, KCNG2, KLHL5, KLK11, LARS2, LRRTM3, MAN1B1, MAS1L, MID1, MRGPRX4, NEK5, NMS, OR2L2, OR2M2, PIK3C2A, PREP, PRSS16, PTPN2, PTX3, QRFP, RABGGTA, RBCK1, RHBDL3, RNF135, RRH, SCD5, SERPINE2, SHARPIN, SLC6A14, SLC7A13, ST3GAL2, STK11, THOP1, TNFRSF10B, TXNDC8, UBE2N, USP12, WNK1, WNT7A |
| KINASE | 22 | CCNA2, CLP1, DLG3, EPO, FBXW7, GMFG, HMGXB3, IKBKG, LRRTM3, NEK5, NLRC5, PIK3C2A, PTPN2, RALB, RBL2, STK11, TNFRSF10B, TOM1L1, TRAF2, UBE2N, WNK1, ZEB2 |
| TUMOR SUPPRESSOR | 18 | BAP1, BCL2, BIRC2, BOP1, CCNA2, CUL9, EIF4G1, EIF4G2, FBXW7, GIGYF2, PRPF40A, RAD17, RBL2, SMC5, STK11, STXBP4, TOM1L1, UBE2E1 |
| PROTEASE | 17 | BAP1, BIRC2, CFLAR, DNPEP, DPEP2, FBLN1, ANOS1, KLK11, NEK5, PREP, PRSS16, RHBDL3, SERPINE2, THOP1, TNFRSF10B, TRAF2, USP12 |
| TRANSPORTER | 15 | ATP6V0C, BCL2, DLG3, EPO, GRIK1, ITGAV, KCNF1, KCNG2, SLC30A3, SLC6A14, SLC7A13, TMEM241, UPF3A, VPS29, WNK1 |
| SERINE THREONINE KINASE | 10 | CCNA2, IKBKG, NEK5, RALB, STK11, TNFRSF10B, TRAF2, UBE2N, WNK1, ZEB2 |
| HISTONE MODIFICATION | 9 | BAP1, CCNA2, ENY2, EYA4, MTA2, PAX7, TBL1X, UBE2E1, UBE2N |
| DNA REPAIR | 8 | CEBPG, CINP, EYA4, NPAS2, RAD17, SMC5, STUB1, UBE2N |
| G PROTEIN COUPLED RECEPTOR | 7 | CRCP, HCRTR1, MAS1L, MRGPRX4, OR2L2, OR2M2, RRH |
| ION CHANNEL | 7 | DLG3, EPO, GRIK1, ITGAV, KCNF1, KCNG2, WNK1 |
| TRANSCRIPTION FACTOR BINDING | 7 | BCL2, CEBPG, EGR2, MTA2, RNF19A, SOX17, TBL1X |
| CELL SURFACE | 6 | EPO, IL1R1, ITGAV, SERPINE2, TNFRSF10B, WNT7A |
| CLINICALLY ACTIONABLE | 6 | BAP1, BCL2, FAT1, FBXW7, SOX17, STK11 |
| TYROSINE KINASE | 5 | FBXW7, GMFG, LRRTM3, PTPN2, WNK1 |
| TRANSCRIPTION FACTOR COMPLEX | 4 | MTA2, NPAS2, RBL2, SOX17 |

| | | |
|---|---|---|
| B30_2 SPRY DOMAIN | 3 | HNRNPUL2, MID1, RNF135 |
| DRUG RESISTANCE | 3 | BCL2, CAT, HADH |
| NEUTRAL ZINC METALLOPEPTIDASE | 3 | DNPEP, DPEP2, THOP1 |
| PROTEASE INHIBITOR | 3 | BIRC2, ANOS1, SERPINE2 |
| PROTEIN PHOSPHATASE | 3 | EYA4, PTPN2, RPAP2 |
| EXTERNAL SIDE OF PLASMA MEMBRANE | 2 | ITGAV, SERPINE2 |
| GROWTH FACTOR | 2 | GMFG, IL9 |
| HORMONE ACTIVITY | 2 | EPO, QRFP |
| NUCLEAR HORMONE RECEPTOR | 2 | ESRRA, SF1 |
| THIOREDOXIN | 2 | DNAJC16, TXNDC8 |
| ABC TRANSPORTER | 1 | ATP6V0C |
| LIPID KINASE | 1 | RBL2 |
| PHOSPHATIDYLINOSITOL 3 KINASE | 1 | PIK3C2A |

#### Table 5: Inhibitors utilized in study

| Inhibitor | Alternative nomenclature | Target | Source |
|---|---|---|---|
| Venetoclax | ABT-199 | BCL2 | Selleckchem |
| Navitoclax | ABT-263 | BCL2 | Selleckchem |
| LCL161 | | BIRC2 | Selleckchem |
| Birinapant | | BIRC2 | Selleckchem |
| Fomepizole | | CAT | Selleckchem |
| Tosedostat | CHR2797 | DNPEP | Tocris |
| XCT790 | | ESRRA | Sigma Aldrich |
| Compound 29 | | ESRRA | Synthesized by collaborator |
| Topiramate | | GRIK1 | Selleckchem |
| ACET | | GRIK1 | Tocris |
| Almorexant | ACT-078573 | HCRTR1 | Selleckchem |
| SB334867 | | HCRTR1 | R&D systems |
| Blocking antibody | AF269 | IL1R1 | R&D systems |
| Blocking antibody | AF209 | IL9 | R&D systems |
| Blocking antibody | ab16821 | ITGAV | Abcam |
| Cilengitide | | ITGAV | Selleckchem |
| Guanidine HCL | | KCNF1 | Sigma Aldrich |
| Dalfampridine | 4-aminopyridine | KCNF1 | Sigma Aldrich |
| Guanidine HCL | | KCNG2 | Sigma Aldrich |
| Dalfampridine | 4-aminopyridine | KCNG2 | Sigma Aldrich |
| Dactolisib | BEZ235 | PIK3C2A | Selleckchem |

| | | | |
|---|---|---|---|
| Apitolisib | GDC-0980 | PIK3C2A | Selleckchem |
| S17092 | | PREP | Sigma Aldrich |
| WNK463 | | WNK1 | Selleckchem |

#### Table 6: sgRNA sequences

| sgRNA Name | sgRNA sequence |
|---|---|
| RPAP2 sgRNA1 | GGCCCAGCGAAGTCCGCCAT |
| RPAP2 sgRNA2 | AAATTCTCGTAACTTGGCAG |
| RPAP2 sgRNA3 | CGCTGCTCTCGAAAAGCCGC |
| BIRC2 sgRNA1 | TGGAGACGTATTCTTAGAGG |
| BIRC2 sgRNA2 | ACATATTCAACTTTCCCCGC |
| BIRC2 sgRNA3 | ATATTCAACTTTCCCCGCCG |
| ESD sgRNA1 | CGTTGTTTTTCGATTGCAAG |
| ESD sgRNA2 | TAGACCACAAACACTTACGA |
| ESD sgRNA3 | CACTTACGAGGGCTGGTATC |
| EGR2 sgRNA1 | GCAAGACGCCGGTGCACGAG |
| EGR2 sgRNA2 | TCAAGGTGTCCGGGTCCGAG |
| EGR2 sgRNA3 | CTCGTGCACCGGCGTCTTGC |
| SCAF11 sgRNA1 | TGTCTTCGAGCTATCTGCGC |
| SCAF11 sgRNA2 | TCTGGTTGGGTATCTAACCG |
| SCAF11 sgRNA3 | CTGTACCGATCATTTCCTCG |
| TOR1AIP1 sgRNA1 | CATAAACTTACGTCGGCTGG |
| TOR1AIP1 sgRNA2 | TCAACAACTATGGCGGGCGA |
| TOR1AIP1 sgRNA3 | GCGCGTACTACCTTCGGTCT |
| LPGAT1 sgRNA1 | AAGCGGTTCTGGTATATCGA |
| LPGAT1 sgRNA2 | CTTCATGGTCGTCAACAACC |
| LPGAT1 sgRNA3 | CTTACTGTCCAGCACTCGAA |
| SERTAD2 sgRNA1 | CCAGGGGCGTAGTGCTTCCG |
| SERTAD2 sgRNA2 | CGTAGTGCTTCCGAGGTCGC |
| SERTAD2 sgRNA3 | GTAGTGCTTCCGAGGTCGCA |
| RBL2 sgRNA1 | CCGCCTCAACATGGACGAGG |
| RBL2 sgRNA2 | CGACGGCATAGCGCACCCCT |
| RBL2 sgRNA3 | CTATGCGTAGCTGTAGAAAC |
| BCL2 sgRNA1 | AAGCGTCCCCGCGCGGTGAA |
| BCL2 sgRNA2 | ACCTGACGCCCTTCACCGCG |
| BCL2 sgRNA3 | GGGGCCGTACAGTTCCACAA |
